# Comprehensive analysis across mammalian tissues and cells decipher the underlying mechanism of m6A specificity

**DOI:** 10.1101/2024.04.19.590363

**Authors:** Guo-Shi Chai, Hong-Xuan Chen, Dong-Zhao Ma, Ze-Hui Ren, Xue-Hong Liu, Guan-Zheng Luo

**Affiliations:** State Key Laboratory of Biocontrol, MOE Key Laboratory of Gene Function and Regulation, Guangdong Province Key Laboratory of Pharmaceutical Functional Genes, School of Life Sciences, Sun Yat-sen University, Guangzhou 510275, China; School of Life Sciences, Westlake University, Hangzhou, Zhejiang, China

## Abstract

N6-methyladenine (m6A) stands out as the most prevalent internal chemical modification on mammalian mRNA, playing a vital role in diverse biological processes. Despite considerable advancements in individual cell line studies, the characteristics of m6A sites across distinct cell lines or tissues remain elusive. In this study, we have successfully identified approximately 1.5 million high-confidence m6A sites in human and mouse cell lines or tissues using published m6A-seq data. By categorizing m6A sites into different consistency levels, we observe that those of high consistency are notably enriched near the stop codon. Furthermore, they exhibit a higher likelihood of interaction with known m6A binding proteins such as YTHDF1-3, RBM15, YTHDC1, and IGF2BP1, thereby influencing gene expression homeostasis. Additionally, these sites display a higher CpG density in the promoter region of the genes they mark, with METTL3 demonstrating a preference for binding to the promoter region of its marked genes. m6A sites of low consistency levels, including unique m6A sites, show a significant enrichment near the start codon. These sites are more prone to binding by newly discovered m6A-binding proteins such as DDX3X, PRPF8, and EIF3G. The identification of these distinct features of m6A sites lays a foundational understanding for unraveling the functional roles of m6A.

## Introduction

More than 150 nucleic acid modifications have been characterized so far^1,2^. They are found in all types of RNA molecules, including ribosomal RNA (rRNA), transfer RNA (tRNA), small nuclear RNA (snRNA), messenger RNA (mRNA), and long non-coding RNA (lncRNA). Various modifications have been described in eukaryotic mRNA, including N6-methyladenosine (m6A), 5-methylcytidine (m5C), inosine (I), pseudouridine (Ψ), and N1-methyladenosine (m1A). Their presence significantly impacts mRNA metabolism and function. Notably, m6A, the most prevalent internal modification in eukaryotic mRNA, has emerged as a focal point of investigation. As a dynamic and reversible chemical modification, m6A involves writers catalyzing methyl group addition, erasers responsible for methyl group removal, and m6A-binding proteins, also known as readers, selectively bind to the modified RNA^3^. These reader proteins play a crucial role in regulating gene expression in an m6A-dependent manner. Experimental endeavors employing in-vitro m6A RNA probe pull-down and protein mass spectrometry in human or mouse cell lines have unveiled numerous RNA-binding proteins (RBPs))^4–6^. Among them, RBPs directly engaging with m6A feature YTH domains, exemplified by YTHDF1, YTHDF2, YTHDF3, YTHDC1, and YTHDC2. Renowned as critical m6A readers, these proteins significantly influence mRNA stability^7^, translation^8^, alternative polyadenylation, splicing^9,10^, and nuclear export^11^. IGF2BP1, IGF2BP2, and IGF2BP3 contribute to mRNA stability, impeding mRNA degradation by binding directly or indirectly to m6A-labeled mRNA^6^.

To comprehensively explore m6A modifications across the transcriptome, various mapping methods have been developed, including miCLIP^12^, m6A-Seq2^13^, DART-seq^14^, m6A-label-seq^15^, MAZTER-seq^16^, m6A-REF-seq^17^, GLORI^18^, eTAM-seq^19^, m6Anet^20^, m6ABasecaller^21^ and SingleMod^22^. Among these, m6A-seq/MeRIP-seq^4,23^ remains the most widely used method, which uses commercial m6A antibodies to enrich m6A-labeled RNA fragments for high-throughput sequencing. While m6A-seq provides valuable information on m6A-marked genes and the position of m6A, it has an average resolution of about 200 nt. Furthermore, this technique has inherent limitations, including a high rate of rRNA contamination, low library complexity, and variable m6A antibody enrichment efficiency, leading to the detection of false positives and low-confidence m6A sites^24,25^. Therefore, the evaluation of m6A-seq datasets becomes crucial for identifying high-confidence m6A sites.

Previous investigations have indicated that m6As regulate gene expression homeostasis across various human fetal tissues, with enrichment in genes exhibiting CpG-rich promoters^26^. However, the interplay between m6A sites of differing consistency levels, m6A-binding proteins, gene expression homeostasis, and their association with CpG density in gene promoter regions remains elusive across diverse tissues and species. In this study, we carefully assessed the quality of 193 m6A-seq samples, identifying approximately 1.5 million high-confidence m6A sites. These sites were categorized into different consistency levels based on the number of their occurrences in cell lines or tissues. Our findings reveal a significant enrichment of m6A sites with high consistency levels near stop codons. Known m6A binding proteins exhibit a preference for binding to m6A sites of high consistency levels. Moreover, the consistency level of m6A sites correlates positively with gene expression homeostasis and CpG density in gene promoter regions. METTL3 displays a preference for binding to the promoter regions of genes marked by m6As of high consistency levels. Conversely, m6A sites of low consistency levels are notably enriched near start codons. Newly identified m6A-binding proteins prefer to bind to low-consistency m6A sites. Notably, the observed features of m6A sites of different consistency levels are conserved across human tissues and cell lines, as well as mouse tissues and cell lines. This conservation underscores the role of m6A in deposition and the regulation of gene expression, offering novel insights into the diverse functions of m6A.

## Results

### Identification of high-confidence m6A sites

To build comprehensive high-confidence m6A datasets, we collected 193 sets of m6A-seq (IP) and corresponding RNA-seq (Input) samples from public databases, covering 36 different research groups, 20 mouse tissues and cell lines, and 34 human tissues and cell lines (Figure 1A, Supplementary Table 1-4). These samples are all in the wild type or control conditions, including at least two biological replicates. To ensure the reliability of further analysis, we setup a series of metrics to evaluate the quality of these datasets. rRNA contamination is a common issue in RNA-seq based experiments. We first checked the rRNA contamination rate of all samples (Supplementary Figure 1A, B). The median rRNA contamination rate in IP samples was 1%, and that in Input samples was 3.3%. The rRNA contamination rate in IP samples was significantly lower than in Input samples, consistent with previous reports^27^. All the IP samples did not have a very high rRNA contamination rate, indicating that IP samples did not suffer from rRNA contamination. Based on the performance of 75% of IP samples, we defined an empirical value for the rRNA contamination rate in IP samples as <5%.

**Figure 1.**
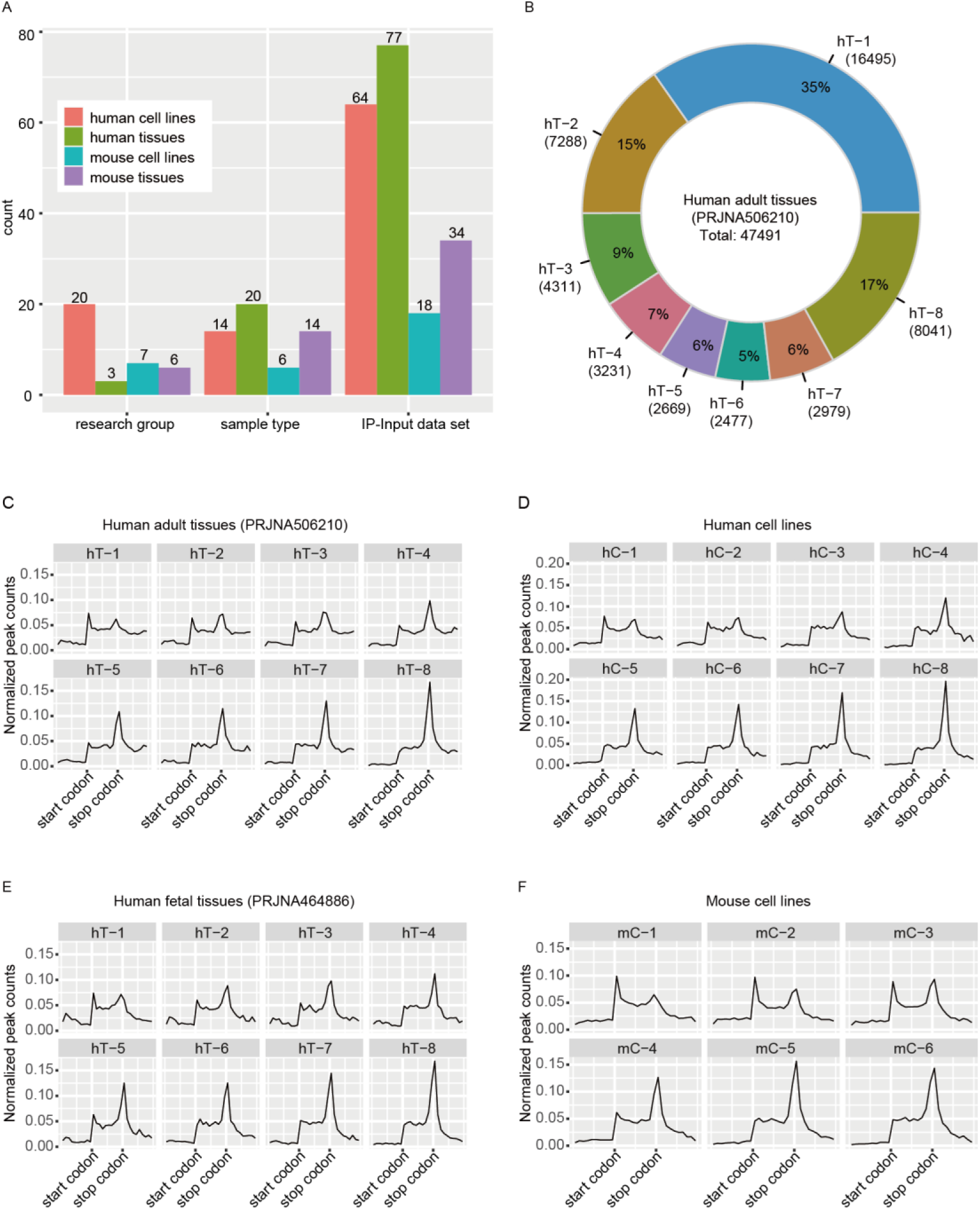
Sample information in this study, and the proportion and distribution patterns of m6A sites of different consistency levels. A, The number of research groups, sample types, and IP-Input datasets in human cell lines, human tissues, mouse cell lines, and mouse tissues. B, m6A sites detected in eight different adult tissues are classified into eight consistency levels, and the number and proportion of m6A sites of eight consistency levels are shown. C, Distribution patterns of m6A sites of eight consistency levels defined by eight different adult tissues along the transcriptome. D, Distribution patterns of m6A sites of eight consistency levels defined by eight different human cell lines along the transcriptome. E, Distribution patterns of m6A sites of eight consistency levels defined by eight different human fetal tissues along the transcriptome. F, Distribution patterns of m6A sites of six consistency levels defined by six different mouse cell lines along the transcriptome. hT denotes human tissue. hC denotes human cell. hT-1 denotes that m6A is present in one type of human tissue. hT-8 denotes that m6A is present in eight different human tissues simultaneously. mT denotes mouse tissue. mC denotes mouse cell. mC-1 denotes that m6A is present in one type of mouse cell line. mC-6 denotes that m6A is present in six different mouse cell lines simultaneously.

Secondly, we interrogated the library complexity of all samples using three metrics defined by the ENCODE project: NRF (Non-Redundant Fraction), PBC1 (PCR Bottlenecking Coefficient 1), and PBC2 (PCR Bottlenecking Coefficient 2)^28^. The median values of NRF, PBC1, and PBC2 in IP samples are 0.16, 0.41, and 2.47, while in Input samples, they are 0.24, 0.59, and 2.91, respectively. The higher library complexity of Input samples compared to IP samples is expected, given that IP sample undergo a sequence sampling process from Input sample (Supplementary Figure 1C-H). According to the ChIP-seq library quality reference values defined by ENCODE^28^, only 4 IP samples performed within the acceptable range for the NRF reference value; 73 IP samples exhibited a moderate PCR bottleneck level or higher based on the PBC1 reference value; and 193 IP samples showed a moderate PCR bottleneck level or higher according to the PBC2 reference value. These results indicate that using the ChIP-seq library quality reference values to evaluate m6A-seq IP samples is inappropriate. Similar to the rRNA contamination rate, based on the performance of 75% of IP samples, we defined the empirical values of NRF, PBC1, and PBC2 as >8%, >0.2, and >2, respectively.

Thirdly, we used FRiP (Fraction of reads in peaks) to evaluate the enrichment efficiency of IP procedure (Supplementary Figure 1I, J). Since the FRiP values were very sensitive to the parameters during peak calling, we adopted consistent criteria across different samples. As expected, the FRiP values in IP samples are significantly higher compared to Input samples that the median FRiP value in IP samples is 59.6%, while the median in Input samples is 8%. Similar to other metrics, based on the performance of 75% of IP samples, we defined the empirical value of FRiP in the IP sample as >45%.

Using these aforementioned metrics, we curated collected m6A-seq datasets, comprising 20 human tissues, 8 human cell lines, 11 mouse tissues, and 6 mouse cell lines. We identified approximately 1.5 million m6A sites with high-quality, with an average of roughly 30,000 m6A sites per human sample (including tissues and cell lines) and about 20,000 m6A sites per mouse sample (Supplementary Figure 2). To verify the authenticity of these sites, we first examined the distribution pattern of m6A sites on the transcriptome (Supplementary Figures 3, 5A, and 6A). The results showed that m6A sites were mainly located in the CDS and 3’-UTR regions and significantly enriched near stop codon. Next, we examined the sequences preference where m6A located (Supplementary Figure 4, 5B, and 6B), revealing that the consensus motif sequence DRACH was significantly enriched. These results are consistent with previously reported classic distribution pattern of m6A^4,23^. Remarkably, m6A sites in mouse cell lines 3T3-L1 and MEF were also significantly enriched near start codon, a distribution pattern reported by many studies^29^. Based on the enriched DRACH motif in 3T3-L1 and MEF cell lines, we inferred that the m6A sites enriched near start codon are also authentic . In summary, the detected m6A sites were deemed authentic and credible, forming the basis for downstream analysis.

### Exploring m6A site consistency and distribution

To further characterize m6A features, we categorized m6A sites into different consistency levels based on the number of samples in which each m6A site occurs. A high proportion of m6A sites appeared in only one sample, while a similarly substantial proportion was detected across all samples (Figure 1B, Supplementary Figure 7). We examined the distribution patterns of m6A sites with varying consistency levels on the transcriptome (Figure 1C-F, Supplementary Figure 8), revealing that the low-consistency m6A sites were bimodally distributed near both the start and stop codons. As the consistency level increases, the enrichment of m6A sites at start codons gradually diminished, while the enrichment at stop codons gradually increases, a pattern consistent across humans and mice. The m6A sites with the highest consistency (such as m6A sites in hT-8, hT-14, hC-8, mT-11, and mC-6) were very stable, representing “hard-coded” modifications predominantly located in CDS and 3’-UTR regions, with a marked preference for stop codons. Conversely, the m6A sites with the lowest consistency, or sites that uniquely occurs in one cell or tissue (such as m6A sites in hT-1, hC-1, mT-1, and mC-1), exhibited the most significant preference for start codons.

### m6A sites with different consistency levels predict novel m6A-binding proteins

To search for potential new RNA binding proteins (RBPs) exhibiting preferential binding to m6A sites with varying consistency levels, we analyzed nearly 34 million binding sites of 171 RBPs^30^. We used two types of m6A sites (the most consistent m6A sites and unique m6A sites) to determine potential m6A-binding proteins. In 8 adult tissues (PRJNA506210), 8 RBPs (YTHDF1-3, RBM15, RBM15B, CPSF7, IGF2BP1, and YTHDC1) preferentially bound to the most consistent m6A sites and 20 RBPs preferentially bound to unique m6A sites were identified by unique m6A sites (Figure 2A). The most consistent m6A sites were used to identify 3 RBPs (YTHDF1-3) that preferentially bind to the most consistent m6A sites and 21 RBPs that preferentially bind to unique m6A sites (Figure 2B). The overlapping ratio of RBPs recognized by the two types of m6A sites was very high, indicating that the results had good reproducibility (Figure 2C, D). We further examined the preference levels of the 38 RBPs for m6A sites with varying consistency levels (hT-1 to hT-8 m6A sites). Results showed that 8 RBPs preferred consistent m6A sites, while 30 RBPs preferred unique m6A sites, and these RBPs were potential m6A-binding proteins (Figure 2E).

**Figure 2.**
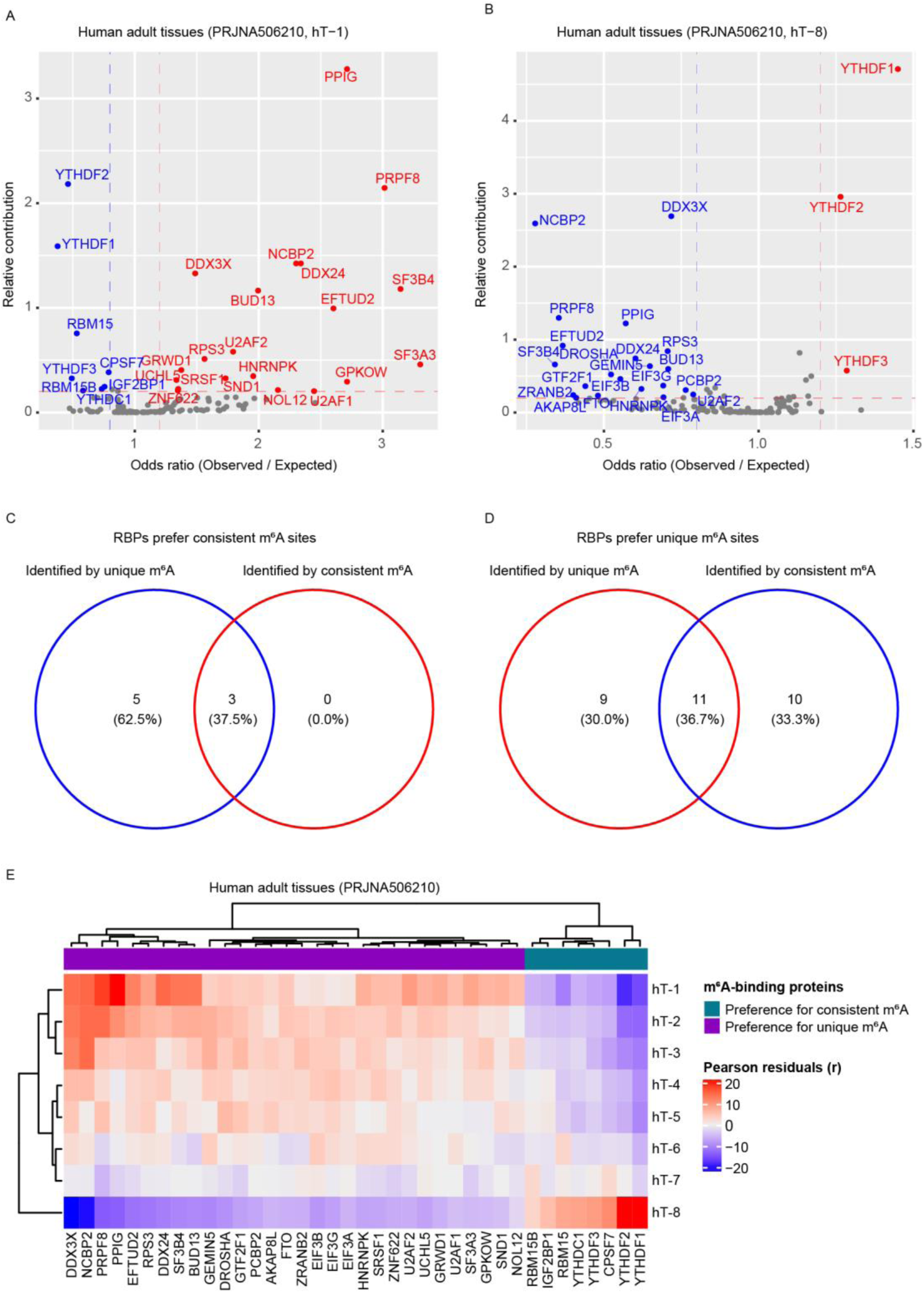
m6A-binding proteins identified by m6A sites of eight consistency levels in eight adult tissues. A, Eight m6A-binding proteins that prefer binding to consistent m6A sites and twenty m6A-binding proteins that prefer binding to unique m6A sites are identified through unique m6A sites enrichment analysis. B, Three m6A-binding proteins that prefer binding to consistent m6A sites and twenty-one m6A-binding proteins that prefer binding to unique m6A sites are identified through the most consistent m6A sites enrichment analysis. C, Overlap analysis of m6A-binding proteins that prefer binding to consistent m6A sites. D, Overlap analysis of m6A-binding proteins that prefer binding to unique m6A sites. E, The affinity of thirty-eight m6A-binding proteins to m6A sites of eight consistency levels in eight adult tissues.

Using the same strategy, in 8 human fetal tissues, 6 RBPs (YTHDF1-3, RBM15, YTHDC1, and IGF2BP1) preferred the consistent m6A sites, and 26 RBPs preferred unique m6A sites (Supplementary Figure 9). In 14 adult tissues (PRJCA001180), 4 RBPs (YTHDF1, YTHDF2, RBM15, and YTHDC1) preferred the consistent m6A sites, and 16 RBPs preferred the unique m6A sites (Supplementary Figure 10). In 8 human cell lines, 6 RBPs (YTHDF1-3, RBM15, CPSF7, and CPSF6) preferred to bind to the consistent m6A sites, and 22 RBPs preferred to bind to unique m6A sites (Supplementary Figure 11). These results showed that RBPs had different binding abilities to m6A sites of varying consistency levels. Known m6A readers such as YTHDF1 and YTHDF2 preferred the high-consistency m6A sites, while the newly identified RBPs preferred to bind to unique m6A sites.

## Novel m6A-binding proteins can interact with known m6A regulators

We performed an overlapping analysis of potential m6A-binding proteins identified in human cell lines and three types of human tissues (two types of adult human tissues and a type of fetal tissue) (Figure 3A, B). A total of 9 RBPs were identified that preferentially bound to the consistent m6A sites, of which 3 RBPs (YTHDF1, YTHDF2, and RBM15) were screened out simultaneously in human cell lines and three types of human tissues. RBM15B was screened out in adult tissues (PRJNA506210), and CPSF6 was identified in human cell lines. A total of 40 RBPs were identified that preferentially bound to unique m6A sites, of which 9 RBPs (BUD13, DDX3X, DROSHA, EFTUD2, GEMIN5, GTF2F1, NCBP2, PPIG, and PRPF8) were found in human cells lines and three types of human tissues. Six RBPs (GPKOW, NOL12, SND1, U2AF1, UCHL5, and EIF3A) were screened out in adult tissues (PRJNA506210). Four RBPs (EIF3D, FKBP4, DDX59, and XRCC6) were screened out in fetal tissues, and three RBPs (SERBP1, GRSF1, and TRA2A) were screened out in human cell lines.

**Figure 3.**
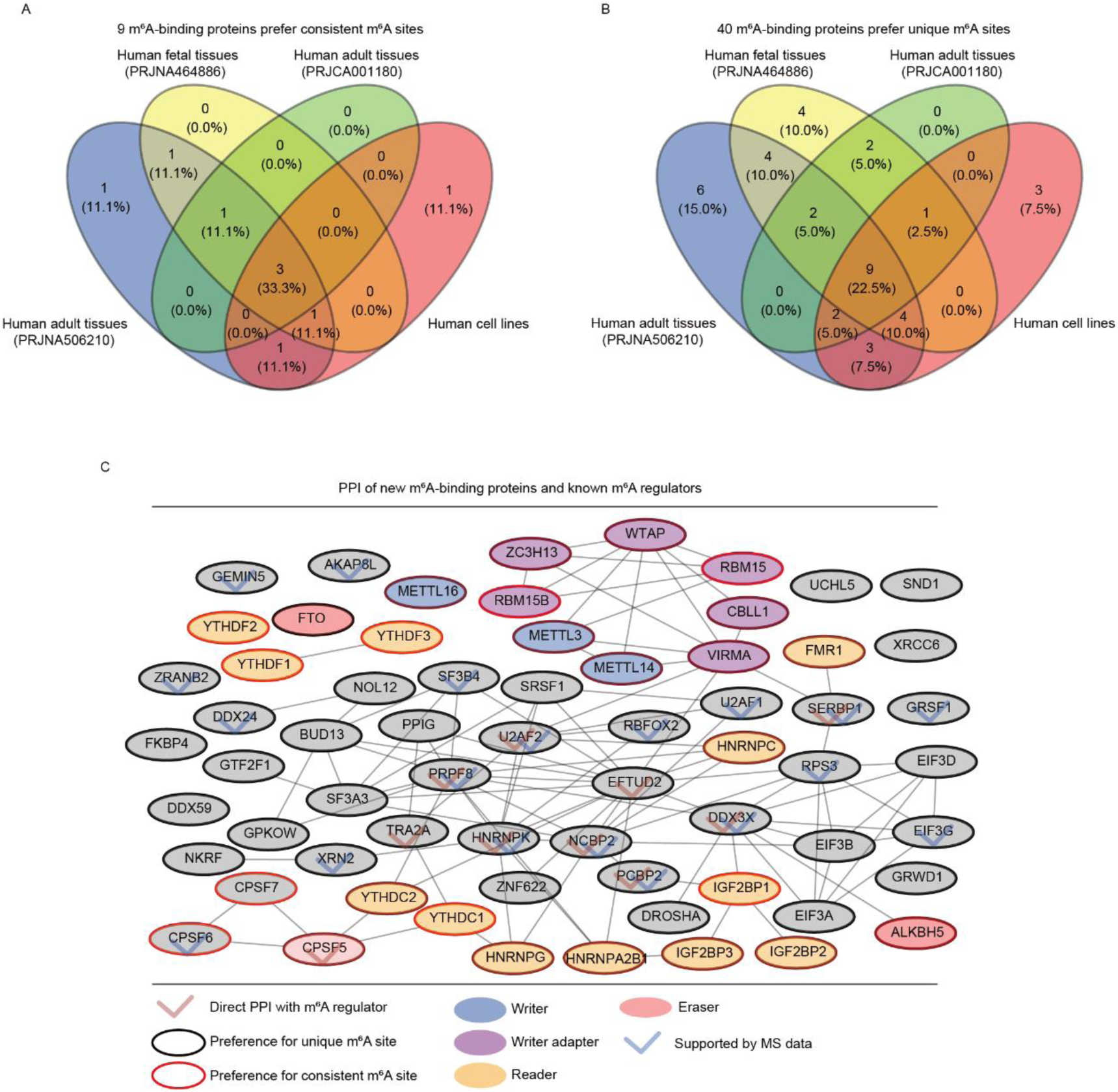
Overlap analysis and protein-protein interaction network of m6A-binding proteins identified in human cell lines and three types of human tissues. A, Overlap analysis of nine m6A-binding proteins that prefer binding to consistent m6A sites. B, Overlap analysis of forty m6A-binding proteins that prefer binding to unique m6A sites. C, Experimentally determined protein-protein interactions between m6A-binding proteins (gray) and known m6A regulators (colors). PPI denotes protein-protein interaction. MS denotes mass spectrometry.

We constructed the protein-protein interaction (PPI) networks to study the interactions between the potential m6A-binding proteins and known m6A regulators (writers, readers, and erasers) (Figure 3C). RNA helicase DDX3X, preferentially bound to the unique m6A site, interacted with the m6A demethylase ALKBH5 and regulated the mRNA demethylation process^29,31^. Nine RBPs (U2AF2, EFTUD2, SERBP1, CPSF5/NUDT21, PRPF8, TRA2A, HNRNPK, NCBP2, and PCBP2) that preferentially bound to unique m6A sites, interacted with m6A writers or readers, and these proteins may function as "adapter" proteins. Two m6A-binding proteins that preferentially bound to the high-consistency m6A sites, CPSF6 and CPSF7, were cleavage and polyadenylation factors that participated in the cleavage and polyadenylation of the 3’ end of pre-mRNA by forming a complex with CPSF5. During the polyadenylation process, this complex can interact with the nuclear reader proteins YTHDC1 and YTHDC2, indicating that YTHDC1 and YTHDC2 were involved in the polyadenylation process at the 3’ end of pre-mRNA. Five m6A-binding proteins (PRPF8, SRSF1, SF3B4, U2AF2, and SF3A3) that preferred unique m6A sites were splicing factors. These proteins formed a complex through interaction. The PRPF8 splicing factor can interact with YTHDC2, indicating that YTHDC2 regulated mRNA splicing. The four m6A-binding proteins (EIF3A, EIF3B, EIF3D, and EIF3G) were translation initiation factors that preferred to bind to unique m6A sites. They can form a translation initiation complex through interaction. Among them, EIF3B interacted with the RNA helicase DDX3X, indicating that unique m6A sites can regulate the initiation of mRNA translation. The demethylase FTO, preferentially bound to unique m6A sites, did not interact with known m6A regulators and newly identified m6A-binding proteins. A total of 19 new m6A binding proteins that preferred to bind to unique m6A sites appeared in in-vitro m6A RNA probe pull-down and protein mass spectrometry experimental results^5,32,33^, indicating that these potential m6A-binding proteins bound to m6A sites blocked the demethylation process ultimately affecting m6A levels at these sites^34^.

### Mapping m6A modification and corresponding gene expression landscape

To study the relationship between gene expression and m6A modification, we first constructed the m6A modification and corresponding gene expression map. Here, we defined m6A present in multiple tissues or cell lines as mRNA or lncRNA-consistent m6A sites, otherwise mRNA or lncRNA-unique m6A sites (view methods for details). Specifically, in the eight adult tissues (PRJNA506210), we identified 30,996 mRNA-consistent m6A sites and 16,495 mRNA-unique m6A sites, accounting for ∼65% and ∼35%, respectively (Figure 4A, B). For lncRNA, 3203 consistent m6A sites and 3306 unique m6A sites were identified, accounting for ∼49% and ∼51%, respectively (Figure 4C, D). In a single tissue, mRNA-consistent m6A sites ranged from ∼16,000 to ∼21,000, with the most significant number of sites detected in the frontal cortex at 21,384 (Supplementary Figure 17A). mRNA-unique m6A sites ranged from ∼1300 to ∼3600, and the number of sites detected in the frontal cortex ranks second at 2332 (Supplementary Figure 18A). lncRNA-consistent m6A sites ranged from ∼1000 to ∼ 2200, and the frontal cortex had the highest number of 2246 (Supplementary Figure 19A). lncRNA-unique m6A sites ranged from 175 to 1000, with the highest number of sites detected in the frontal cortex at 1000 (Supplementary Figure 20A). Compared with the other seven tissues, the frontal cortex had the most m6A sites, indicating that m6A played an essential regulatory role in the frontal cortex. If the mRNA-consistent m6A sites appeared in more tissues, the expression level of its marked genes was more stable among tissues (Figure 4A). This trend was also actual for lncRNA-consistent m6A sites (Figure 4C). Genes marked by the mRNA-unique m6A sites had no specific expression trend, while genes characterized by the lncRNA-unique m6A sites had a specific expression trend among tissues (Figure 4B, D).

**Figure 4.**
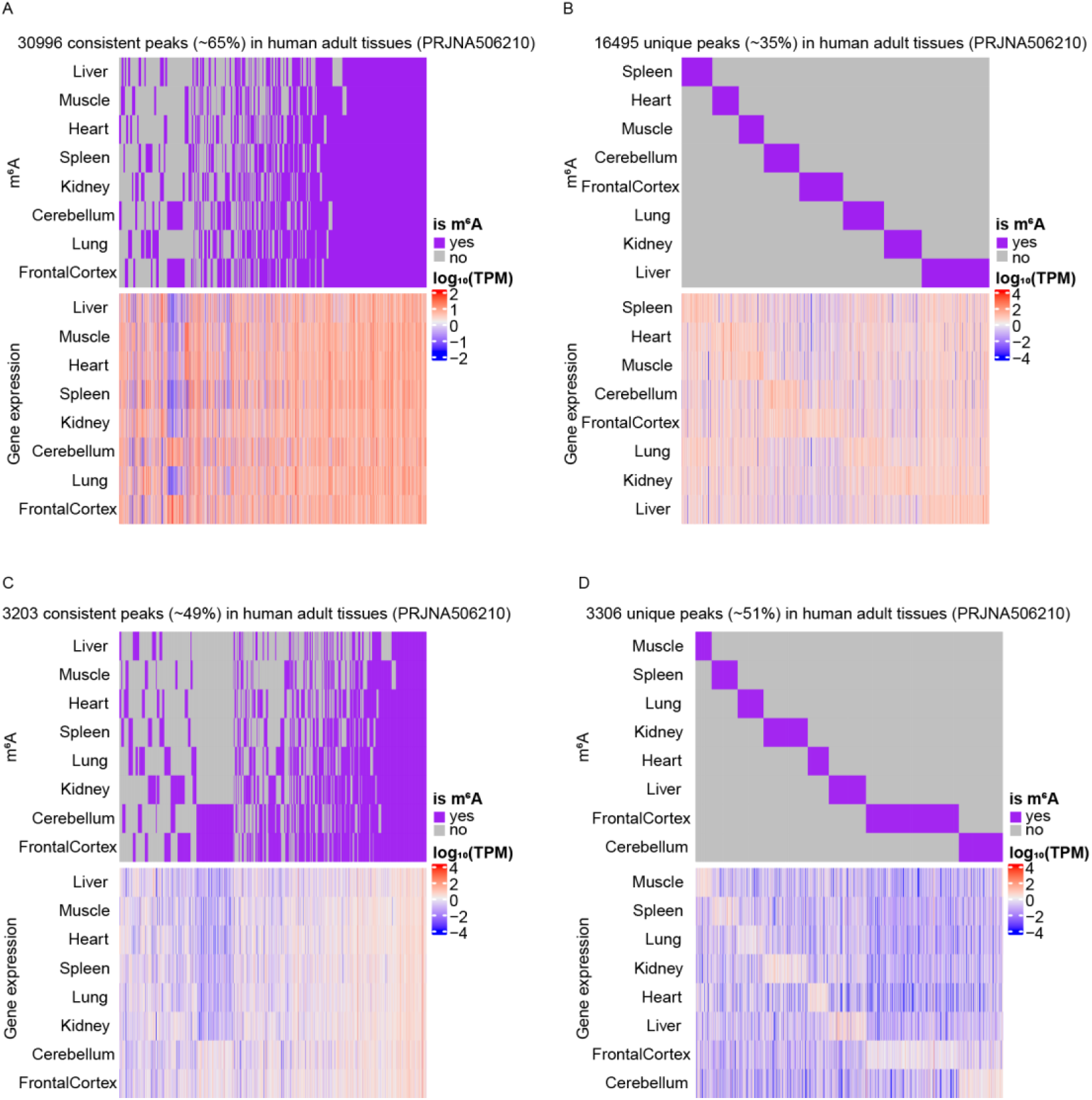
mRNA and lncRNA m6A modification and gene expression profiles in eight different adult tissues. A, mRNA-consistent m6A sites and expression levels of genes marked by these sites. B, mRNA-unique m6A sites and expression levels of genes marked by these sites. C, lncRNA-consistent m6A sites and expression levels of genes marked by these sites. D, lncRNA-unique m6A sites and expression levels of genes marked by these sites. Rows denote different tissues, and columns denote different m6A sites (upper panel) and genes marked by these sites (lower panel).

In the additional eight human fetal tissues (PRJNA464886), fourteen adult tissues (PRJCA001180), eight human cell lines, eleven mouse tissues, and six mouse cell lines, a similar proportion of consistent and unique m6A sites on mRNA and lncRNA was observed, consistent with the above eight adult tissues (PRJNA506210) (Supplementary Figure 12-20). The brain tissue of eight fetal tissues (except for lncRNA), the cerebrum of fourteen adult tissues, and the midbrain of eleven mouse tissues had the most significant number of mRNA- and lncRNA-unique m6A sites, further elucidating that unique m6A sites were critical for brain development (Supplementary Figure 18B, C, E, and Supplementary Figure 20B, C, E). Importantly, we found that if mRNA- and lncRNA-consistent m6A sites appeared in more tissues or cell lines, expression homeostasis of its marked genes was more apparent, indicating m6A sites can help genes stabilize their expression levels among different tissues or cell lines (Supplementary Figure 12-16).

To determine whether the unique m6A sites resulted from gene-specific expression, we first defined transcriptome-specific m6A sites and epi-transcriptome-specific m6A sites (view Methods for details). Then, we examined the ratio of these two types of m6A sites. In human tissues, the proportion of transcriptome-specific mRNA-unique m6A sites ranged from ∼3% to ∼13%, and for lncRNA-unique m6A sites, the proportion ranged from ∼6% to ∼15% (Supplementary Figure 21A-C). In human cell lines, the ratios of transcriptome-specific mRNA- and lncRNA-unique m6A sites were ∼9% and ∼20%, respectively (Supplementary Figure 21D). In mouse tissues, their proportions were ∼3% and ∼7%, respectively, and in mouse cell lines, the proportions were ∼10% and ∼21%, respectively (Supplementary Figure 21E, F). These results showed a low proportion of transcriptome-specific m6A sites, indicating that gene-specific expression was not the main reason for generating unique m6A sites in mRNA and lncRNA. In addition, we also examined the proportion of transcriptome-specific m6A sites in different regions of mRNA (5’-UTR, CDS, and 3’-UTR) and found no preference for 5’-UTR, CDS, and 3’-UTR (Supplementary Figure 21).

### m6A site consistency levels positively correlate with gene expression homeostasis and the CpG density of the gene promoter region

To investigate the relationship between m6A and gene expression homeostasis, we first classified m6A sites into Low-, Middle-, and High-consistency levels (view Methods for details). We defined genes marked by three types of m6A sites and used the Tau value to measure gene expression stability among different tissues or cell lines^35^. We first examined the relationship between the mRNA m6A sites and mRNA gene expression homeostasis. The results showed that the higher the consistency level of the mRNA m6A site in eight different human fetal tissues, the more stable the expression level of its marked gene (Figure 5B). The mRNA m6A consistency level positively correlated with mRNA gene expression homeostasis, consistent with previous research results^26^. The positive correlation was still very substantial in 8 adult tissues (PRJNA506210), 14 adult tissues (PRJCA001180), 8 human cell lines, 11 mouse tissues, and 6 mouse cell lines (Figure 5A, C-F). Next, we examined the relationship between the lncRNA m6A site and lncRNA gene expression homeostasis. Like mRNA, lncRNA m6A consistency level positively correlated with lncRNA gene expression homeostasis, and the positive correlation was significant in humans and mice (Supplementary Figure 22). Finally, for m6A sites on different regions of mRNA (5’-UTR, CDS, and 3’-UTR), the consistency level still positively correlated with the expression homeostasis of its marked mRNA gene (Supplementary Figure 23, 24). In sum, the m6A consistency level was positively associated with gene expression homeostasis, and the positive correlation widely existed in human tissues, human cell lines, mouse tissues, and mouse cell lines.

**Figure 5.**
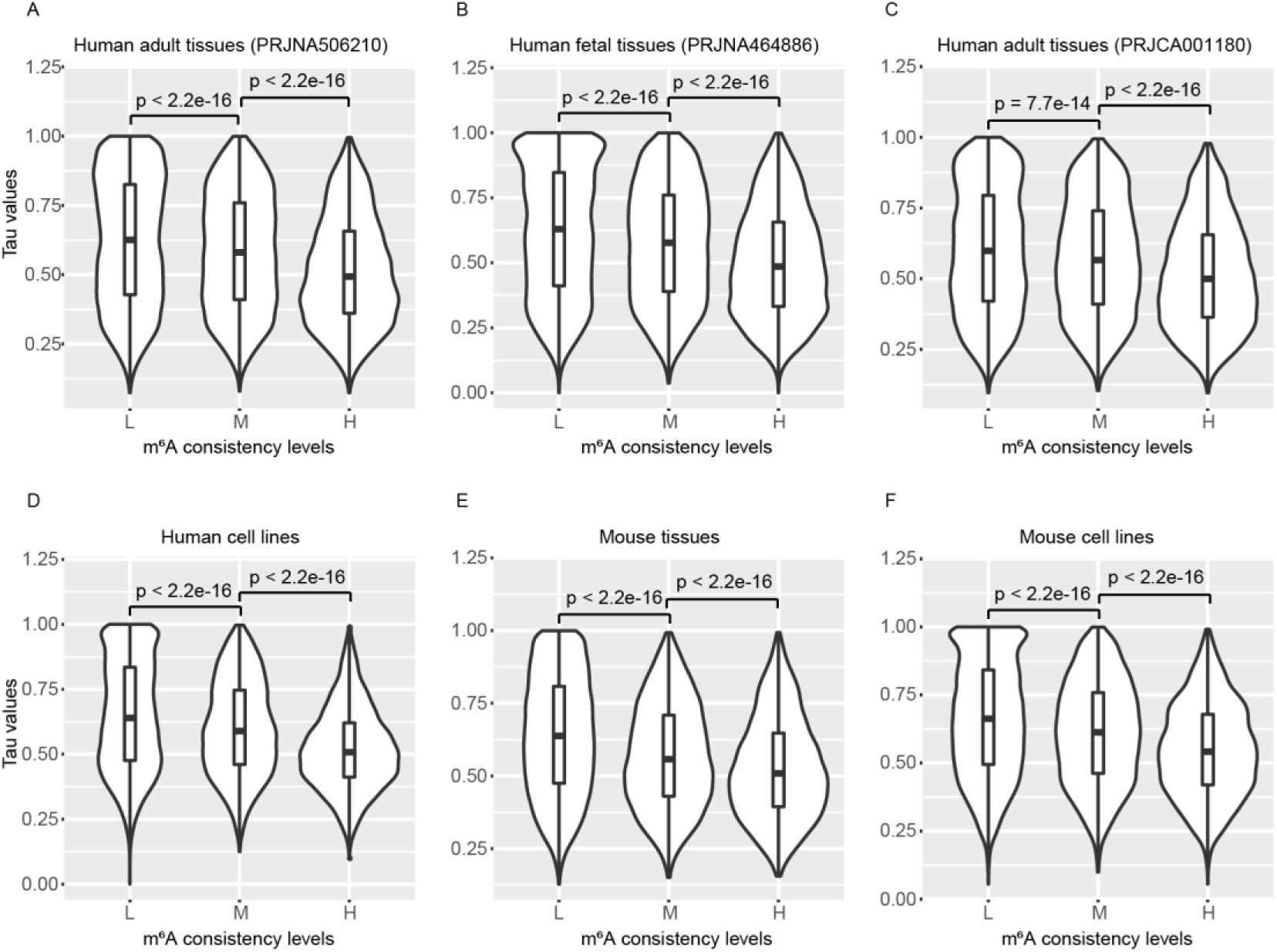
mRNA m6A consistency levels positively correlated with gene expression homeostasis. A, Relative expression stability of m6A-modified genes (L, M, and H sets containing 12228, 5824, and 7130 mRNA genes, respectively) among eight different adult tissues. B, Relative expression stability of m6A-modified genes (L, M, and H sets containing 8735, 4242, and 5279 mRNA genes, respectively) among eight different human fetal tissues. C, Relative expression stability of m6A-modified genes (L, M, and H sets containing 10470, 5104, and 6842 mRNA genes, respectively) among fourteen different adult tissues. D, Relative expression stability of m6A-modified genes (L, M, and H sets containing 10031, 4769, and 4442 mRNA genes, respectively) among eight different human cell lines. E, Relative expression stability of m6A-modified genes (L, M, and H sets containing 12906, 3913, and 2913 mRNA genes, respectively) among eleven different mouse tissues. F, Relative expression stability of m6A-modified genes (L, M, and H sets containing 10653, 4754, and 4449 mRNA genes, respectively) among six different mouse cell lines. Significance was evaluated by the two-sided Mann-Whitney test.

We used the same classification method to study the relationship between m6A and the CpG density of the gene promoter region. The results showed that in eight adult tissues (PRJNA506210), the higher the consistency levels of the mRNA m6A sites, the higher the CpG density of the promoter region of its marked genes. The consistency levels of the mRNA m6A sites positively correlated with the CpG density of the mRNA gene promoter region (Supplementary Figure 25A). The positive correlation still existed in eight fetal tissues, fourteen adult tissues (PRJCA001180), eight human cell lines, eleven mouse tissues, and six mouse cell lines (Supplementary Figure 25B-F). Like mRNA, the higher the consistency level of the lncRNA m6A site, the higher the CpG density of its marked lncRNA gene promoter region (Supplementary Figure 26). The consistency levels of the lncRNA m6A sites positively correlated with the CpG density of the lncRNA gene promoter region. This positive correlation was prevalent in human tissues and cell lines, mouse tissues, and cell lines. Finally, for m6A sites on different regions of mRNA (5’-UTR, CDS, and 3’-UTR), the consistency level still positively correlated with the CpG density of its marked mRNA gene promoter region (Supplementary Figure 27, 28).

### METTL3 preferred binding to promoter regions of genes marked by consistent m6A sites

METTL3, the core catalytic component of the m6A writer complex, can bind directly or indirectly to the genome and regulate the chromatin state and transcription process^36–39^. To determine whether METTL3 preferred promoter regions of genes, we first mapped METTL3 binding sites on the genome (Supplementary Figure 29, 30). In HeLa cells, we detected 3477 METTL3 binding sites. Approximately 96% of the binding sites were in the gene promoter region and highly enriched around the transcription start site (TSS). Mapping METTL3 binding sites to genes showed that METTL3 mainly binds to the promoter regions of 2907 protein-coding genes (Supplementary Figure 29). For mESCs, we detected 16169 METTL3 binding sites. About 46% of the binding sites were in the gene promoter region and highly enriched in TSS, consistent with HeLa cells. METTL3 mainly binds to the promoter regions of 5250 protein-coding genes (Supplementary Figure 30). In summary, METTL3 preferred the promoter regions of protein-coding genes and significantly enriched near TSS.

To study whether METTL3 had a preference for genes marked by consistent m6A sites, we classified the 2907 protein-coding genes detected in HeLa cells into three kinds: genes marked by the most consistent m6A sites (consistent kinds of genes), genes marked by unique m6A sites (unique kinds of genes), and genes marked by both types of m6A sites (both kinds of genes). In the human tissues from three different research groups (PRJNA506210, PRJNA464886, and PRJCA001180), compared to unique kinds of genes, METTL3 had a higher binding ability to consistent and both kinds of genes, indicating that METTL3 had a preference for genes marked by consistent m6A sites (Figure 6A-C). Like human tissues, METTL3 had a higher binding ability to consistent and both kinds of genes in human cell lines (Figure 6D). Next, we classified the 5250 protein-coding genes detected in mESC into three categories. Like human tissues and cell lines, METTL3 had a higher binding ability to both kinds of genes in mouse tissues and cell lines (Supplementary Figure 31). In short, METTL3 preferred binding to genes marked by consistent m6A sites in humans and mice.

**Figure 6.**
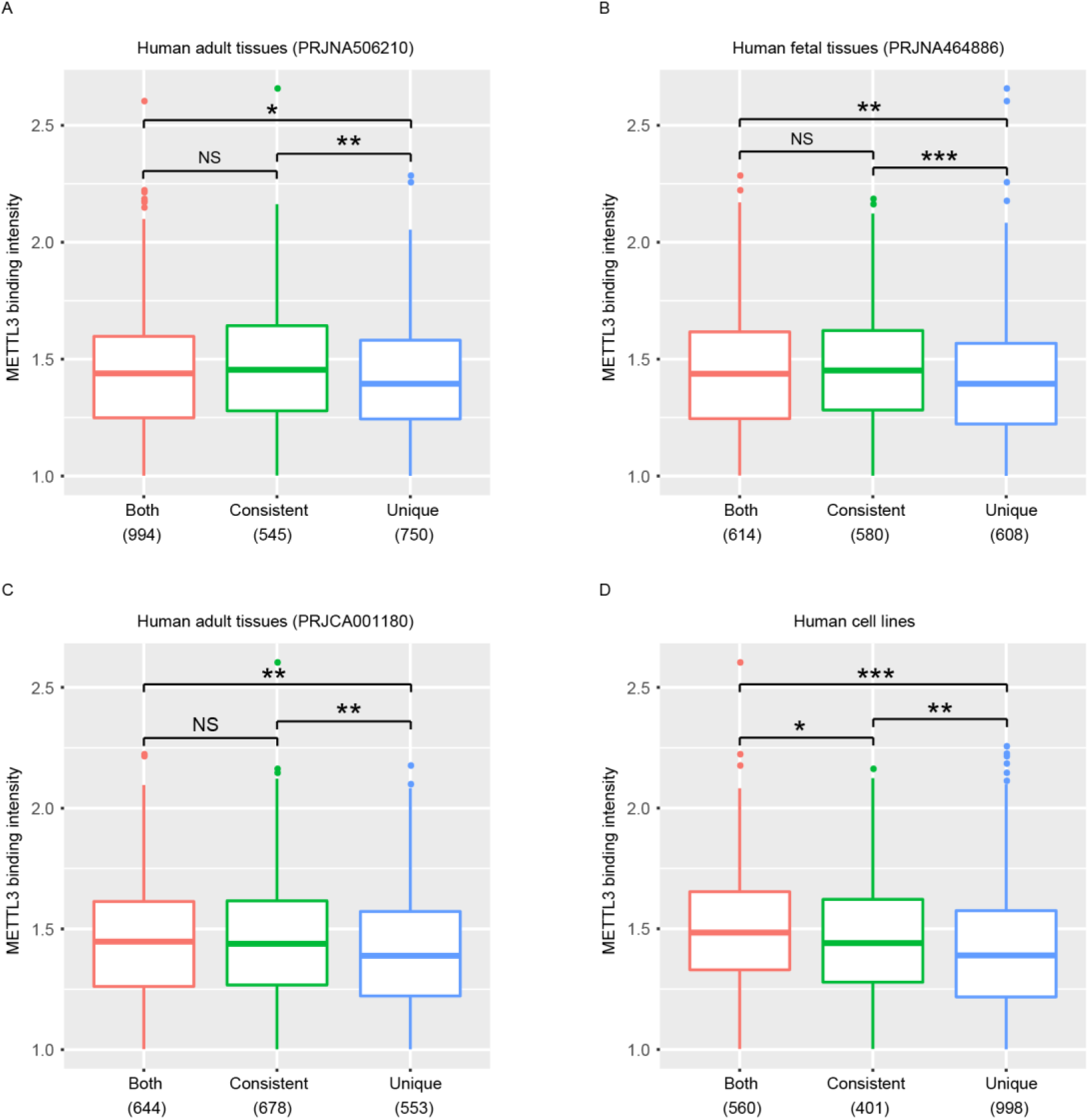
Binding intensity of METTL3 on three types of genes for HeLa. A-D, The METTL3-binding genes in HeLa were classified into three categories, and METTL3 binding intensity was shown for the adult tissues (A, C), human fetal tissues (B), and human cell lines (D). Significance was evaluated by the two-sided Mann-Whitney test. An asterisk (*) denoted that the P value was between 0.01 and 0.05; Two asterisks (**) denoted that the P value was less than 0.01; Three asterisks (***) denoted that the P value was less than 0.001. NS denoted no significant difference. The numbers in parentheses denoted the number of genes.

Our research showed that promoter regions of genes marked by high-consistency m6A sites possessed higher CpG density, and METTL3 preferred binding to promoter regions of genes marked by these m6A sites. Here, we assumed that METTL3 had a preference for gene promoter regions with higher CpG density. To test the assumption, we compared the CpG density in the promoter regions of protein-coding genes bound by METTL3. We classified genes into two categories: genes with high CpG density (High) and low CpG density (Low). The results showed that METTL3 preferred binding to gene promoter regions with high CpG density in HeLa cells and mESC (Supplementary Figure 32).

## Discussion and Conclusion

The widespread use of the m6A-seq detection method has generated a large number of sequencing datasets. Due to inherent flaws in this detection method, m6A-seq data quality varies greatly^24,25^. Here, by evaluating 193 sets of m6A-seq libraries rRNA contamination rate, library complexity NRF, PBC1, PBC2, and antibody enrichment efficiency FRiP, we defined reference values for these metrics, which can evaluate newly produced m6A-seq data quality. Based on the recommended reference values, we screened out high-quality m6A-seq datasets and identified about 1.5 million high-confidence m6A sites.

We constructed m6A modification and gene expression maps for consistent m6A sites and unique m6A sites in mRNA and lncRNA. We found that human and mouse brain tissues possess the most significant number of unique m6A sites. The specific expression of genes can produce some unique m6A sites in mRNA and lncRNA. However, most unique m6A sites are not regulated by gene-specific expression. The production mechanism for these unique m6A sites currently needs to be clarified. The higher the consistency levels of the m6A sites, the more stable the expression levels of its marked genes are across tissues and cell lines. m6A can promote gene (including mRNA and lncRNA) expression homeostasis, and this promotion effect is conserved in humans and mice. The CpG density in the promoter region of genes marked by the high-consistency m6As is very high, indicating that the CpG density of the promoter region may regulate the addition of m6A sites of different consistency levels to nascent transcripts, further regulating the homeostasis of gene expression.

m6A sites of varying consistency levels have specific distribution patterns on the transcriptome. Very stable, hard-coded, and high-consistency m6A sites co-exist in different tissues and cells, and highly enrich near stop codons. Cells- or tissues-unique m6A sites significantly enrich near the start and stop codons, showing a bimodal distribution pattern. These distribution patterns suggest that m6A sites with different consistency levels may be involved in different processes of gene expression regulation.

We identified 40 m6A-binding proteins that preferentially bind to unique m6A sites and 9 m6A-binding proteins that preferentially bind to high-consistency m6A sites. The known m6A readers (YTHDF1-3, RBM15, RBM15B, YTHDC1, and IGF2BP1) prefer to bind to the consistent m6A sites. The high-consistency m6A sites on the transcript can regulate pre-mRNA 3’ end polyadenylation processing, while the unique m6A sites on the transcript play an essential role in regulating mRNA splicing and translation initiation.

On the genome, METTL3 mainly binds to gene promoter regions and has a preference for high CpG promoters. METTL3 also preferentially binds to genes marked by high-consistency m6A sites, indicating that METTL3 and high CpG promoter may be necessary for establishing consistent m6A sites on transcripts.

## Materials and methods

### Data source

The m6A-seq and corresponding RNA-seq sequencing data were downloaded from GEO^40^, EBI^41^, and GSA^42^. Details of m6A-seq sequencing data from human cell lines, human tissues, mouse cell lines, and mouse tissues were provided in Supplementary Tables 1, 2, 3, and 4. ChIP-seq sequencing data of human cell lines and mESCs were downloaded from GEO and EBI databases, and details were shown in Supplementary Tables 5.

The human and mouse genome sequences and annotation files were downloaded from GENCODE^43^.

These URLs were http://ftp.ebi.ac.uk/pub/databases/gencode/Gencode_human/ release_36/GRCh38.primary_assembly.genome.fa.gz, http://ftp.ebi.ac.uk/pub/databases/gencode/Gencode_human/release_36/gencode.v36.an notation. gtf.gz, http://ftp.ebi.ac.uk/pub/databases/gencode/Gencode_mouse/release_M23/GRCm38.prim ary_assembly.genome.fa.gz, and http://ftp.ebi.ac.uk/pub/databases/gencode/Gencode_mouse/release_M23/gencode.vM23.annotation.gtf.gz.

The binding sites of 171 RBPs were downloaded from the POSTAR2 database^30^. The database URL was http://lulab.life.tsinghua.edu.cn/postar/.

### Reads alignment

Use the trimfastq.py (https://github.com/georgimarinov/GeorgiScripts) script^44^ to extract the first 36 bp of the read sequence in the FastQ file. FastQC (https://www.bioinformatics.babraham.ac.uk/projects/fastqc/) was used to check the quality of the reads. Use HISAT2^45^ to align reads to the mouse or human genome. For paired-end sequencing, use the parameters "--no-mixed --no-discordant --no-unal". For single-end sequencing, use "--no-unal". Use SAMtools (http://www.htslib.org/doc/samtools.html) to convert SAM files into BAM files. To avoid bias, we removed reads mapped to the mitochondrial genome. Use the infer_experiment.py script in the RSeQC package to infer whether RNA-seq and m6A-seq libraries are strand-specific^46^. If the library is strand-specific, divide the BAM file into two files (forward and reverse) according to the SAM flag. Use the bamCoverage tool in deepTools^47^ to convert BAM files into bigwig files. Visualize read signal distribution using IGV^48^.

### m6A sites detection

Use BEDTools^49^ to convert BAM files into BED files. Use MACS2^50^ to detect m6A sites, and the parameter of "-g hs -f BEDPE -p 1e-1 --keep-dup all --nomodel --min-length 150 - -max-gap 50" was set. Change the parameter to "-g mm" for the mouse genome. For samples with strand-specific library construction, m6A sites were detected separately by aligning reads to the positive and negative strands of the genome. Use the IDR framework^44^ to identify co-occurring m6A sites between replicate samples. The IDR framework does not rely on artificial p values settings. Use IGV to check whether the m6A site is credible.

### m6A site annotation, distribution pattern analysis, and motif analysis

ChIPseeker^51^ was used to annotate the detected m6A site to the gene, and IGV was used to check the accuracy of the annotation. The MetaPlotR pipeline^52^ was used to analyze the distribution pattern of m6A sites on the transcriptome. Use the script findMotifsGenome.pl in HOMER (http://homer.ucsd.edu/homer/motif/) software to check the m6A site-enriched motif sequences.

### m6A-seq library quality evaluation

Use the metrics NRF, PBC1, PBC2, and FRiP defined by ENCODE to evaluate library quality. The URL is https://www.encodeproject.org/data-standards/terms/#library, where NRF, PBC1, and PBC2 were used to assess library complexity. FRiP was used to evaluate antibody enrichment efficiency.

(1) rRNA contamination rate M0

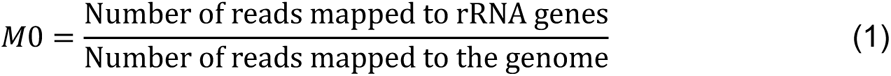
(2) NRF (Non-Redundant Fraction)

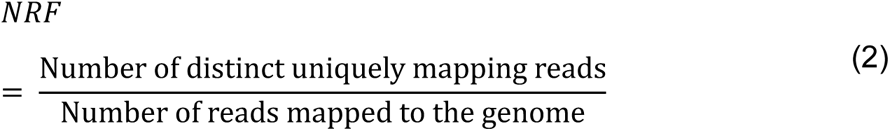
(3) PBC1 (PCR Bottlenecking Coefficient 1)

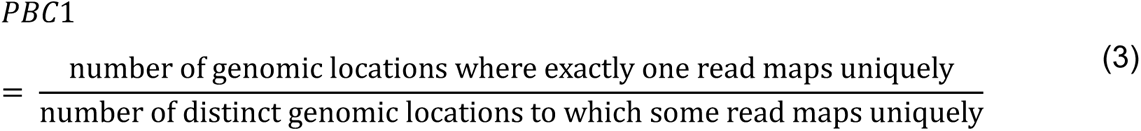
(4) PBC2 (PCR Bottlenecking Coefficient 2)

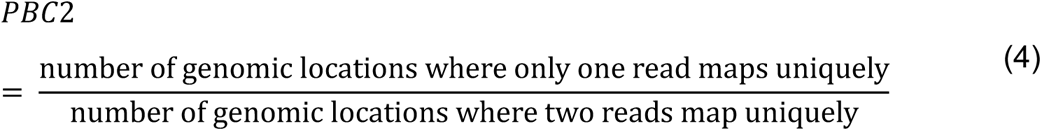
(5) FRiP (Fraction of reads in peaks)

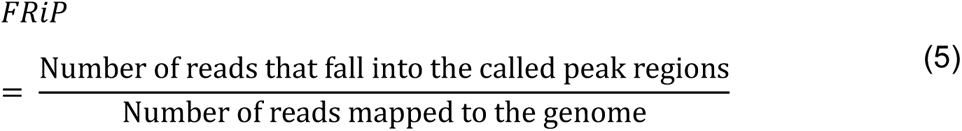

## RNA-seq data analysis

Use featureCounts^53^ to calculate the count of reads mapped to the gene, and use R script to calculate the gene expression TPM value.

## Classifying m6A sites into different consistency levels

Use the BEDTools command to merge m6A sites. The parameters are "-s -d 10 -c 4,6 -o collapse,collapse". Based on the number of cell lines or tissues, m6A sites were classified into different consistency levels. In the m6A and gene expression landscape section, we classified m6A sites into mRNA- or lncRNA-consistent m6A sites and mRNA- or lncRNA-unique m6A sites. m6A sites present simultaneously in at least two tissues or cell lines are defined as mRNA- or lncRNA-consistent m6A sites. m6A sites present only in one tissue or cell line are defined as mRNA- or lncRNA-unique m6A sites.

In the m6A and gene expression homeostasis section, m6A sites are classified into three categories. For 8 adult tissues (PRJNA506210), m6A sites present in 1 to 3 tissues are defined as Low-consistency m6A sites (L); m6A sites present in 4 to 6 tissues are defined as Middle-consistency m6A sites (M); m6A sites present in 7 to 8 tissues are defined as High-consistency m6A sites (H). For 8 fetal tissues (PRJNA464886), the classification method is the same as 8 adult tissues. For 14 adult tissues, m6A sites present in 1 to 5 tissues are defined as Low-consistency m6A sites (L); m6A sites present in 6 to 10 tissues are defined as Middle-consistency m6A sites (M); m6A sites present in 11 to 14 tissues are defined as High-consistency m6A sites (H). For 8 human cell lines, m6A sites present in 1 to 3 cell lines are defined as Low-consistency m6A sites (L); m6A sites present in 4 to 6 cell lines are defined as Middle-consistency m6A sites (M); m6A sites present in 7 to 8 cell lines are defined as High-consistency m6A sites (H). For 11 mouse tissues, m6A sites present in 1 to 3 tissues are defined as Low-consistency m6A sites (L); m6A sites present in 4 to 7 tissues are defined as Middle-consistency m6A sites (M); m6A sites present in 8 to 11 tissues are defined as High-consistency m6A sites (H). For 6 mouse cell lines, m6A sites present in 1 to 2 cell lines are defined as Low-consistency m6A sites (L); m6A sites present in 3 to 4 cell lines are defined as Middle-consistency m6A sites (M); m6A sites present in 5 to 6 cell lines are defined as High-consistency m6A sites (H).

### Tissue- or cell-specific m6A sites analysis

m6A sites only present in one tissue or cell line are defined as unique m6A sites. Unique m6A sites are classified into two categories according to whether the genes marked by the unique m6A sites are exclusively expressed in the specific tissue or cell line. Unique m6A sites produced by gene-specific expression are defined as transcriptome-specific unique m6As. Unique m6A sites resulting from epi-transcriptome regulation are defined as epi- transcriptome-specific unique m6As. Tissue- or cell-line-specific gene expression is defined as the expression level in this tissue or cell line being more than ten times higher than in other tissues or cell lines.

### Gene expression homeostasis analysis

The tissue-specific index Tau value^35^ was used to measure gene expression homeostasis among tissues or cell lines.

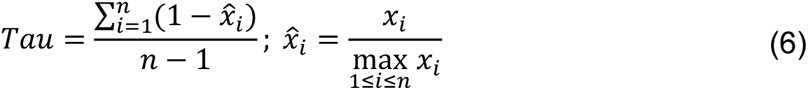

Xi represents the TPM value of the gene in tissue i, and n represents the number of tissues. The lower the Tau value, the more stable the gene expression level is between tissues or cell lines.

### CpG density of gene promoter region

Define 1000 bp upstream and 100 bp downstream of the transcription start site (TSS) as the gene promoter region. For genes with multiple TSS, the average CpG o/e ratio is used as the CpG o/e ratio of the gene. Use the script CpGoe.pl in the Notos tool^54^ to calculate the CpG o/e ratio of each gene promoter region. The calculation method is as follows:

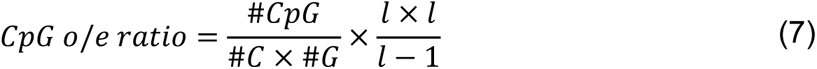

C, #G, and #CpG represent the number of C, G, and CG bases in the promoter sequence, respectively, and l represents the length of the promoter sequence.

### Enrichment analysis between RBP binding sites and m6A sites of varying consistency levels

To identify novel m6A-binding proteins that prefer m6A sites of different consistency levels, we performed enrichment analysis between m6A sites and RBP binding sites. The 171 RBP binding sites were downloaded from the POSTAR2 database, and then the chi-square test^55^ was used to conduct enrichment analysis. The steps are as follows:

(1) Use the BEDTools intersect command to calculate the number of overlapping sites between 171 RBPs binding sites and m6A sites of different consistency levels. The number of overlapping sites is the observed number and is named O (Observed).
(2) Use the R function chisq.test to calculate the expected number of overlapping sites, named E (Expected). The calculation formula is as follows: 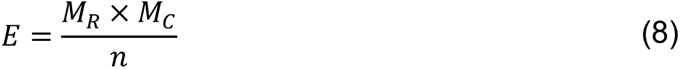 E represents the expected number of overlapping sites between RBP binding sites such as YTHDF1 and m6A sites of a specific consistency level. MR represents the sum of the rows, and MC represents the sum of the columns. n represents the sum of overlapping sites between RBP binding sites and m6A sites.
(3) Calculation of Odds ratio 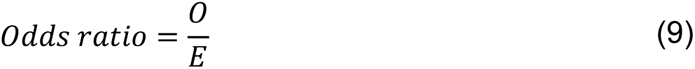
(4) Calculation of Pearson residuals (r) 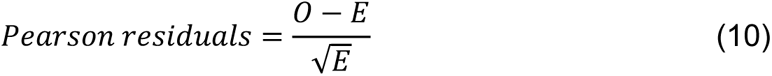
(5) Calculation of the chi-square statistic 𝑋^2^ 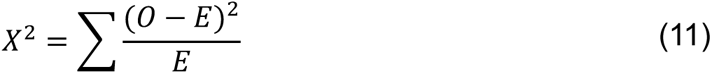
(6) Calculation of Relative contribution 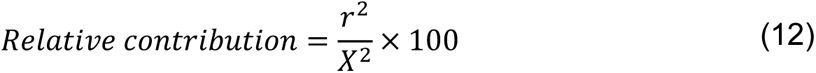

To identify potential m6A-binding proteins, we use the following thresholds: (1) Odds ratio >= 1.2 or Odds ratio <= 0.8; (2) relative contribution > 0.2.

### Protein-protein interaction network analysis

Use the STRING database^56^ to construct an interaction network between potential m6A- binding proteins and known m6A regulators (writer, reader, eraser), retaining only experimentally confirmed interaction information. Visualize interaction networks using Cytoscape^57^.

### ChIP-seq data analysis

Use the trimfastq.py script to extract the first 36 bp of the read sequence. Use FastQC to check the read quality. Reads were mapped to the human or mouse genome using Bowtie2^58^. Use the bamCoverage tool in deepTools to convert the BAM file into a bigwig file. Use the computeMatrix and plotHeatmap tools to analyze the distribution of read signals in the gene body region. Use IGV to visualize read signal distribution. MACS2 is used to detect the binding sites of the target protein on the genome. Use ChIPseeker to examine the distribution pattern of binding sites in different genomic regions.

### Statistical analysis and visualization

Use the R function Wilcox.test to test significant differences between two samples. The results are visualized using the R package ggplot2^59^. Heatmaps are generated using the R package ComplexHeatmap^60^.

