## Supplementary materials for "Comprehensive analysis across mammalian tissues and cells decipher the underlying mechanism of m6A specificity"

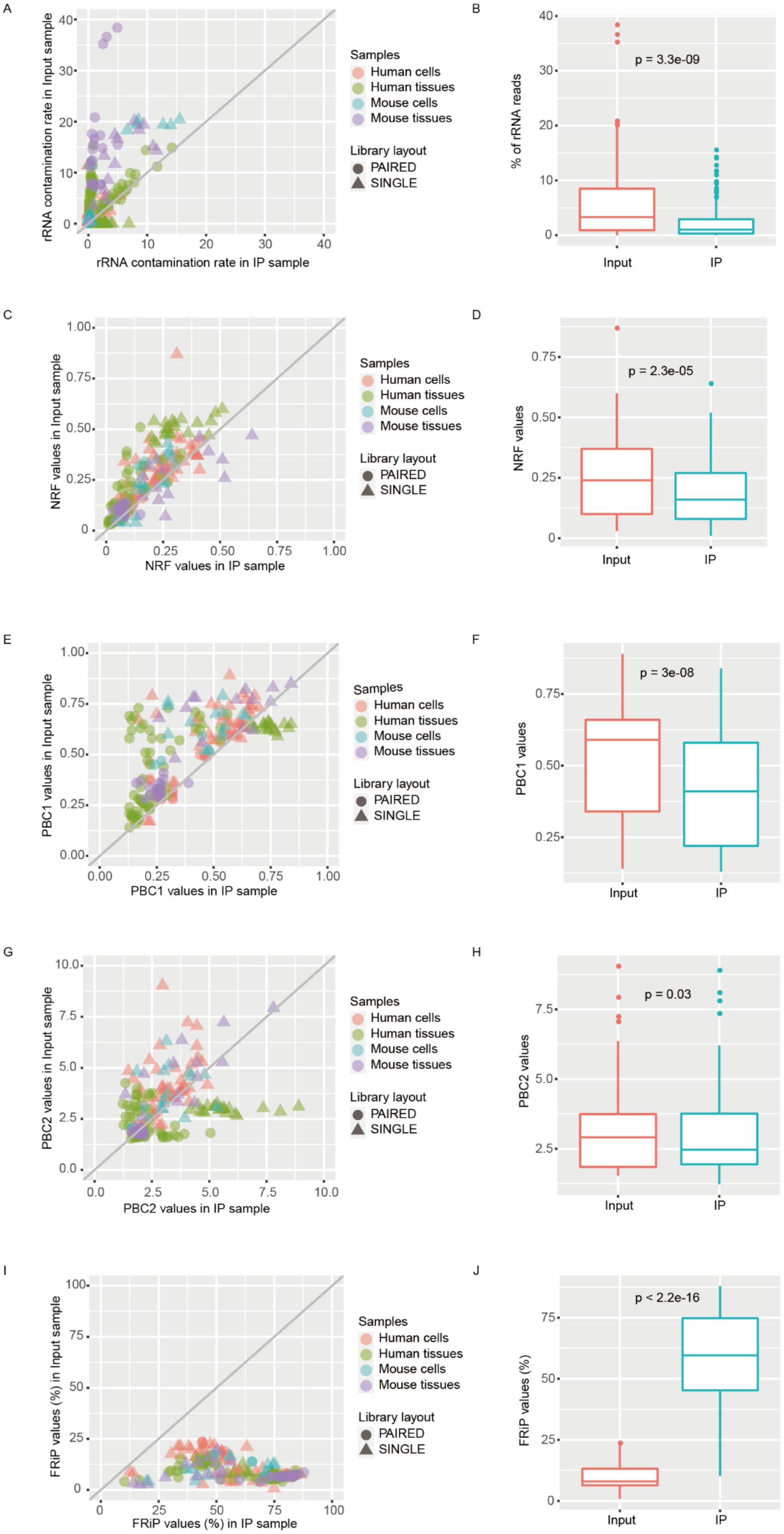
A
rRNA contamination rate in Input sample
rRNA contamination rate in IP sample
Samples
Human cells
Human tissues
Mouse cells
Mouse tissues
Library layout
PAIRED
SINGLE
B
% of rRNA reads
p = 3.3e-09
Input
IP
C
NRF values in Input sample
NRF values in IP sample
Samples
Human cells
Human tissues
Mouse cells
Mouse tissues
Library layout
PAIRED
SINGLE
D
NRF values
p = 2.3e-05
Input
IP
E
PBC1 values in Input sample
PBC1 values in IP sample
Samples
Human cells
Human tissues
Mouse cells
Mouse tissues
Library layout
PAIRED
SINGLE
F
PBC1 values
p = 3e-08
Input
IP
G
PBC2 values in Input sample
PBC2 values in IP sample
Samples
Human cells
Human tissues
Mouse cells
Mouse tissues
Library layout
PAIRED
SINGLE
H
PBC2 values
p = 0.03
Input
IP
I
FRiP values (%) in Input sample
FRiP values (%) in IP sample
Samples
Human cells
Human tissues
Mouse cells
Mouse tissues
Library layout
PAIRED
SINGLE
J
FRiP values (%)
p < 2.2e-16
Input
IP

Supplementary Figure 1 **Quality evaluation of 193 pairs of IP and input samples.** A, B, Distribution of rRNA contamination rates in IP and Input samples. C, D, Distribution of NRF values in IP and Input samples. E F, Distribution of PBC1 values in IP and Input samples. G, H, Distribution of PBC2 values in IP and Input samples. I, J, Distribution of FRiP values in IP and Input samples.

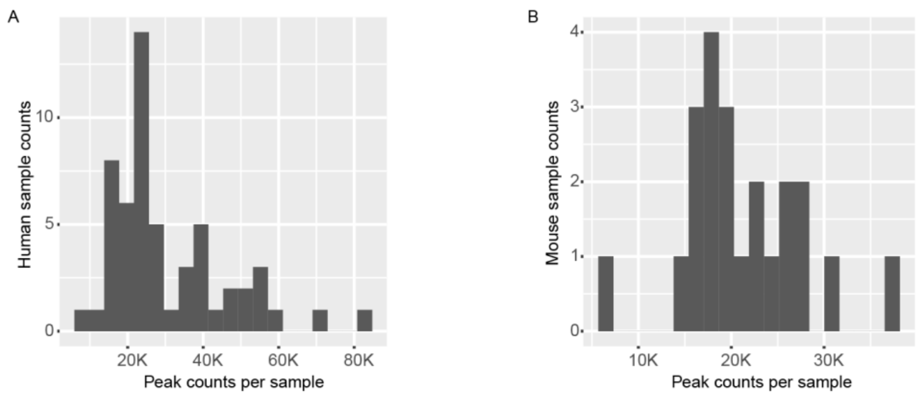


Supplementary Figure 2 **Number of detected m6A sites.** The number of m6A sites detected in human samples (A) and mouse samples (B).

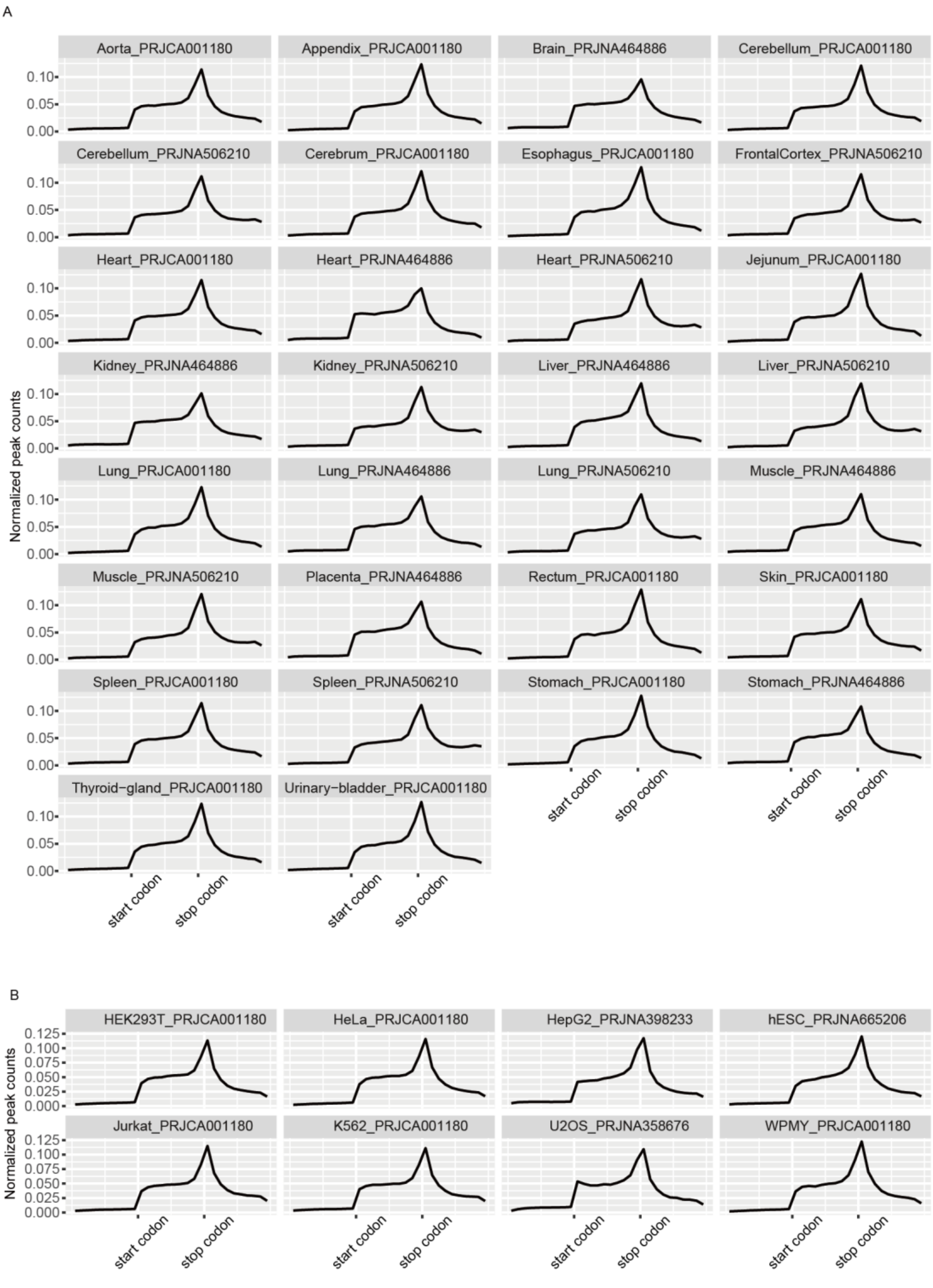


Supplementary Figure 3 **The distribution pattern of m6A sites on the transcriptome.** The distribution pattern of m6A sites in 30 human tissues (A) and 8 human cell lines (B).

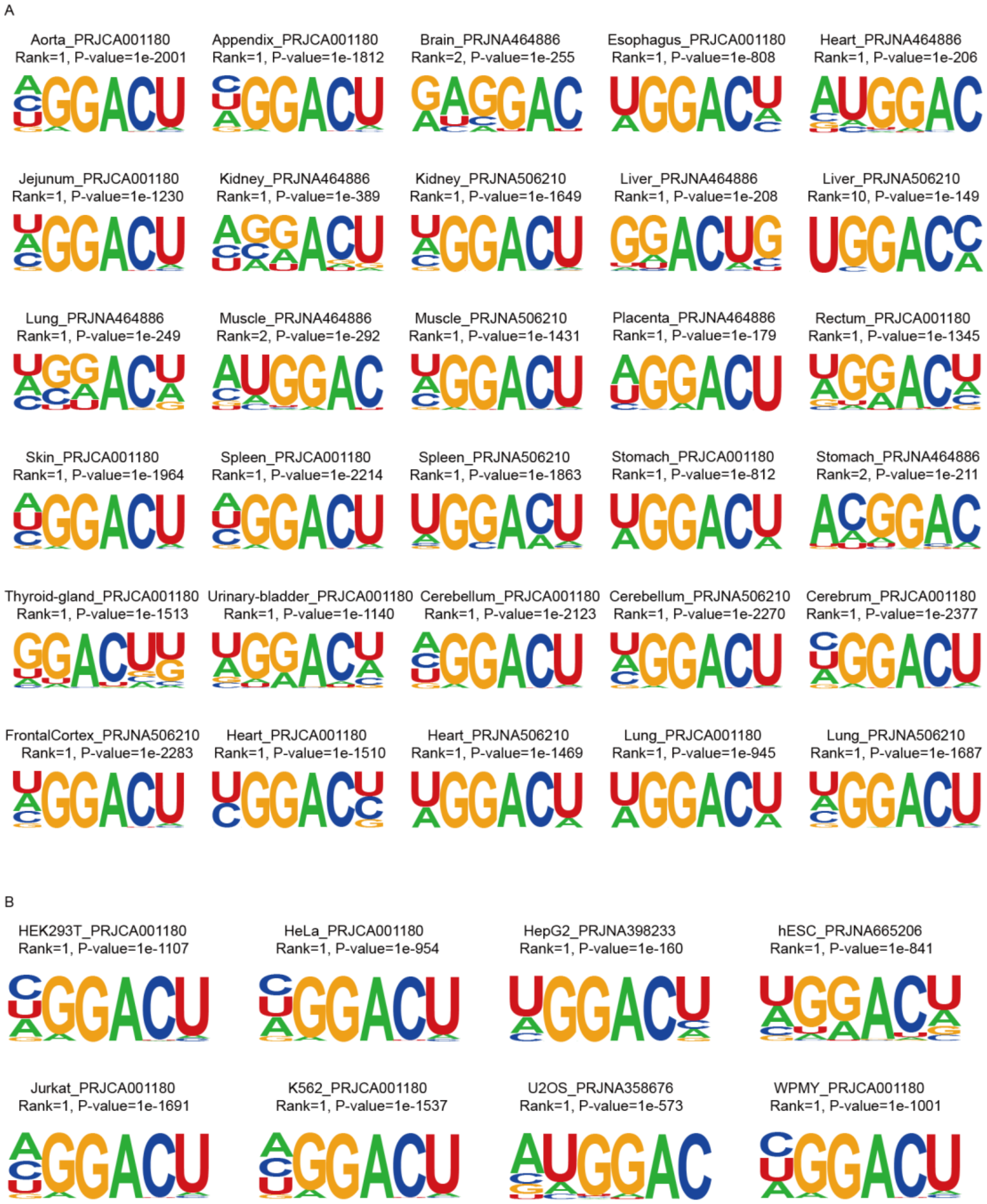


Supplementary Figure 4 **Consensus motif sequences significantly enriched at m6A sites.** Enriched motif sequences in 30 human tissues (A) and 8 human cell lines (B).

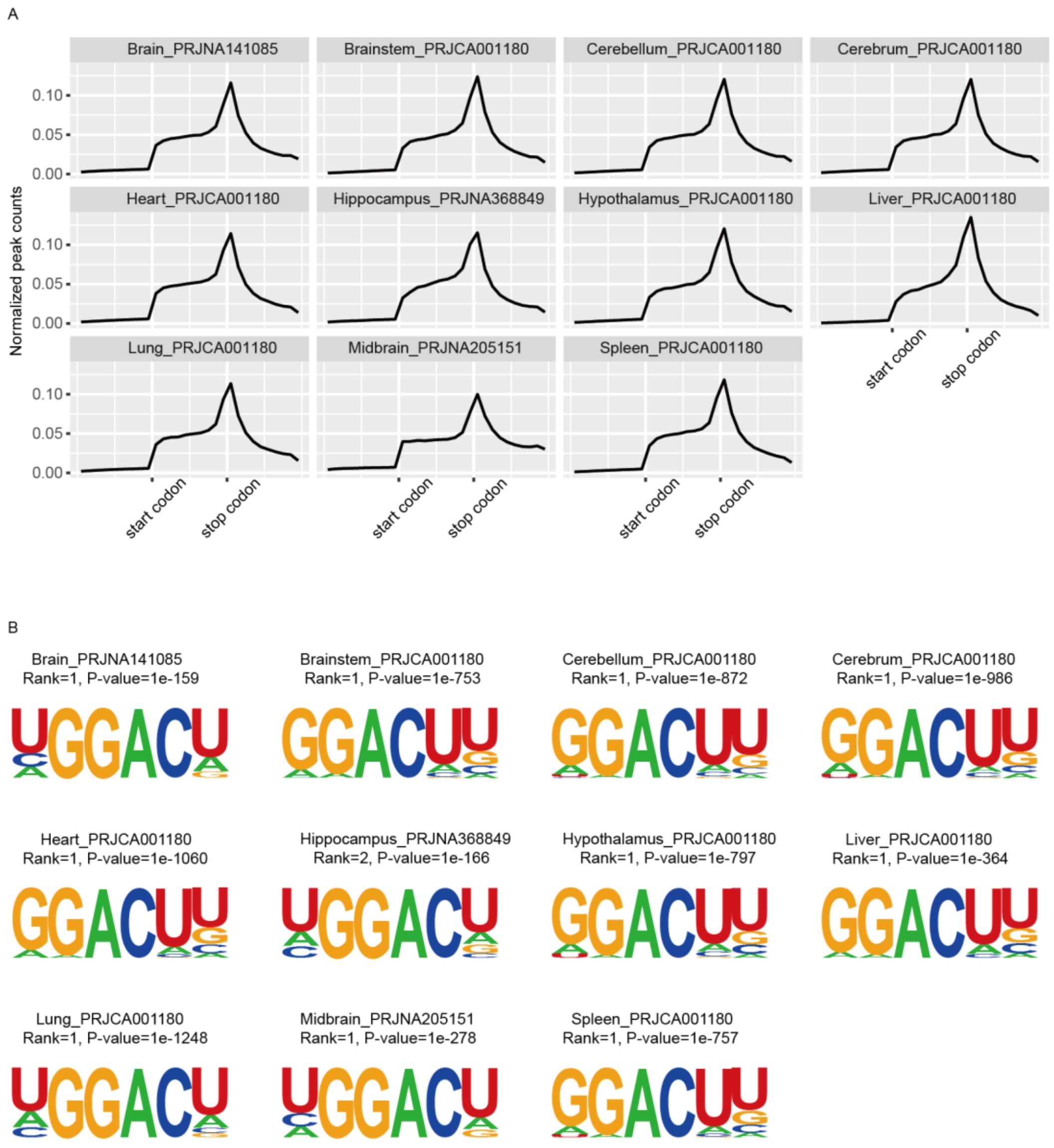


Supplementary Figure 5 A, The distribution pattern of m6A sites along the transcriptome for 11 mouse tissues. B, Consensus motif sequences significantly enriched at m6A sites for 11 mouse tissues.

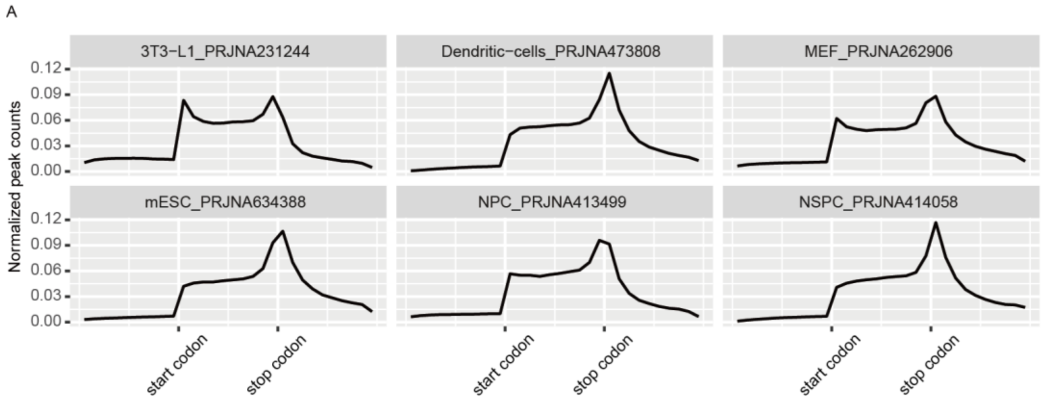


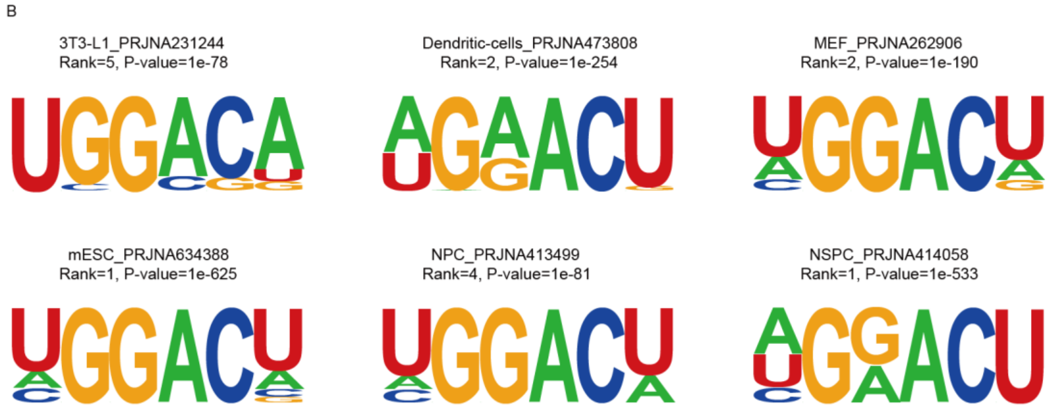


Supplementary Figure 6 A, The distribution pattern of m6A sites along the transcriptome for 6 mouse cell lines. B, Consensus motif sequences significantly enriched at m6A sites for 6 mouse cell lines.

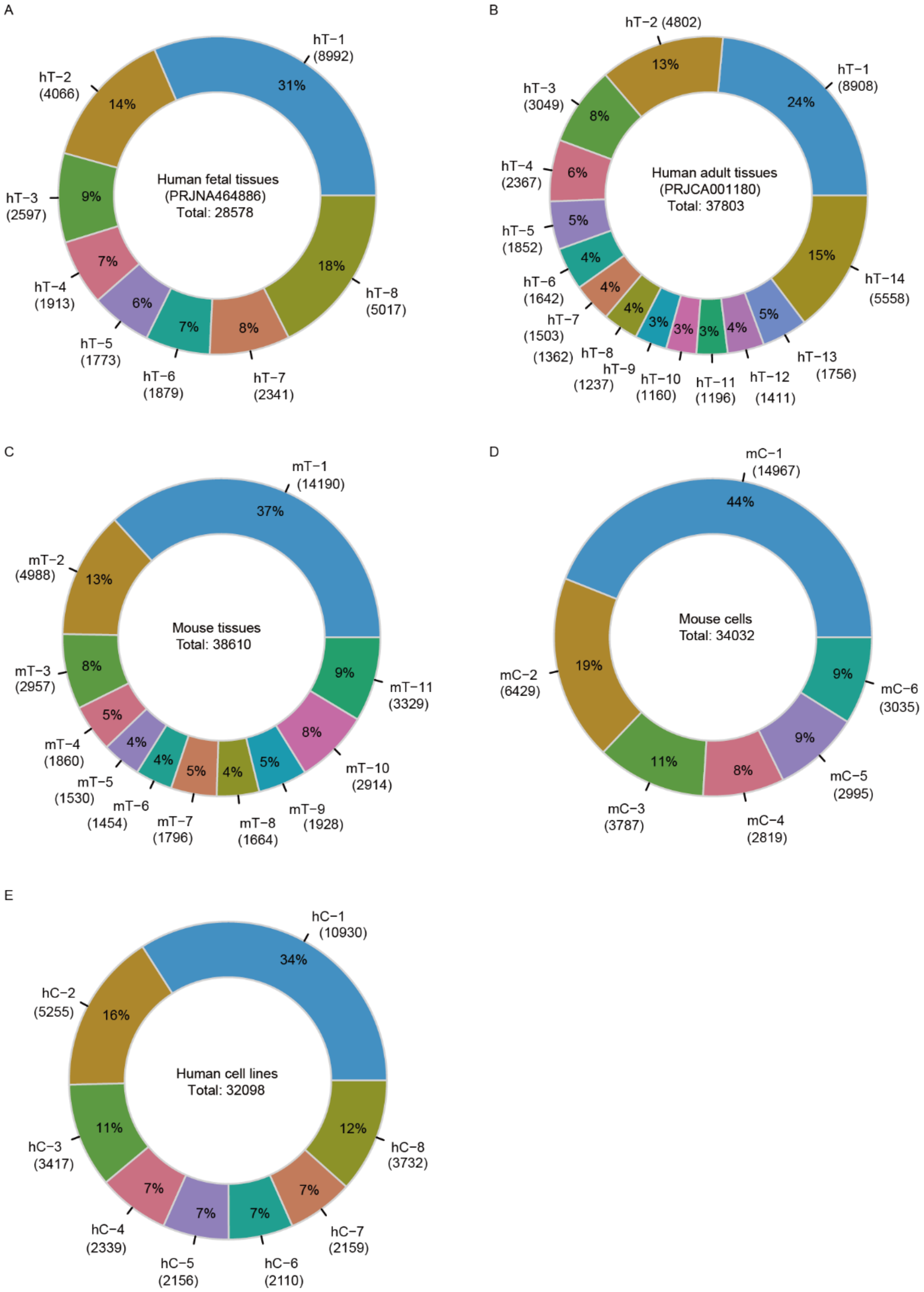


Supplementary Figure 7 **The number and proportion of m6A sites of different consistency levels in human tissues and cell lines, and mouse tissues and cell lines.** A, The m6A sites detected in eight different human fetal tissues are classified into eight consistency levels, and the number and proportion of m6A sites of eight consistency levels are shown. B, The m6A sites detected in fourteen different adult tissues are classified into fourteen consistency levels, and the number and proportion of m6A sites of fourteen consistency levels are shown. C, m6A sites detected in eleven different mouse tissues are classified into eleven consistency levels, and the number and proportion of m6A sites of eleven consistency levels are shown. D, m6A sites detected in six different mouse cell lines are classified into six consistency levels, and the number and proportion of m6A sites of six consistency levels are shown. E, m6A sites detected in eight different human cell lines are classified into eight consistency levels, and the number and proportion of m6A sites of eight consistency levels are shown. hT denotes human tissue. hT-1 denotes that m6A is present in one type of human tissue. hT-8 denotes that m6A is present in eight different human tissues simultaneously. mT denotes mouse tissue. mT-1 denotes that m6A is present in one type of mouse tissue. mT-11 denotes that m6A is present in eleven different mouse tissues simultaneously.

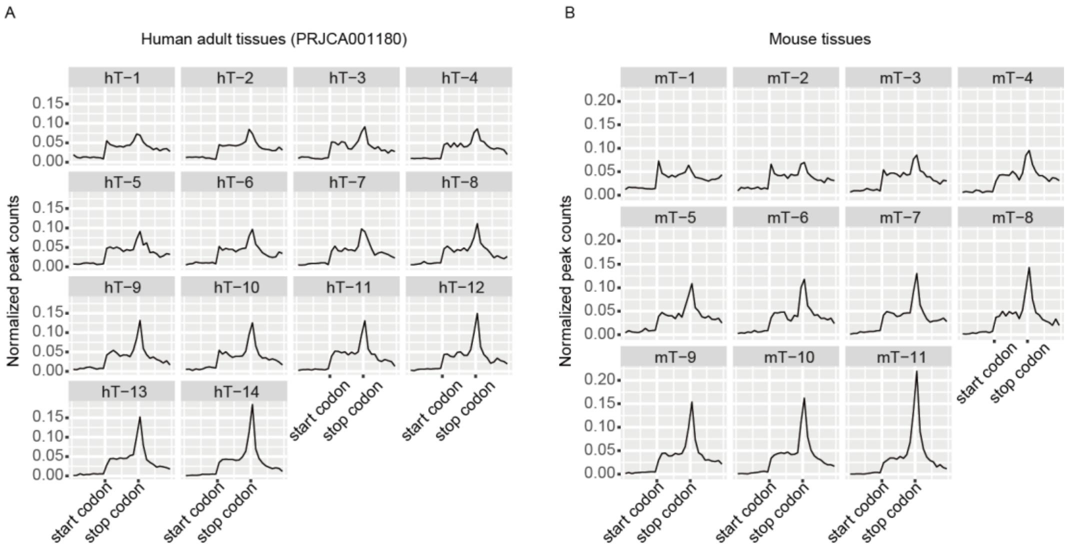


Supplementary Figure 8 **The distribution patterns of m6A sites with different consistency levels in human and mouse tissues.** A, Distribution patterns of m6A sites of fourteen consistency levels defined by fourteen different adult tissues along the transcriptome. B, Distribution patterns of m6A sites of eleven consistency levels defined by eleven different mouse tissues along the transcriptome. hT denotes human tissue. hT-1 denotes that m6A is present in one type of human tissue. hT-14 denotes that m6A is present in fourteen different human tissues simultaneously. mT denotes mouse tissue. mT-1 denotes that m6A is present in one type of mouse tissue. mT-11 denotes that m6A is present in eleven different mouse tissues simultaneously.

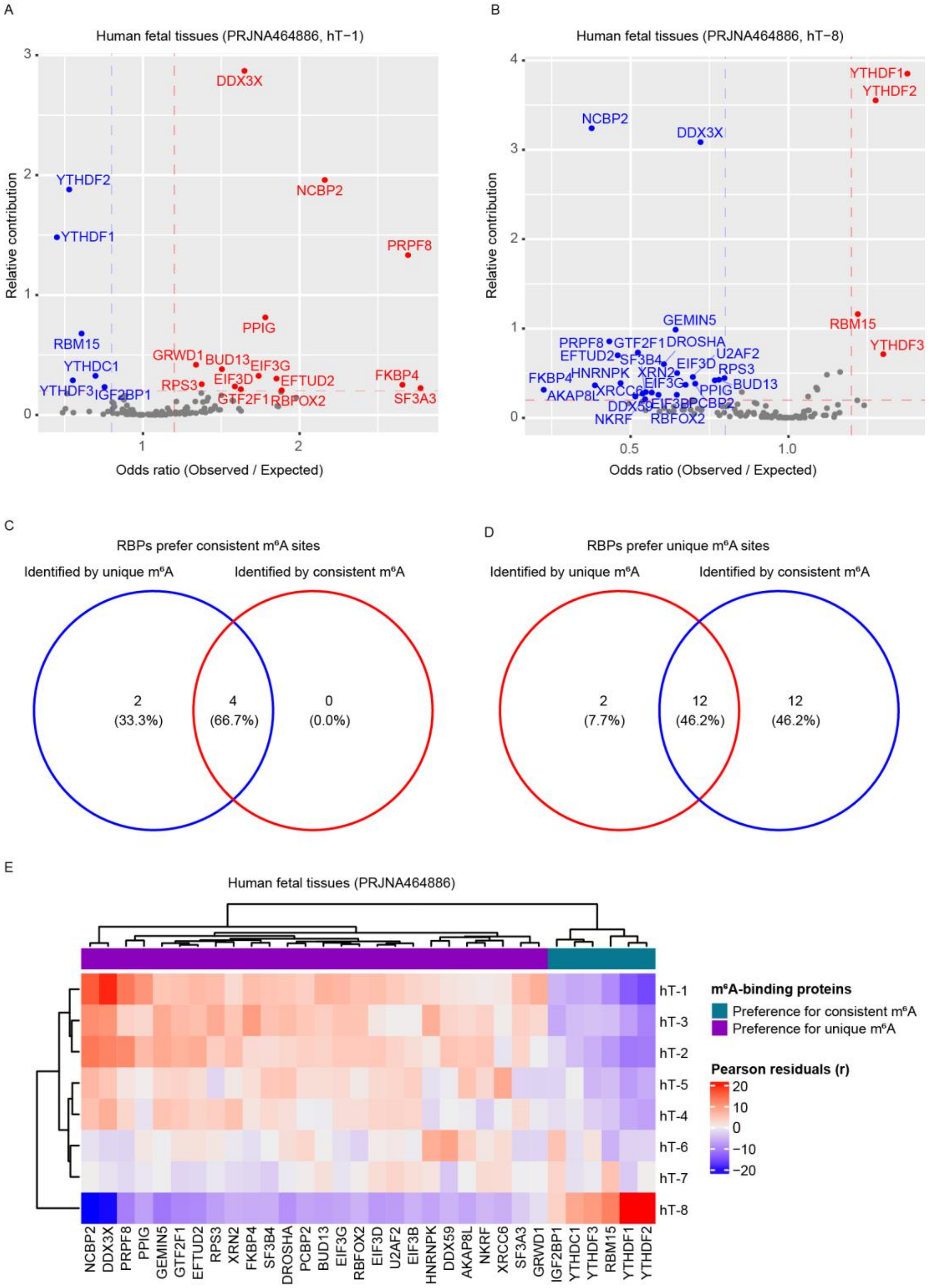


Supplementary Figure 9 **m6A-binding proteins identified by m6A sites of eight consistency levels in eight human fetal tissues.** A, Six m6A-binding proteins that prefer binding to consistent m6A sites and fourteen m6A-binding proteins that prefer binding to unique m6A sites are identified through unique m6A sites enrichment analysis. B, Four m6A-binding proteins that prefer binding to consistent m6A sites and twenty-four m6A-binding proteins that prefer binding to unique m6A sites are identified through the most consistent m6A sites enrichment analysis. C, Overlap analysis of m6A-binding proteins that prefer binding to consistent m6A sites. D, Overlap analysis of m6A-binding proteins that prefer binding to unique m6A sites. E, The affinity of thirty-two m6A-binding proteins to m6A sites of eight consistency levels in eight human fetal tissues.

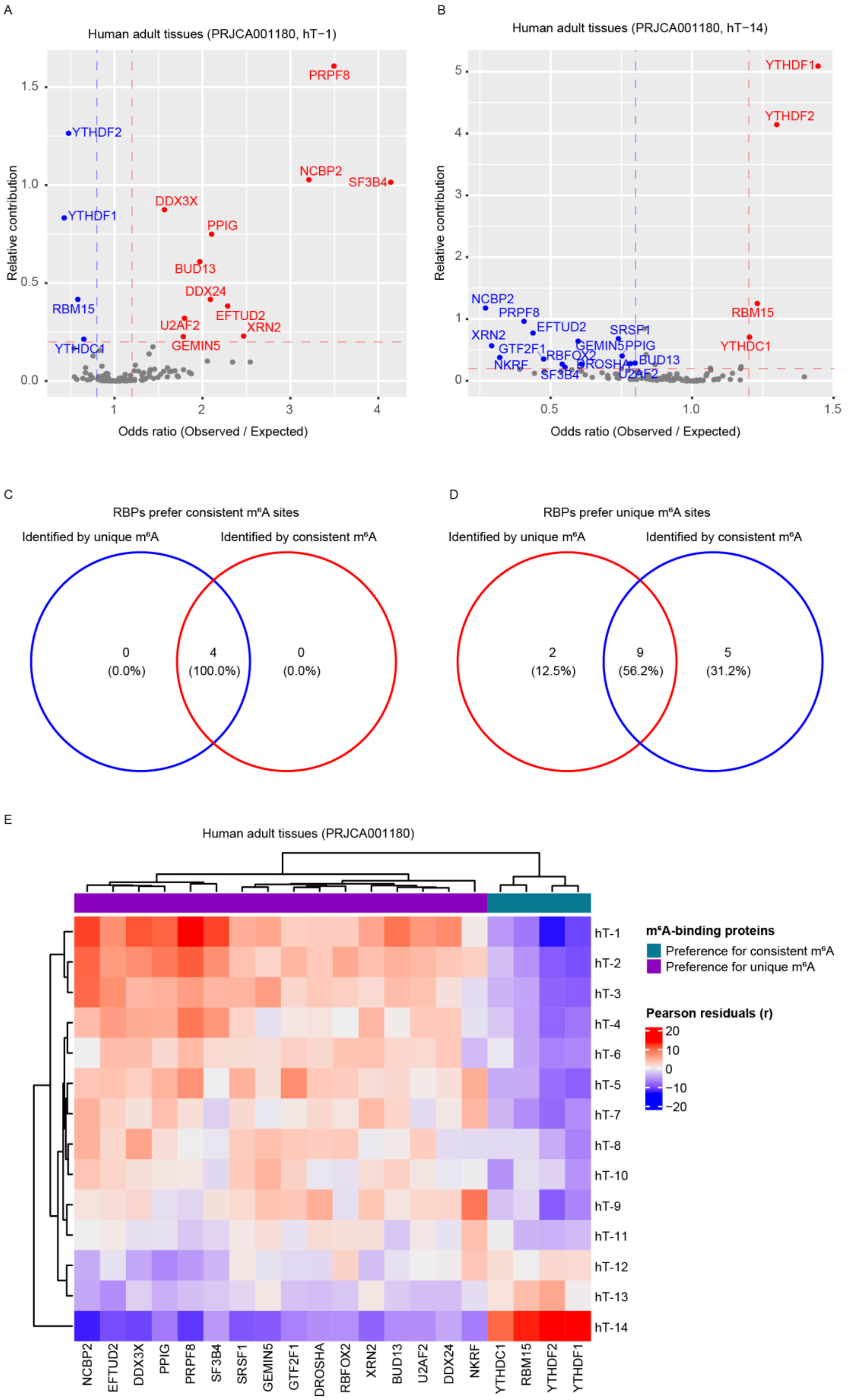


Supplementary Figure 10 **m6A-binding proteins identified by m6A sites of fourteen**

**consistency levels in fourteen adult tissues.** A, Four m6A-binding proteins that prefer binding to consistent m6A sites and eleven m6A-binding proteins that prefer binding to unique m6A sites are identified through unique m6A sites enrichment analysis. B, Four m6A-binding proteins that prefer binding to consistent m6A sites and fourteen m6A-binding proteins that prefer binding to unique m6A sites are identified through the most consistent m6A sites enrichment analysis. C, Overlap analysis of m6A-binding proteins that prefer binding to consistent m6A sites. D, Overlap analysis of m6A-binding proteins that prefer binding to unique m6A sites. E, The affinity of twenty m6A-binding proteins to m6A sites of fourteen consistency levels in fourteen adult tissues.

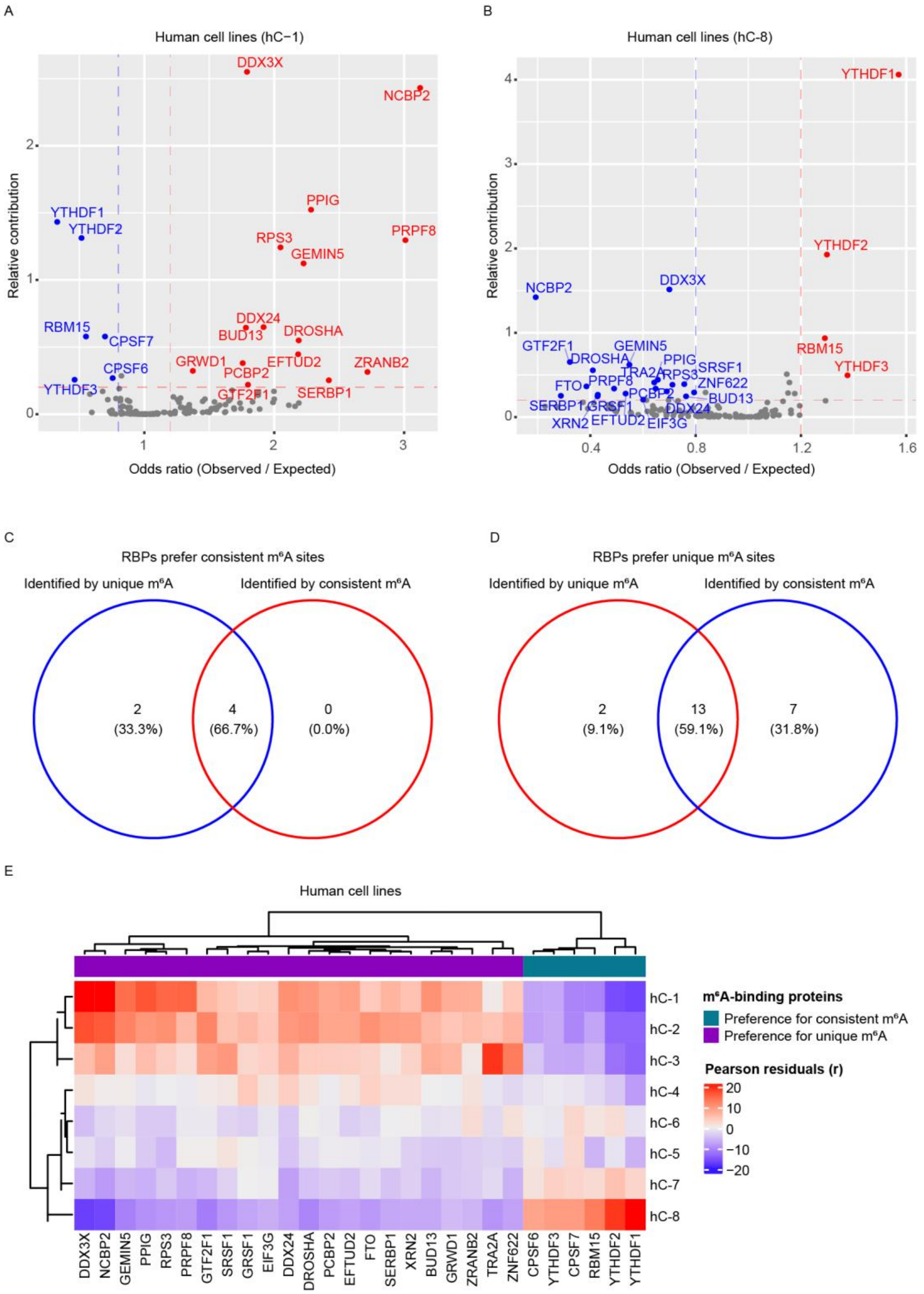


Supplementary Figure 11 **m6A-binding proteins identified by m6A sites of eight consistency levels in eight human cell lines.** A, Six m6A-binding proteins that prefer binding to consistent m6A sites and fifteen m6A-binding proteins that prefer binding to unique m6A sites are identified through unique m6A sites enrichment analysis. B, Four m6A-binding proteins that prefer binding to consistent m6A sites and twenty m6A-binding proteins that prefer binding to unique m6A sites are identified through the most consistent m6A sites enrichment analysis. C, Overlap analysis of m6A-binding proteins that prefer binding to consistent m6A sites. D, Overlap analysis of m6A-binding proteins that prefer binding to unique m6A sites. E, The affinity of twenty-eight m6A-binding proteins to m6A sites of eight consistency levels in eight human cell lines.

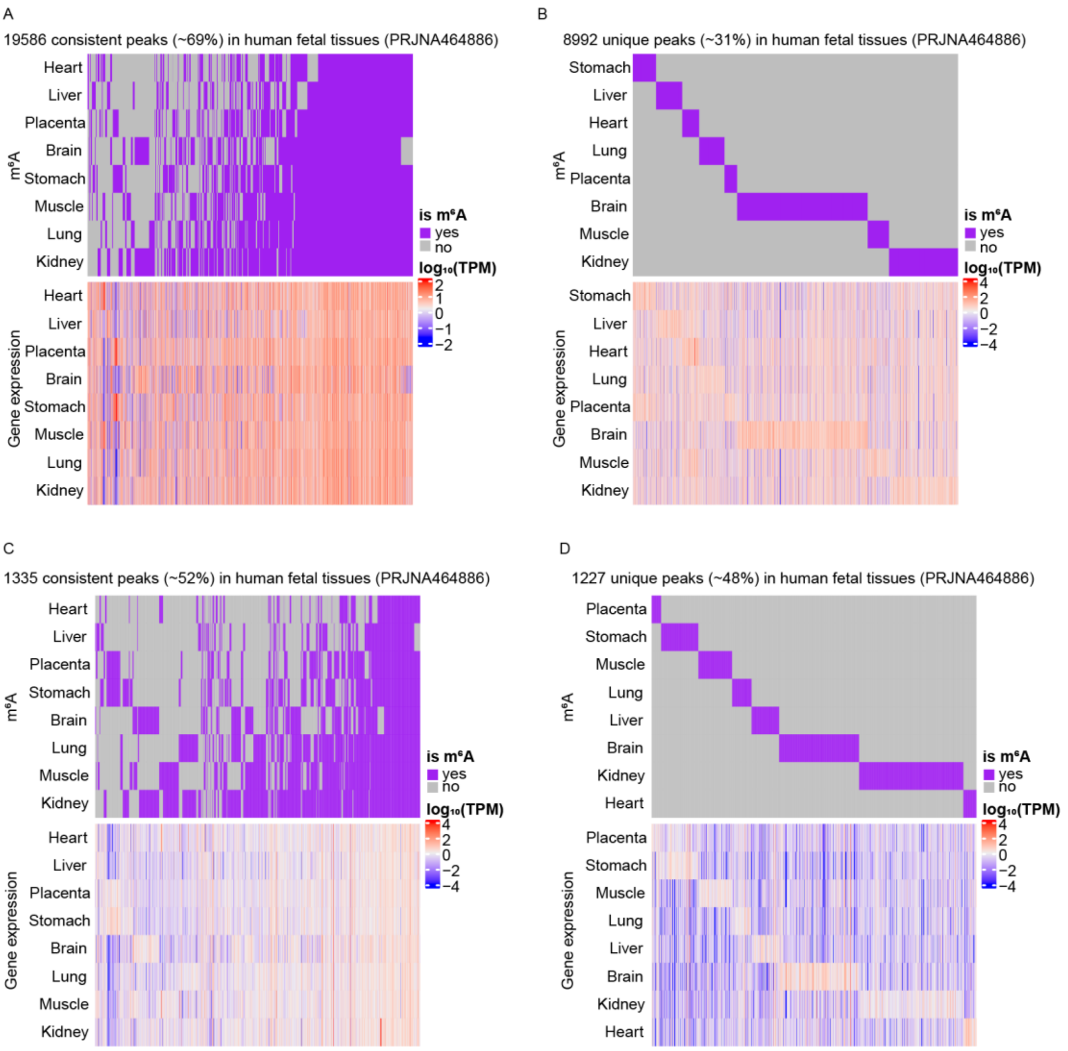


Supplementary Figure 12 **mRNA and lncRNA m6A modification and gene expression profiles in eight different human fetal tissues.** A, mRNA-consistent m6A sites and expression levels of genes marked by these sites. B, mRNA-unique m6A sites and expression levels of genes marked by these sites. C, lncRNA-consistent m6A sites and expression levels of genes marked by these sites. D, lncRNA-unique m6A sites and expression levels of genes marked by these sites. Rows denote different tissues, and columns denote different m6A sites (upper panel) and genes marked by these sites (lower panel).

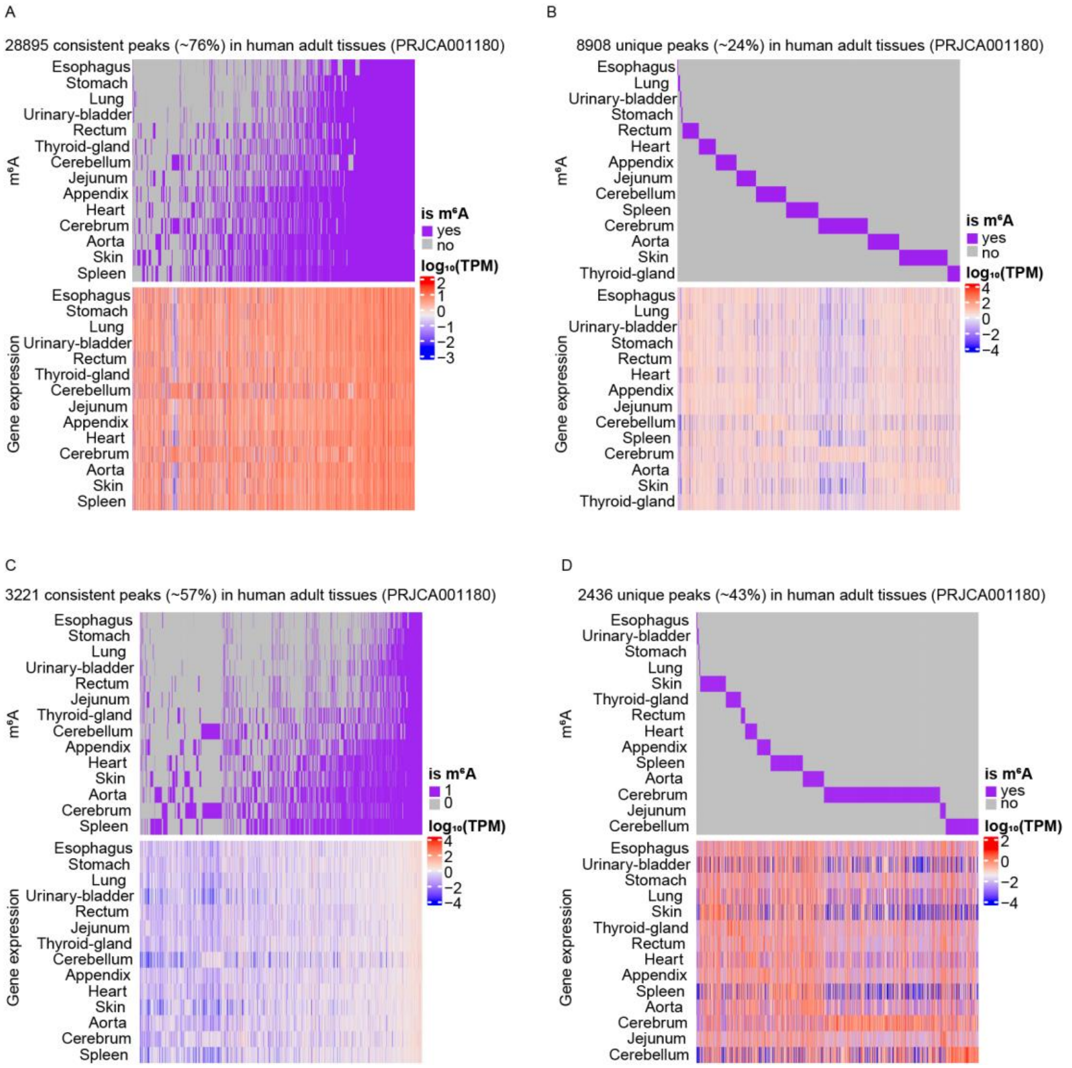


Supplementary Figure 13 **mRNA and lncRNA m6A modification and gene expression profiles in fourteen different adult tissues.** A, mRNA-consistent m6A sites and expression levels of genes marked by these sites. B, mRNA-unique m6A sites and expression levels of genes marked by these sites. C, lncRNA-consistent m6A sites and expression levels of genes marked by these sites. D, lncRNA-unique m6A sites and expression levels of genes marked by these sites. Rows denote different tissues, and columns denote different m6A sites (upper panel) and genes marked by these sites (lower panel).

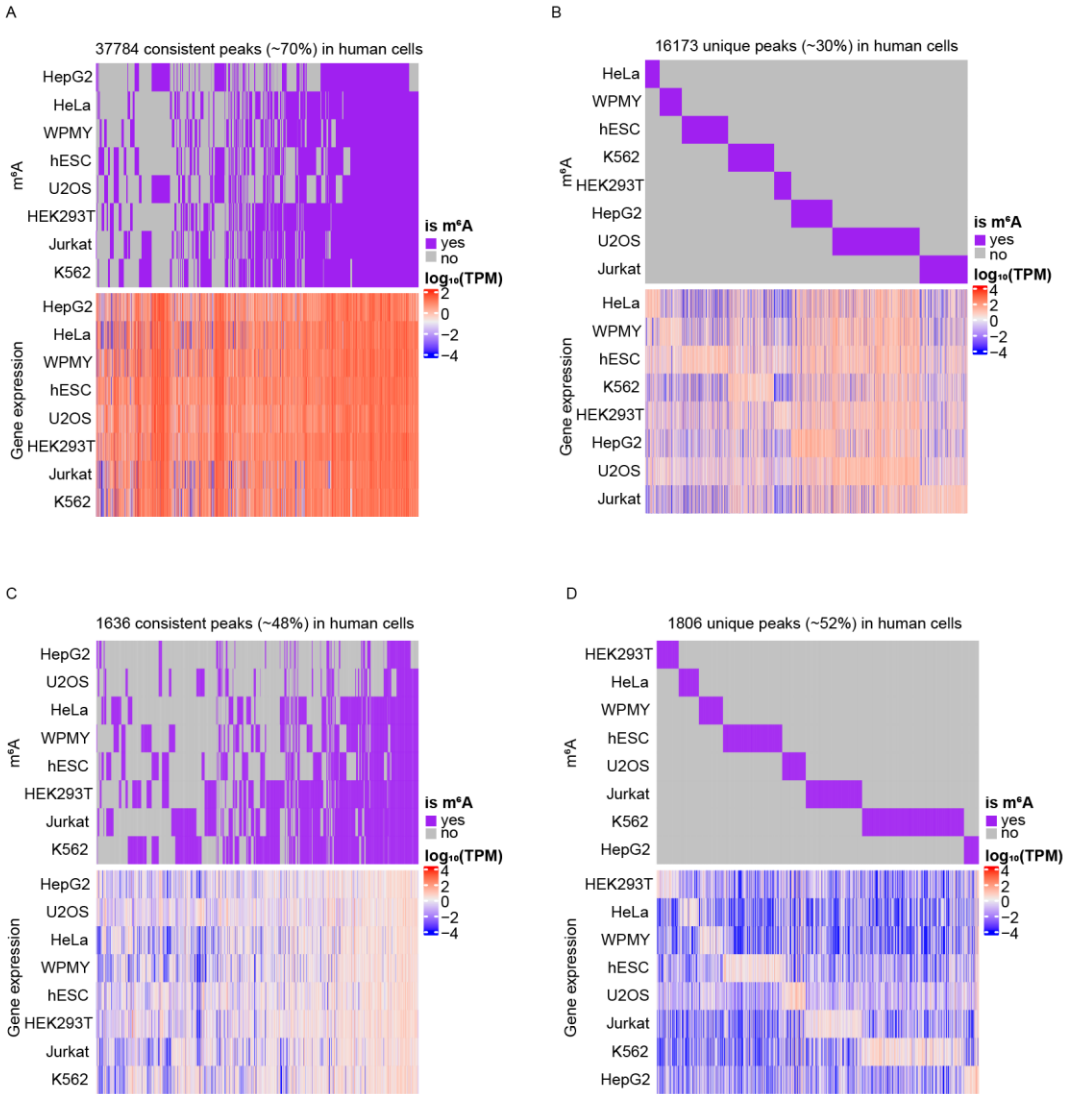


Supplementary Figure 14 **mRNA and lncRNA m6A modification and gene expression profiles in eight different human cell lines.** A, mRNA-consistent m6A sites and expression levels of genes marked by these sites. B, mRNA-unique m6A sites and expression levels of genes marked by these sites. C, lncRNA-consistent m6A sites and expression levels of genes marked by these sites. D, lncRNA-unique m6A sites and expression levels of genes marked by these sites. Rows denote different tissues, and columns denote different m6A sites (upper panel) and genes marked by these sites (lower panel).

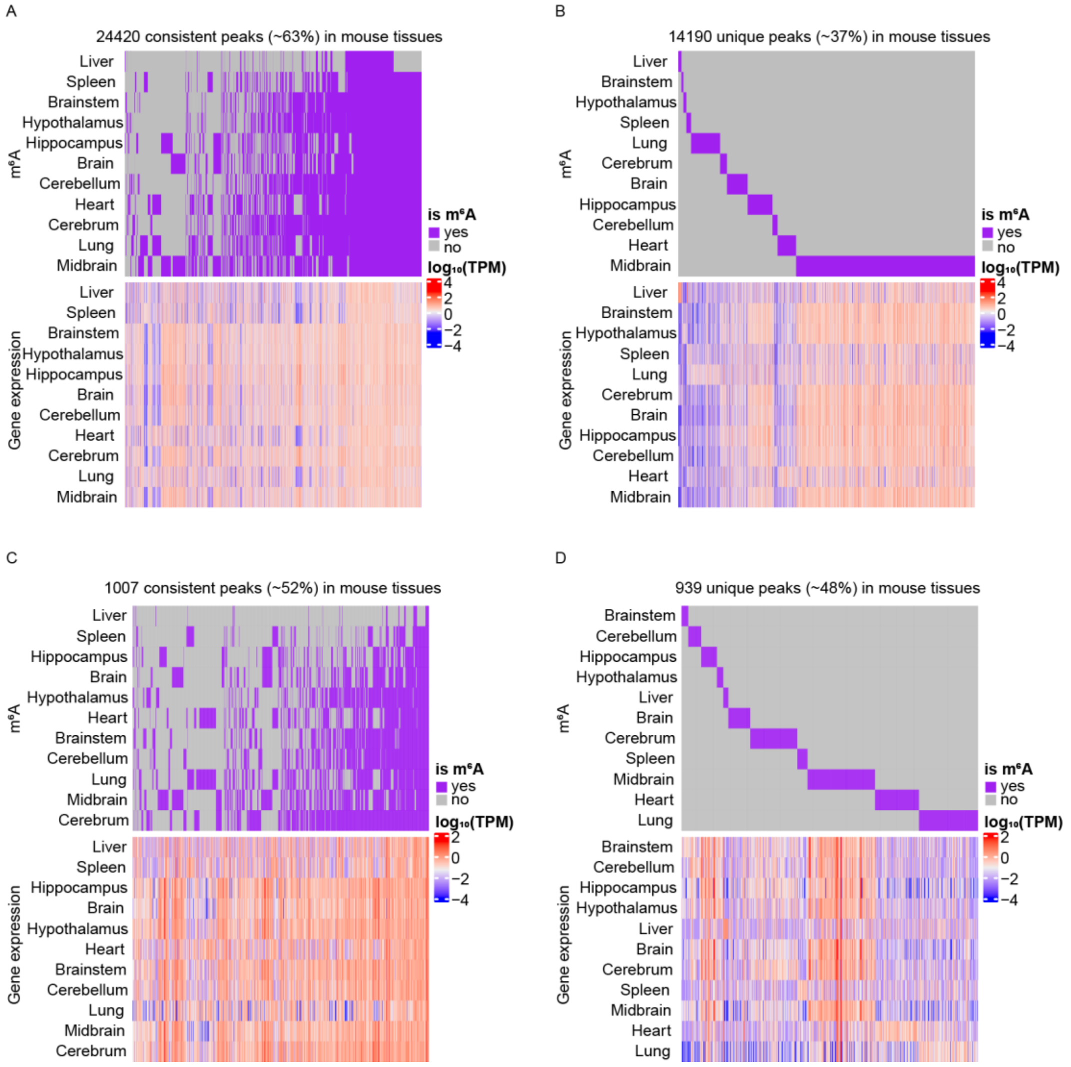


Supplementary Figure 15 **mRNA and lncRNA m6A modification and gene expression profiles in eleven different mouse tissues.** A, mRNA-consistent m6A sites and expression levels of genes marked by these sites. B, mRNA-unique m6A sites and expression levels of genes marked by these sites. C, lncRNA-consistent m6A sites and expression levels of genes marked by these sites. D, lncRNA-unique m6A sites and expression levels of genes marked by these sites. Rows denote different tissues, and columns denote different m6A sites (upper panel) and genes marked by these sites (lower panel).

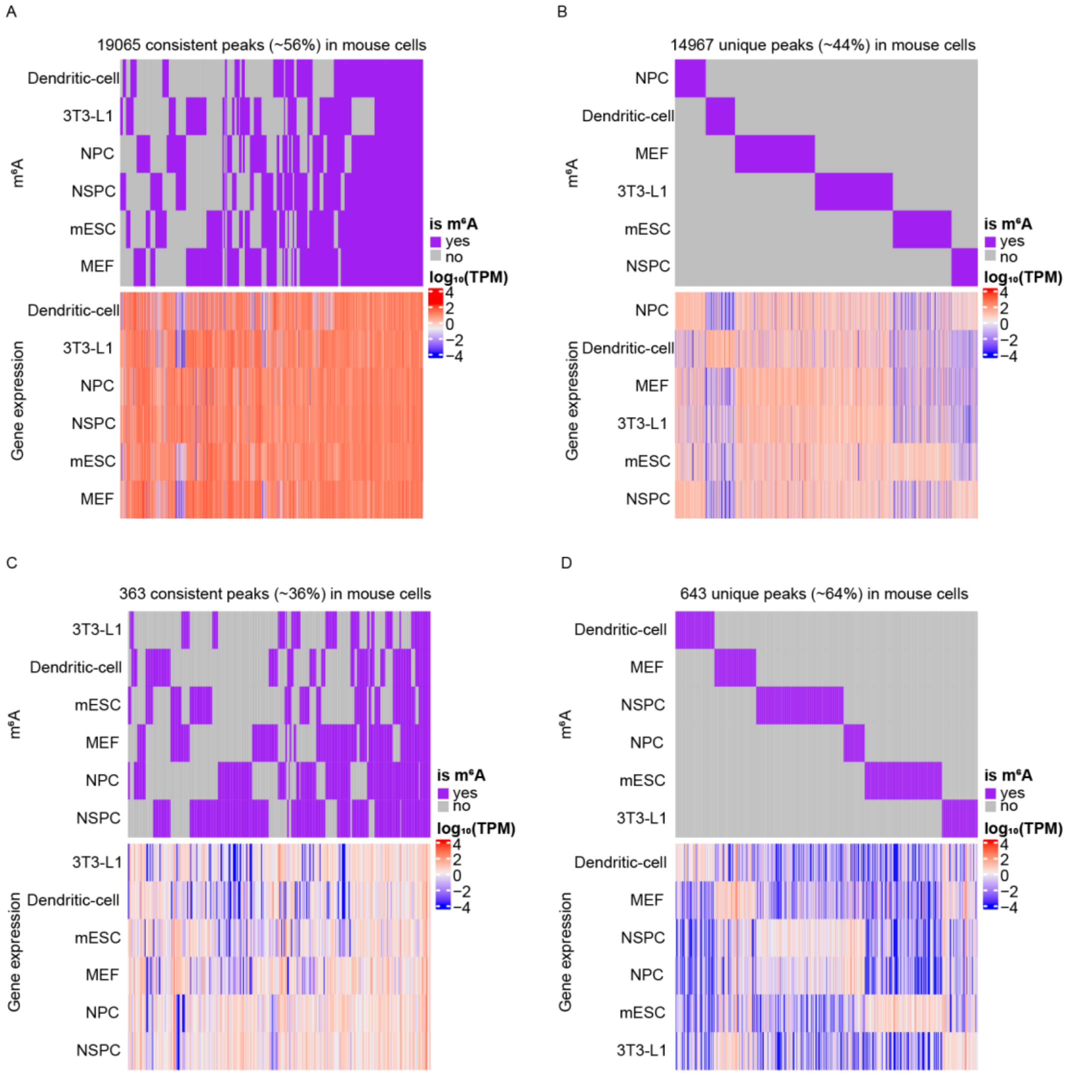


Supplementary Figure 16 **mRNA and lncRNA m6A modification and gene expression profiles in six different mouse cell lines.** A, mRNA-consistent m6A sites and expression levels of genes marked by these sites. B, mRNA-unique m6A sites and expression levels of genes marked by these sites. C, lncRNA-consistent m6A sites and expression levels of genes marked by these sites. D, lncRNA-unique m6A sites and expression levels of genes marked by these sites. Rows denote different tissues, and columns denote different m6A sites (upper panel) and genes marked by these sites (lower panel).

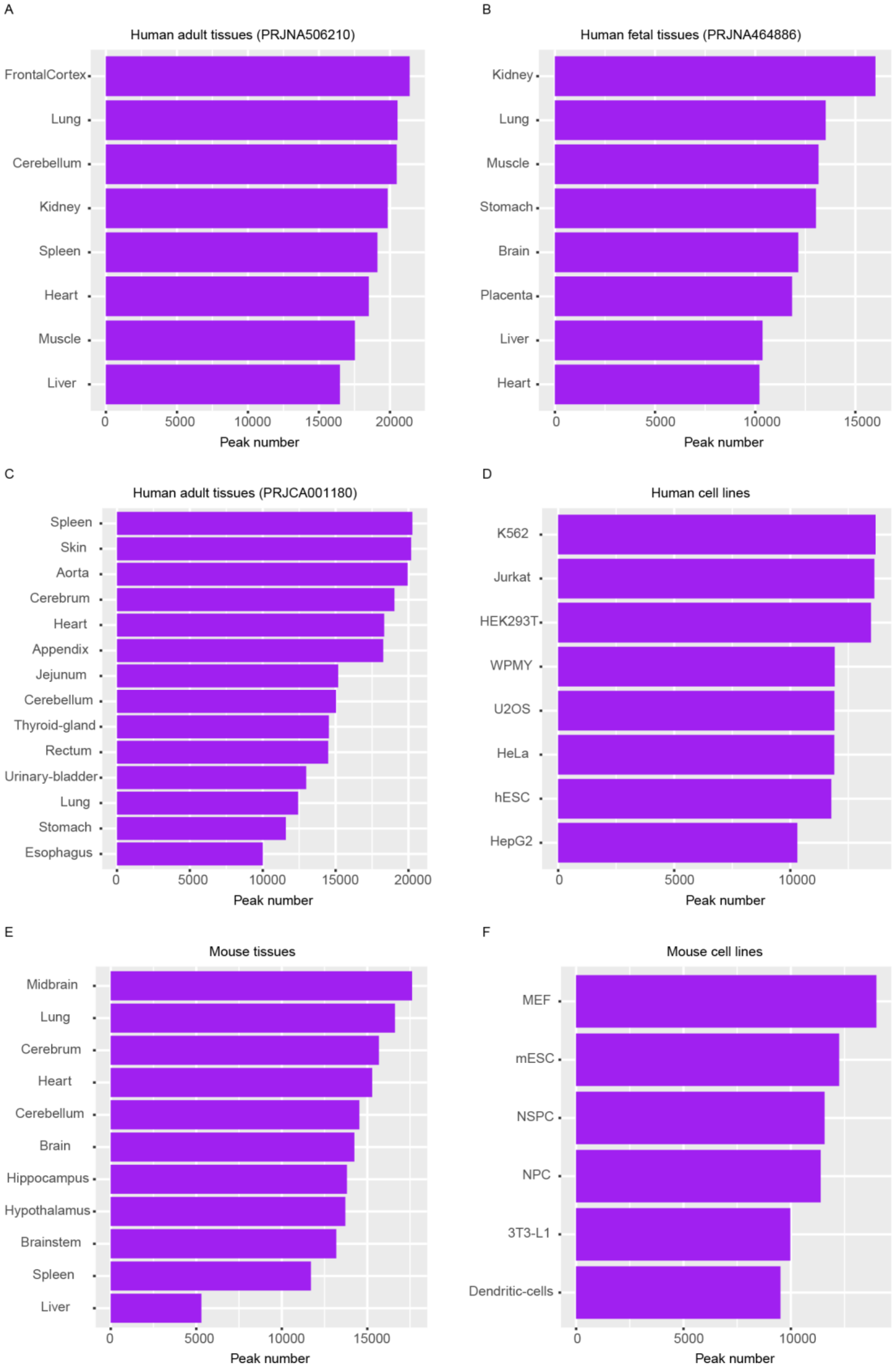


Supplementary Figure 17 **Number of mRNA-consistent m6A sites in different tissues and cell lines.** A-F, Number of mRNA-consistent m6A sites in eight adult tissues (A), eight human fetal tissues (B), fourteen adult tissues (C), eight human cell lines (D), eleven mouse tissues (E), and six mouse cell lines (F).

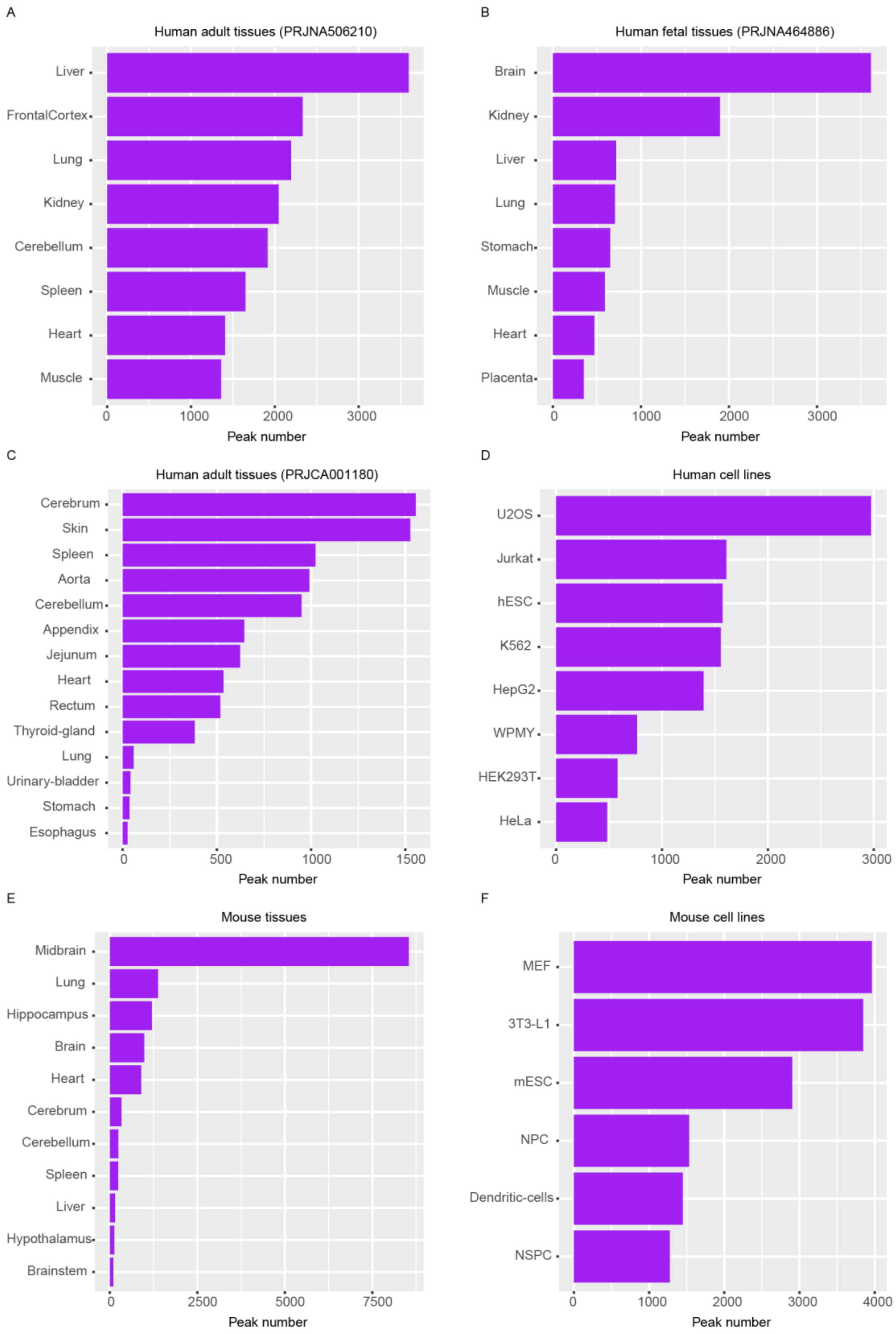


Supplementary Figure 18 **Number of mRNA-unique m6A sites in different tissues and cell lines.** A-F, Number of mRNA-unique m6A sites in eight adult tissues (A), eight human fetal tissues (B), fourteen adult tissues (C), eight human cell lines (D), eleven mouse tissues (E), and six mouse cell lines (F).

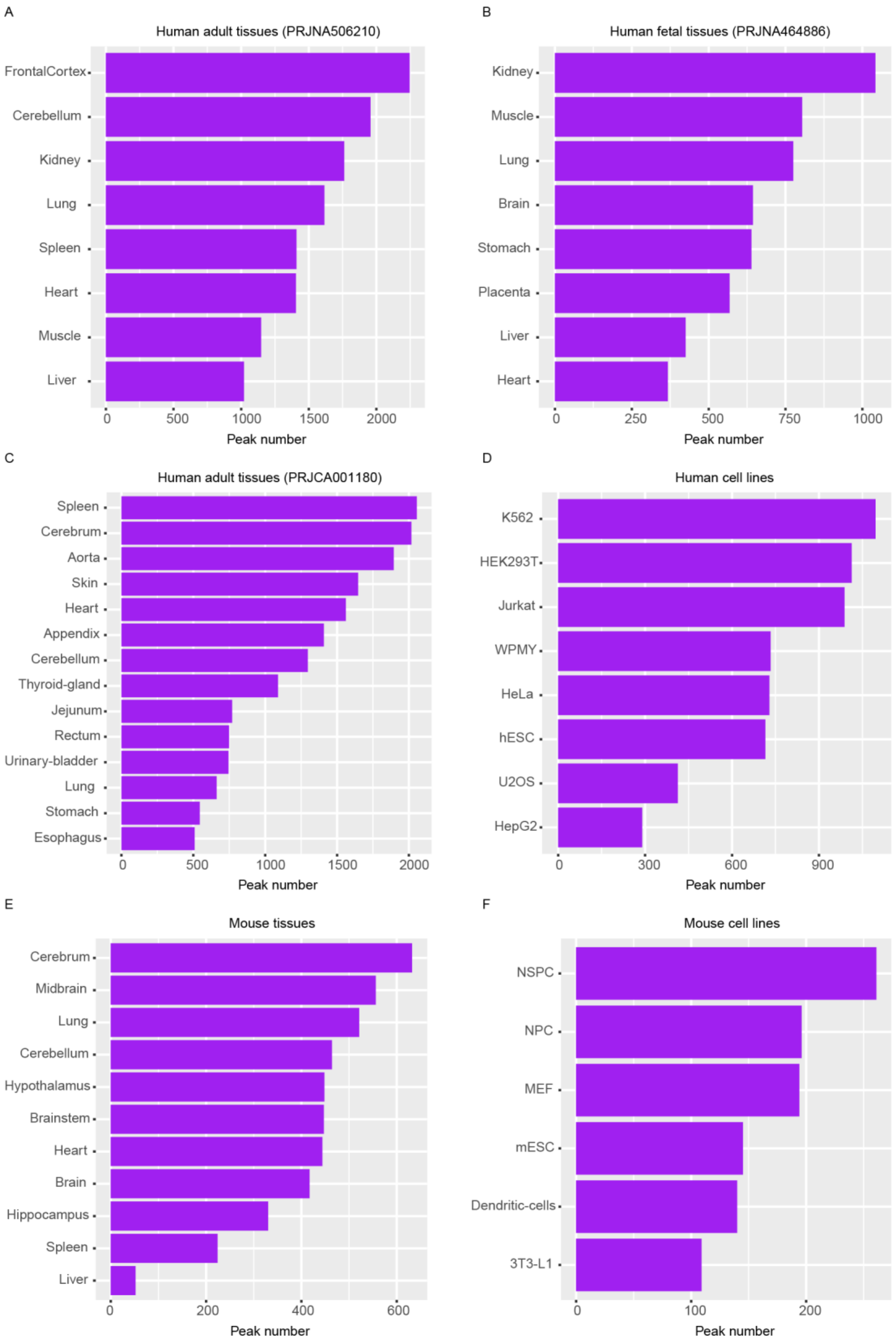


Supplementary Figure 19 **Number of lncRNA-consistent m6A sites in different tissues and cell lines.** A-F, Number of lncRNA-consistent m6A sites in eight adult tissues (A), eight human fetal tissues (B), fourteen adult tissues (C), eight human cell lines (D), eleven mouse tissues (E), and six mouse cell lines (F).

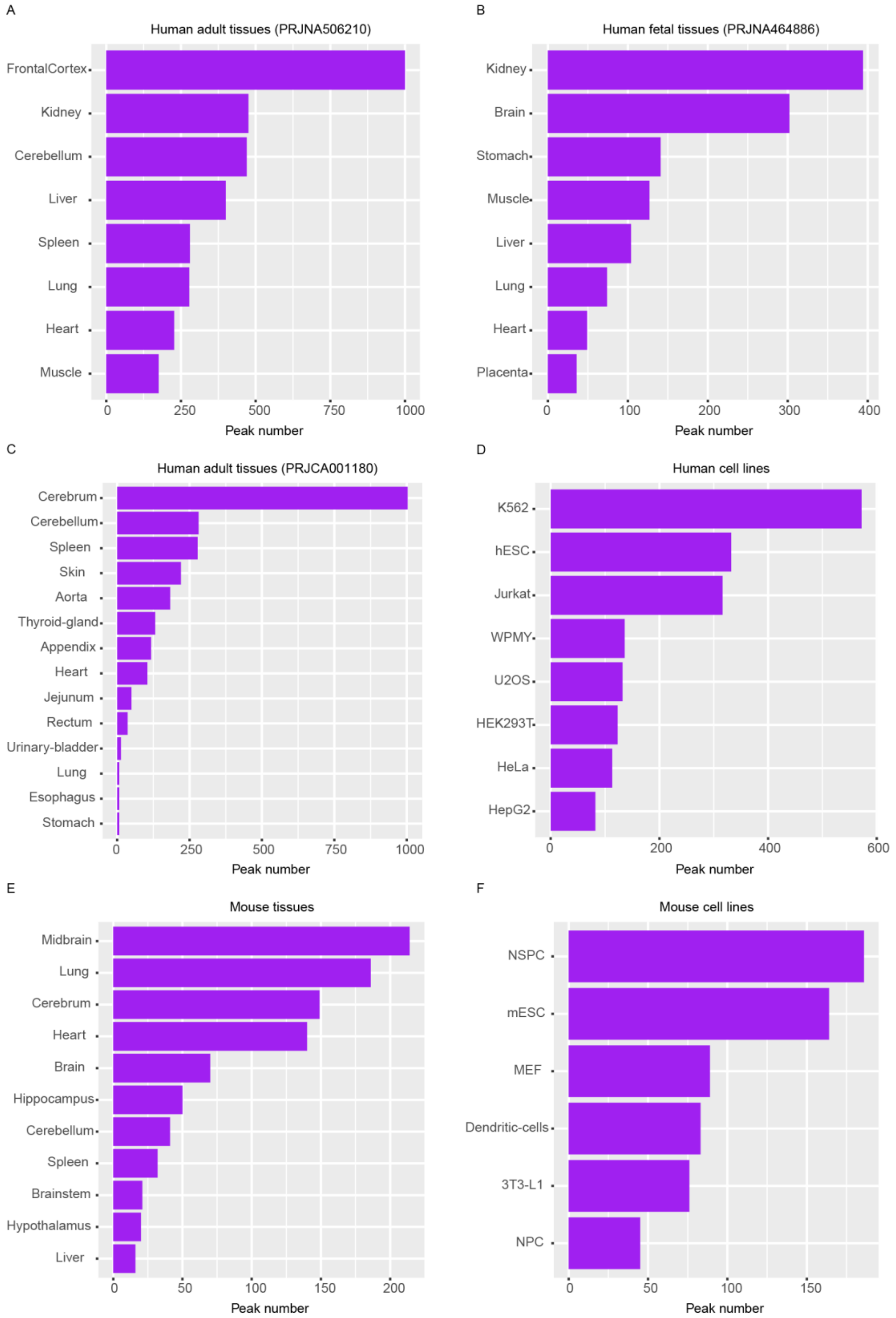


Supplementary Figure 20 **Number of lncRNA-unique m6A sites in different tissues and cell lines.** A-F, Number of lncRNA-unique m6A sites in eight adult tissues (A), eight human fetal tissues (B), fourteen adult tissues (C), eight human cell lines (D), eleven mouse tissues (E), and six mouse cell lines (F).

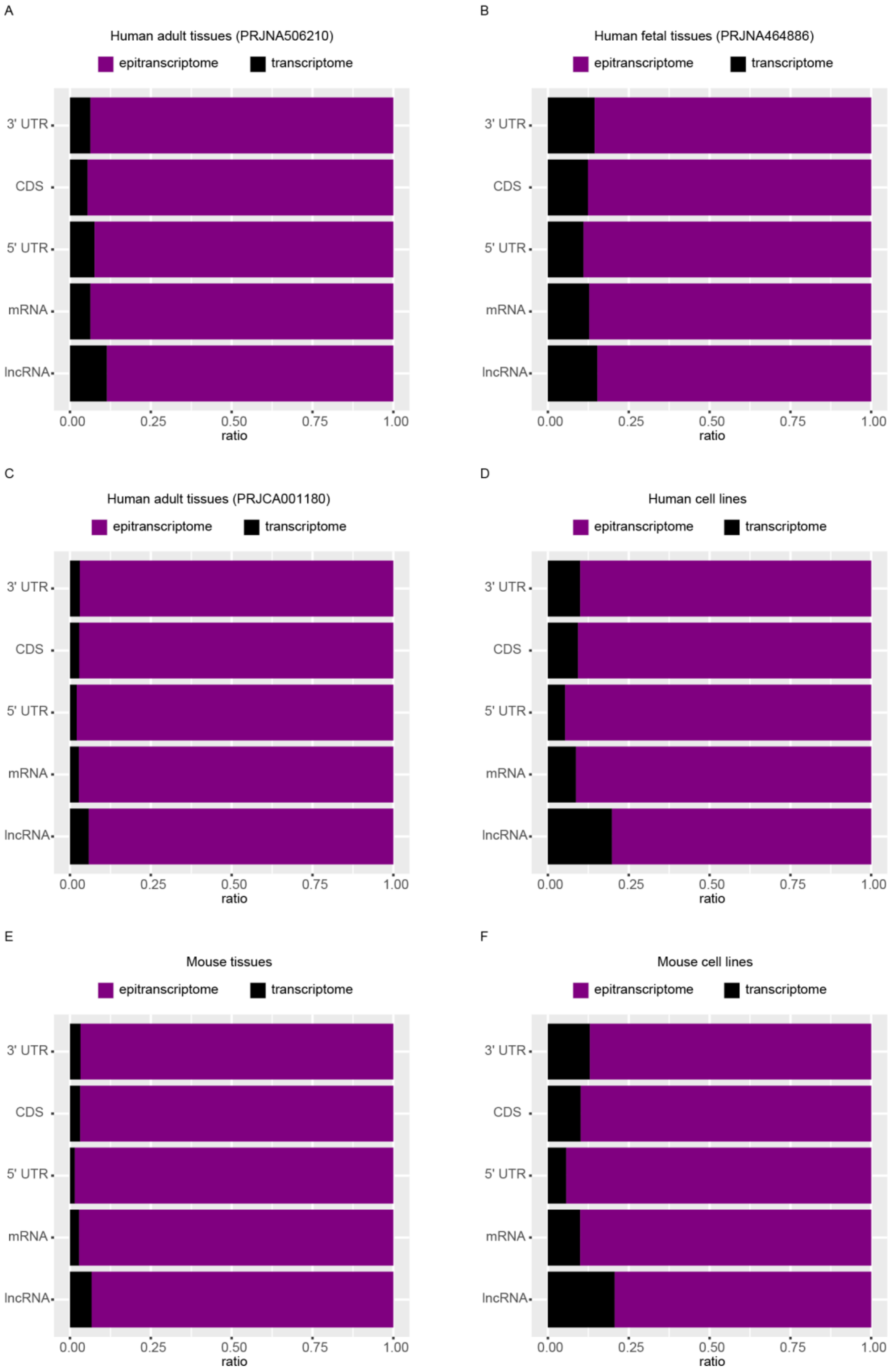


Supplementary Figure 21 **The proportion of transcriptome-specific and epi-transcriptome-specific m6A sites in mRNA, lncRNA, mRNA 5'-UTR, CDS, and 3'-UTR.** A-F, The proportion of transcriptome-specific m6A sites and epi-transcriptomes-specific m6A sites in eight adult tissues (A), eight human fetal tissues (B), fourteen adult tissues

(C), eight human cell lines (D), eleven mouse tissues (E), and six mouse cell lines (F).

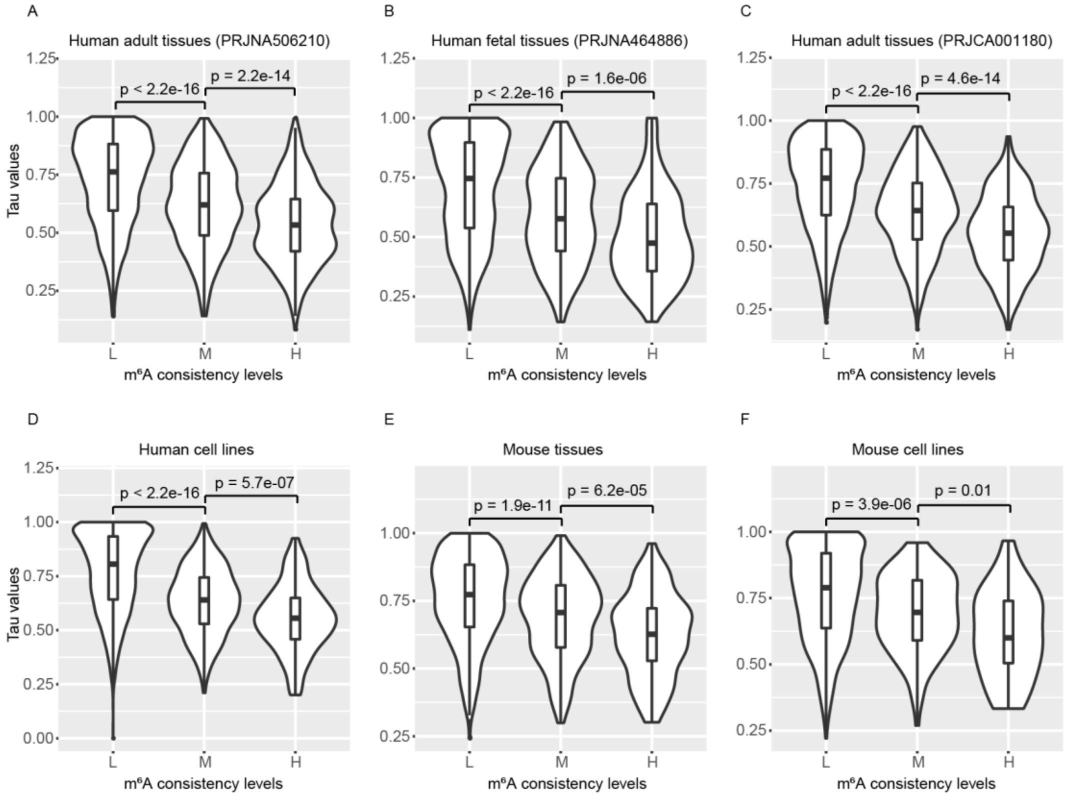


Supplementary Figure 22 **lncRNA m6A site consistency levels positively correlated with gene expression homeostasis.** A, Relative expression stability of m6A-modified genes (L, M, and H sets containing 3037, 706, and 468 lncRNA genes, respectively) among eight different adult tissues. B, Relative expression stability of m6A-modified genes (L, M, and H sets containing 1378, 345, and 184 lncRNA genes, respectively) among eight different human fetal tissues. C, Relative expression stability of m6A-modified genes (L, M, and H sets containing 2740, 604, and 327 lncRNA genes, respectively) among fourteen different adult tissues. D, Relative expression stability of m6A-modified genes (L, M, and H sets containing 1788, 434, and 140 lncRNA genes, respectively) among eight different human cell lines. E, Relative expression stability of m6A-modified genes (L, M, and H sets containing 1074, 290, and 152 lncRNA genes, respectively) among eleven different mouse tissues. F, Relative expression stability of m6A-modified genes (L, M, and H sets containing 618, 120, and 33 lncRNA genes, respectively) among six different mouse cell lines. Significance was evaluated by the two-sided Mann-Whitney test.

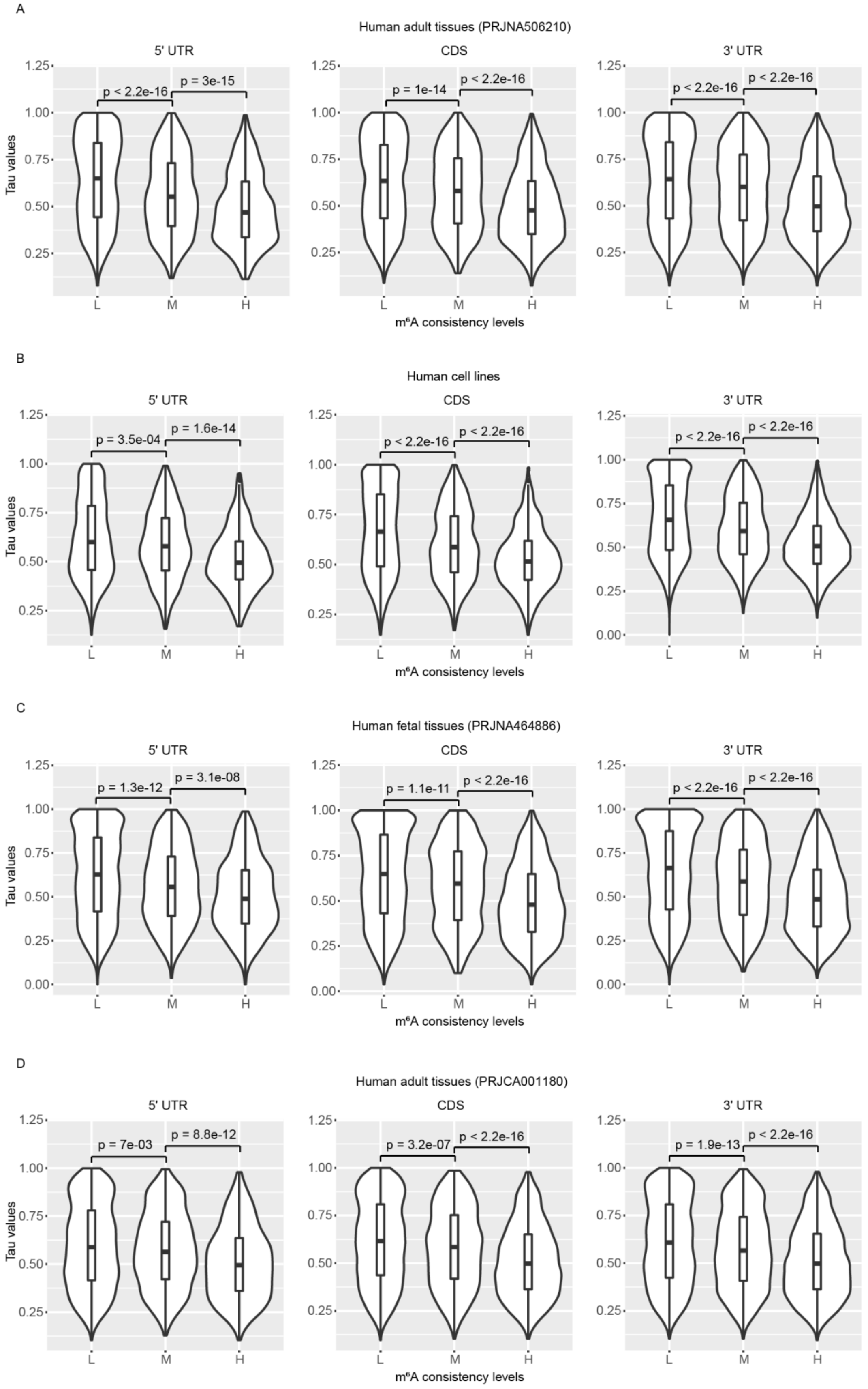


Supplementary Figure 23 **Consistency levels of m6A sites in the mRNA 5’-UTR, CDS, and 3’-UTR positively correlated with gene expression homeostasis.** A, Relative expression stability of m6A-modified genes (for 5’-UTR, L, M, and H sets containing 5124,

1257, and 969 mRNA genes, respectively; for CDS, L, M, and H sets containing 5561, 1746, and 1925 mRNA genes, respectively; for 3’-UTR, L, M, and H sets containing 8060, 3894, and 5916 mRNA genes, respectively) among eight different adult tissues. B, Relative expression stability of m6A-modified genes (for 5’-UTR, L, M, and H sets containing 3699, 780, and 483 mRNA genes, respectively; for CDS, L, M, and H sets containing 4199, 1536, and 1235 mRNA genes, respectively; for 3’-UTR, L, M, and H sets containing 6276, 3253, and 3468 mRNA genes, respectively) among eight different human cell lines. C, Relative expression stability of m6A-modified genes (for 5’-UTR, L, M, and H sets containing 3522, 1005, and 854 mRNA genes, respectively; for CDS, L, M, and H sets containing 3562, 1398, and 1783 mRNA genes, respectively; for 3’-UTR, L, M, and H sets containing 4744, 2465, and 3742 mRNA genes, respectively) among eight different human fetal tissues. D, Relative expression stability of m6A-modified genes (for 5’-UTR, L, M, and H sets containing 3219, 1017, and 841 mRNA genes, respectively; for CDS, L, M, and H sets containing 4615, 1603, and 2143 mRNA genes, respectively; for 3’-UTR, L, M, and H sets containing 6916, 3275, and 5311 mRNA genes, respectively) among fourteen different adult tissues. Significance was evaluated by the two-sided Mann-Whitney test.

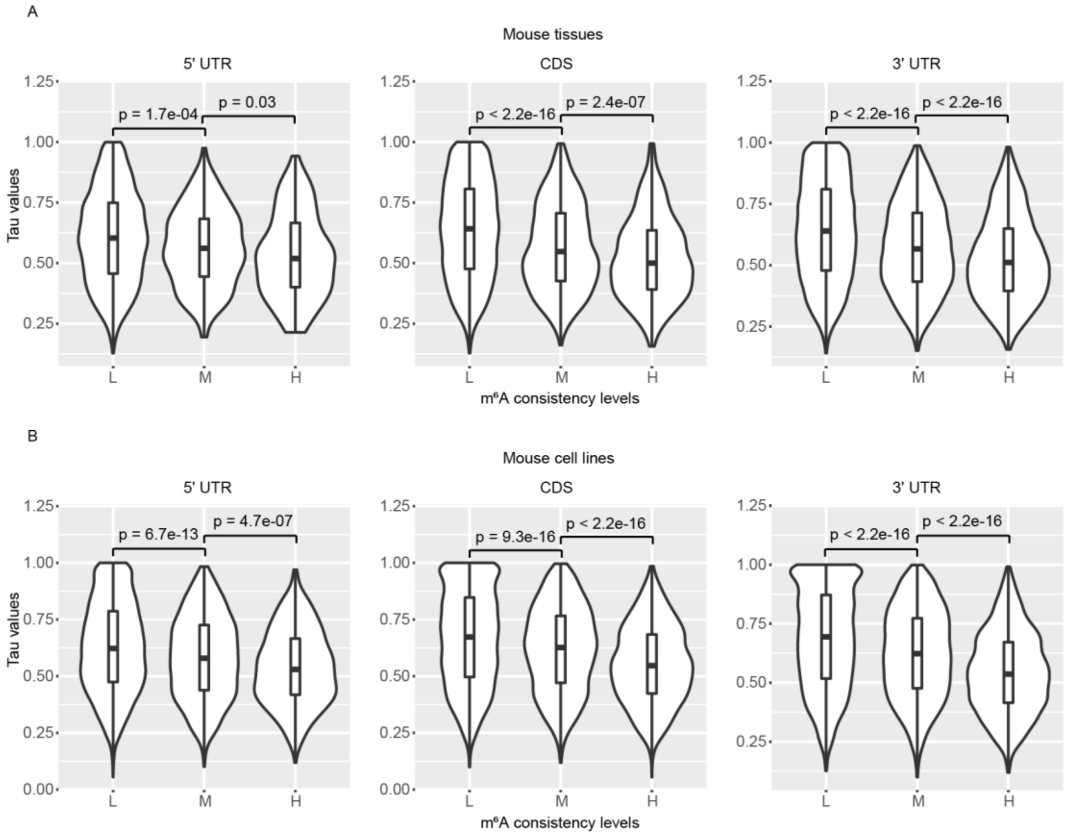


Supplementary Figure 24 **Consistency levels of m6A sites in the mRNA 5'-UTR, CDS, and 3'-UTR positively correlated with gene expression homeostasis.** A, Relative expression stability of m6A-modified genes (for 5'-UTR, L, M, and H sets containing 4844, 325, and 144 mRNA genes, respectively; for CDS, L, M, and H sets containing 5923, 1258, and 756 mRNA genes, respectively; for 3'-UTR, L, M, and H sets containing 9445, 2776, and 2273 mRNA genes, respectively) among eleven different mouse tissues. B, Relative expression stability of m6A-modified genes (for 5'-UTR, L, M, and H sets containing 5185, 1432, and 656 mRNA genes, respectively; for CDS, L, M, and H sets containing 4723, 1525, and 1376 mRNA genes, respectively; for 3'-UTR, L, M, and H sets containing 5902, 2801, and 3318 mRNA genes, respectively) among six different mouse cell lines. Significance was evaluated by the two-sided Mann-Whitney test.

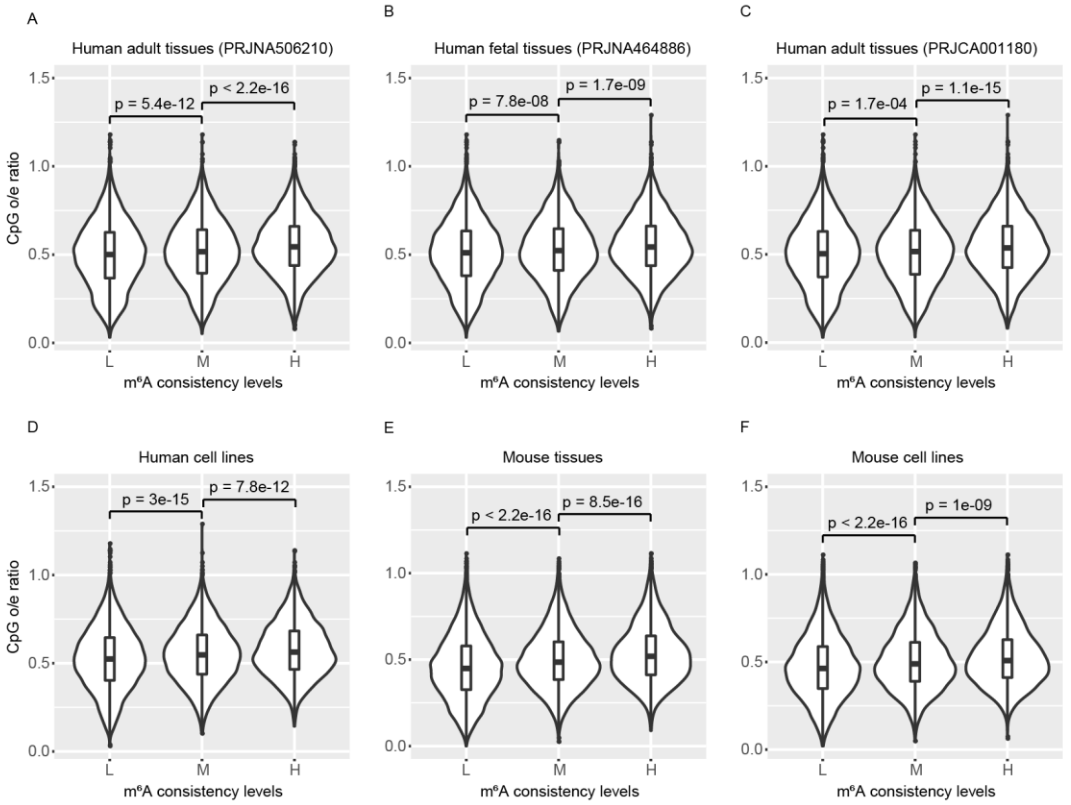


Supplementary Figure 25 **mRNA m6A consistency levels positively correlated with the CpG density of the gene promoter region**. A, CpG density of promoter regions of m6A-modified genes (L, M, and H sets containing 12228, 5824, and 7130 mRNA genes, respectively) among eight different adult tissues. B, CpG density of promoter regions of m6A-modified genes (L, M, and H sets containing 8735, 4242, and 5279 mRNA genes, respectively) among eight different human fetal tissues. C, CpG density of promoter regions of m6A-modified genes (L, M, and H sets containing 10470, 5104, and 6842 mRNA genes, respectively) among fourteen different adult tissues. D, CpG density of promoter regions of m6A-modified genes (L, M, and H sets containing 10031, 4769, and 4442 mRNA genes, respectively) among eight different human cell lines. E, CpG density of promoter regions of m6A-modified genes (L, M, and H sets containing 12906, 3913, and 2913 mRNA genes, respectively) among eleven different mouse tissues. F, CpG density of promoter regions of m6A-modified genes (L, M, and H sets containing 10653, 4754, and 4449 mRNA genes, respectively) among six different mouse cell lines. Significance was evaluated by the two-sided Mann-Whitney test.

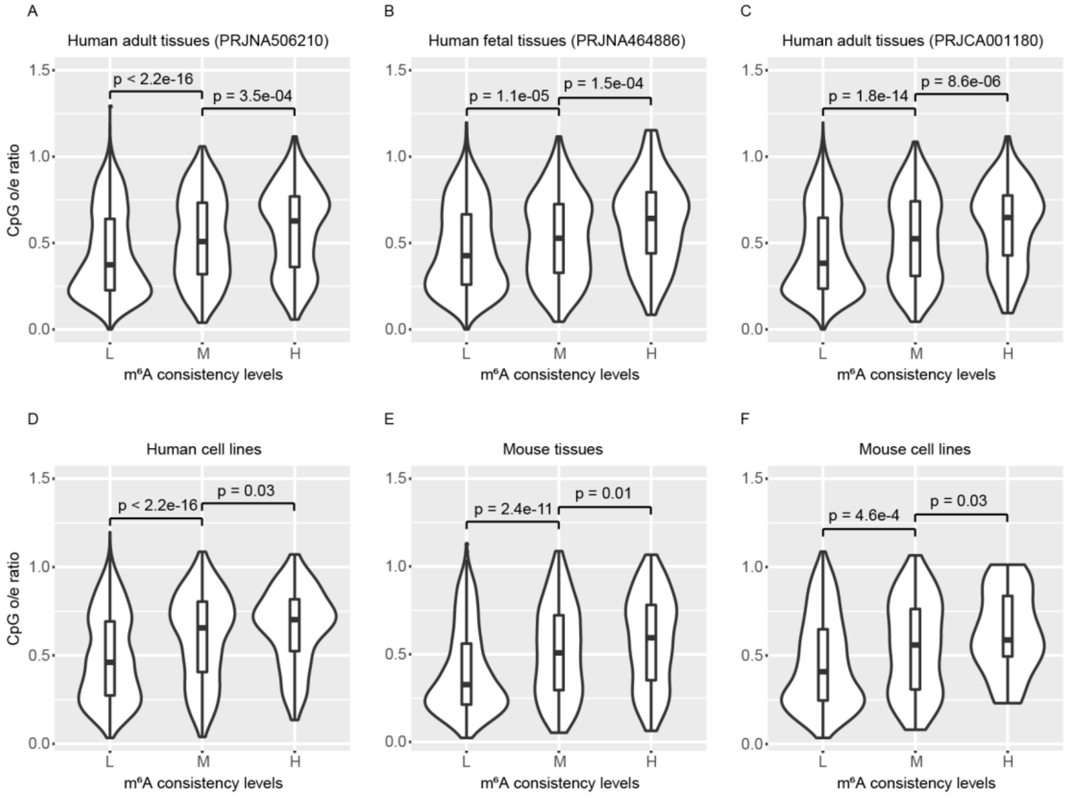


Supplementary Figure 26 **lncRNA m6A consistency levels positively correlated with the CpG density of the lncRNA gene promoter region.** A, CpG density of promoter regions of m6A-modified genes (L, M, and H sets containing 3037, 706, and 468 lncRNA genes, respectively) among eight different adult tissues. B, CpG density of promoter regions of m6A-modified genes (L, M, and H sets containing 1378, 345, and 184 lncRNA genes, respectively) among eight different human fetal tissues. C, CpG density of promoter regions of m6A-modified genes (L, M, and H sets containing 2740, 604, and 327 lncRNA genes, respectively) among fourteen different adult tissues. D, CpG density of promoter regions of m6A-modified genes (L, M, and H sets containing 1788, 434, and 140 lncRNA genes, respectively) among eight different human cell lines. E, CpG density of promoter regions of m6A-modified genes (L, M, and H sets containing 1074, 290, and 152 lncRNA genes, respectively) among eleven different mouse tissues. F, CpG density of promoter regions of m6A-modified genes (L, M, and H sets containing 618, 120, and 33 lncRNA genes, respectively) among six different mouse cell lines. Significance was evaluated by the two-sided Mann-Whitney test.

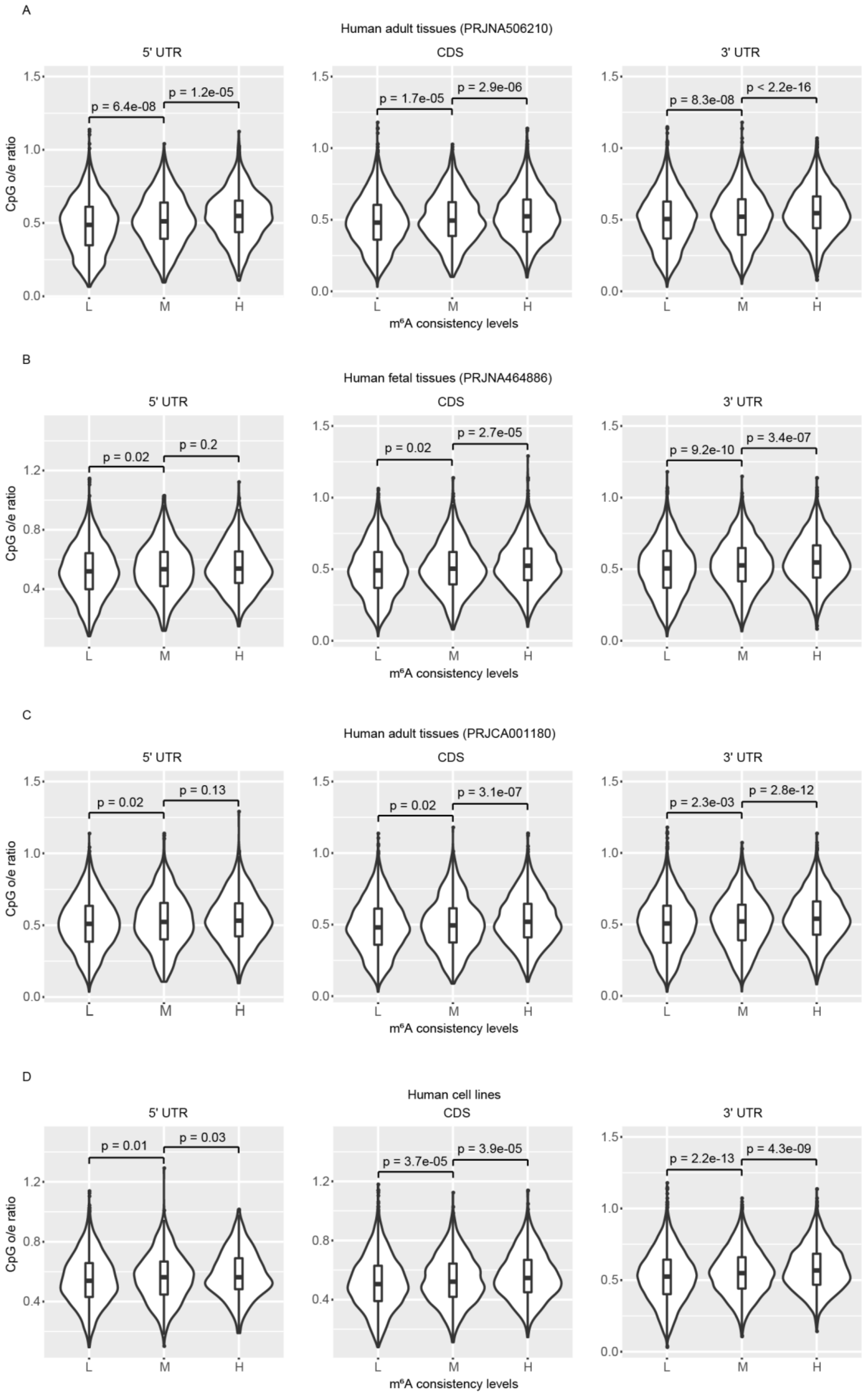


Supplementary Figure 27 **Consistency levels of m6A sites in the mRNA 5'-UTR, CDS, and 3'-UTR positively correlated with the CpG density of the mRNA gene promoter**

**region.** A, CpG density of promoter regions of m6A-modified genes (for 5'-UTR, L, M, and H sets containing 5124, 1257, and 969 mRNA genes, respectively; for CDS, L, M, and H sets containing 5561, 1746, and 1925 mRNA genes, respectively; for 3'-UTR, L, M, and H sets containing 8060, 3894, and 5916 mRNA genes, respectively) among eight different adult tissues. B, CpG density of promoter regions of m6A-modified genes (for 5'-UTR, L, M, and H sets containing 3522, 1005, and 854 mRNA genes, respectively; for CDS, L, M, and H sets containing 3562, 1398, and 1783 mRNA genes, respectively; for 3'-UTR, L, M, and H sets containing 4744, 2465, and 3742 mRNA genes, respectively) among eight different human fetal tissues. C, CpG density of promoter regions of m6A-modified genes (for 5'-UTR, L, M, and H sets containing 3219, 1017, and 841 mRNA genes, respectively; for CDS, L, M, and H sets containing 4615, 1603, and 2143 mRNA genes, respectively; for 3'-UTR, L, M, and H sets containing 6916, 3275, and 5311 mRNA genes, respectively) among fourteen different adult tissues. D, CpG density of promoter regions of m6A-modified genes (for 5'-UTR, L, M, and H sets containing 3699, 780, and 483 mRNA genes, respectively; for CDS, L, M, and H sets containing 4199, 1536, and 1235 mRNA genes, respectively; for 3'-UTR, L, M, and H sets containing 6276, 3253, and 3468 mRNA genes, respectively) among eight different human cell lines. Significance was evaluated by the two-sided Mann-Whitney test.

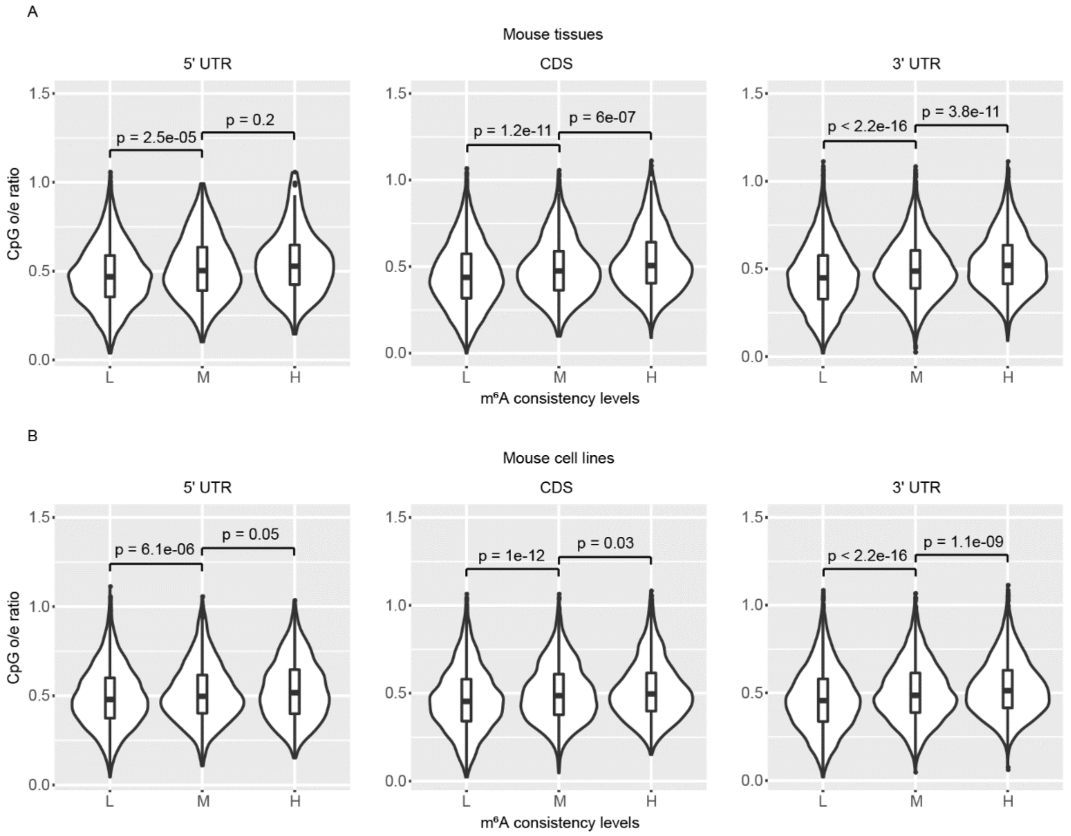


Supplementary Figure 28 **Consistency levels of m6A sites in the mRNA 5'-UTR, CDS, and 3'-UTR positively correlated with the CpG density of the mRNA gene promoter region.** A, CpG density of promoter regions of m6A-modified genes (for 5'-UTR, L, M, and H sets containing 4844, 325, and 144 mRNA genes, respectively; for CDS, L, M, and H sets containing 5923, 1258, and 756 mRNA genes, respectively; for 3'-UTR, L, M, and H sets containing 9445, 2776, and 2273 mRNA genes, respectively) among eleven different mouse tissues. B, CpG density of promoter regions of m6A-modified genes (for 5'-UTR, L, M, and H sets containing 5185, 1432, and 656 mRNA genes, respectively; for CDS, L, M, and H sets containing 4723, 1525, and 1376 mRNA genes, respectively; for 3'-UTR, L, M, and H sets containing 5902, 2801, and 3318 mRNA genes, respectively) among six different mouse cell lines. Significance was evaluated by the two-sided Mann-Whitney test.

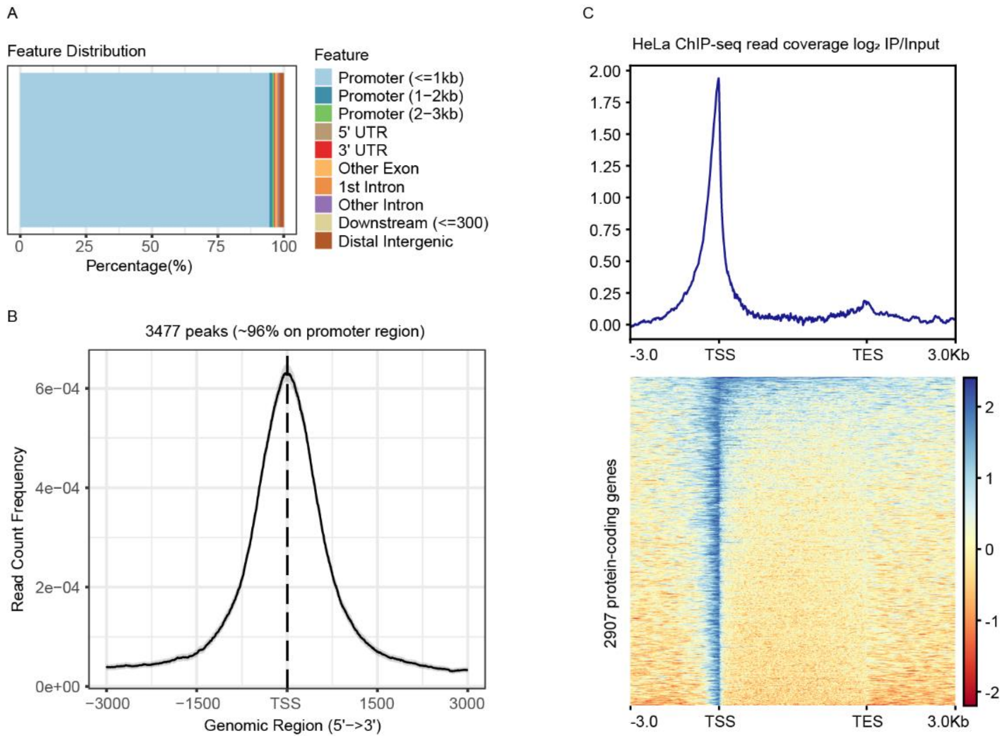


Supplementary Figure 29 **Distribution of METTL3 binding sites on the genome for HeLa.** A, Distribution of METTL3 binding sites in different genomic regions. B, The METTL3 binding sites were significantly enriched near the transcription start site (TSS). C, Distribution of METTL3 ChIP-seq read signal in 2907 protein-coding genes and their upstream and downstream 3Kb regions.

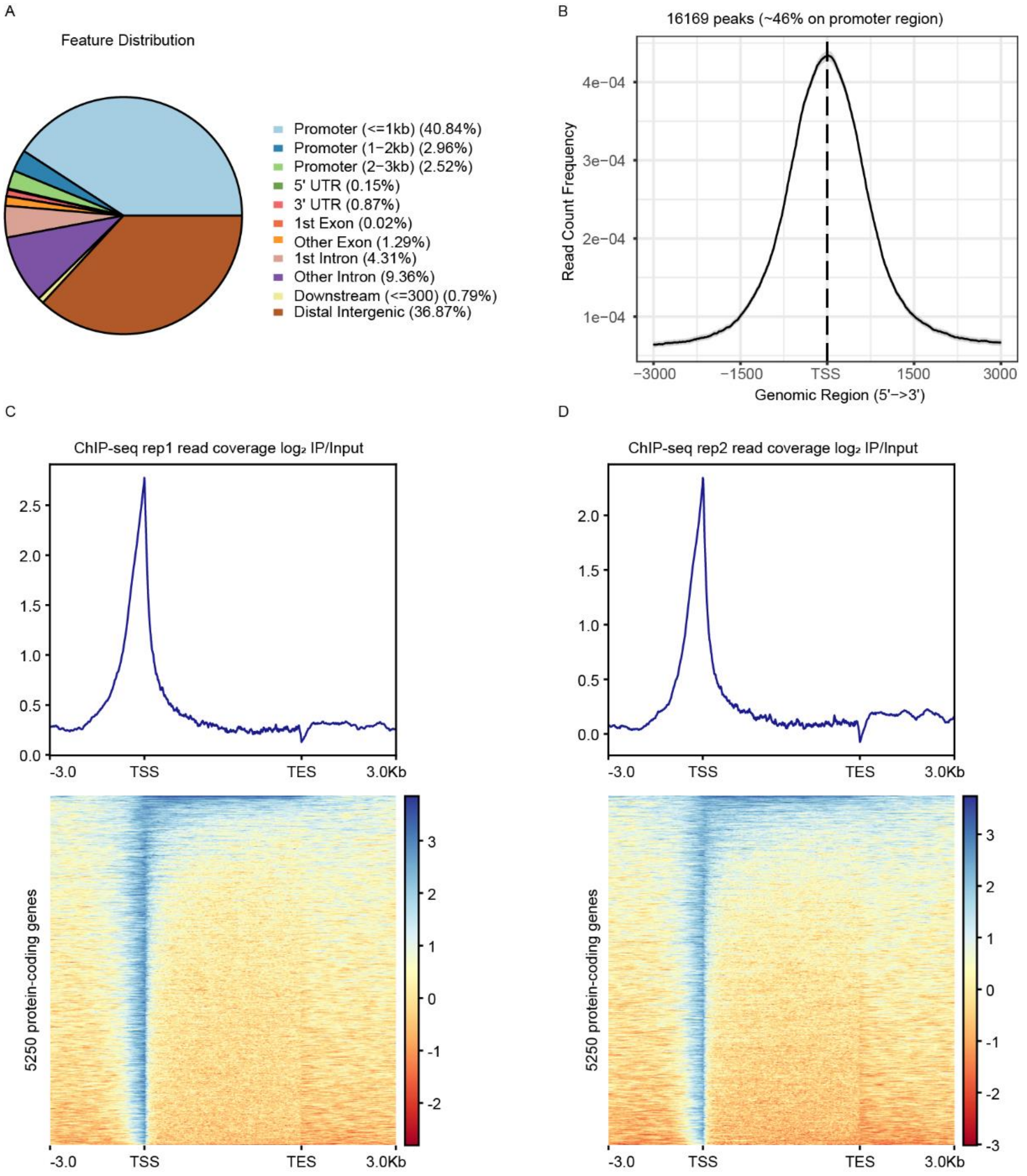


Supplementary Figure 30 **Distribution of METTL3 binding sites on the genome for mESC.** A, Distribution of METTL3 binding sites in different genomic regions. B, The METTL3 binding sites are significantly enriched near the transcription start site (TSS). C, D, Distribution of METTL3 ChIP-seq read signal in 5250 protein-coding genes and their upstream and downstream 3Kb regions.

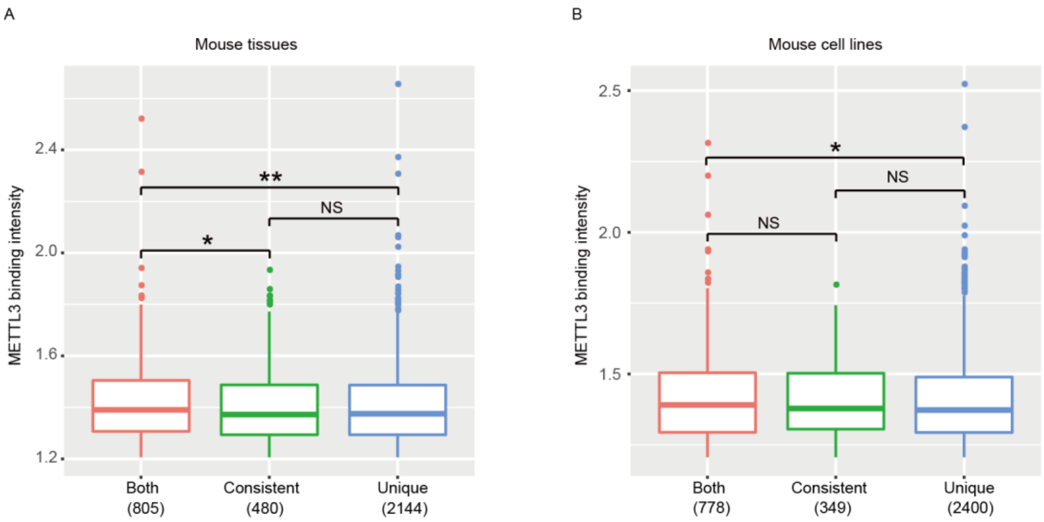


Supplementary Figure 31 **Binding intensity of METTL3 on three types of genes for mESC.** A, B, The METTL3-binding genes in mESC were classified into three categories, and METTL3 binding intensity was shown for mouse tissues (A) and mouse cell lines (B). Significance was evaluated by the two-sided Mann-Whitney test. An asterisk (*) denoted that the P value was between 0.01 and 0.05; Two asterisks (**) denoted that the P value was less than 0.01; Three asterisks (***) denoted that the P value was less than 0.001. NS denoted no significant difference. The numbers in parentheses denoted the number of genes.

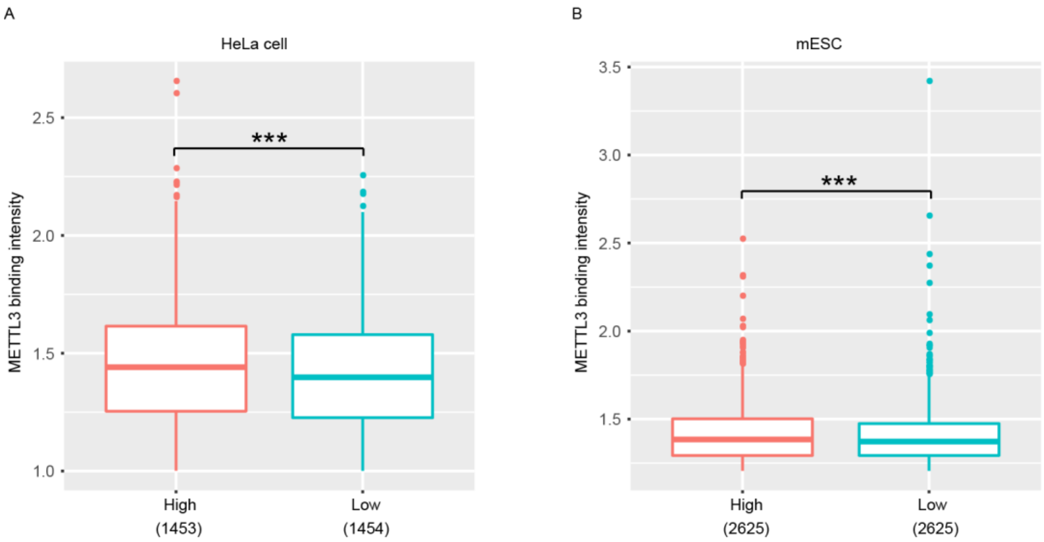


Supplementary Figure 32 **Binding intensity of METTL3 on High-CpG and Low-CpG gene promoter regions.** A, In HeLa, 2907 METTL3-binding genes were classified into two categories, and METTL3 binding intensity was shown. B, In mESC, 5250 METTL3-binding genes were classified into two categories, and METTL3 binding intensity was shown. Significance was evaluated by the two-sided Mann-Whitney test. An asterisk (*) denoted that the P value was between 0.01 and 0.05; Two asterisks (**) denoted that the P value was less than 0.01; Three asterisks (***) denoted that the P value was less than 0.001. NS denoted no significant difference. The numbers in parentheses denoted the number of genes.

Supplementary Table 1 m6A-seq data information for different human cell lines

| run accession | sample title | library layout | sample type | study accession | antibody info |
|---|---|---|---|---|---|
| SRR4310464 | GSC-11-rep1 | SINGLE | input | PRJNA345000 | SYSY |
| SRR4310465 | GSC-11-rep2 | SINGLE | input | PRJNA345000 | SYSY |
| SRR4310468 | GSC-11-rep1 | SINGLE | ip | PRJNA345000 | SYSY |
| SRR4310469 | GSC-11-rep2 | SINGLE | ip | PRJNA345000 | SYSY |
| SRR3066066 | Mono-mac6-rep1 | SINGLE | input | PRJNA307239 | SYSY |
| SRR3066067 | Mono-mac6-rep1 | SINGLE | ip | PRJNA307239 | SYSY |
| SRR3066068 | Mono-mac6-rep2 | SINGLE | input | PRJNA307239 | SYSY |
| SRR3066069 | Mono-mac6-rep2 | SINGLE | ip | PRJNA307239 | SYSY |
| SRR5128023 | U2OS-rep1 | SINGLE | input | PRJNA358676 | SYSY |
| SRR5128024 | U2OS-rep1 | SINGLE | ip | PRJNA358676 | SYSY |
| SRR5128025 | U2OS-rep2 | SINGLE | input | PRJNA358676 | SYSY |
| SRR5128026 | U2OS-rep2 | SINGLE | ip | PRJNA358676 | SYSY |
| SRR5128027 | U2OS-rep3 | SINGLE | input | PRJNA358676 | SYSY |
| SRR5128028 | U2OS-rep3 | SINGLE | ip | PRJNA358676 | SYSY |
| SRR494613 | HEK293T-rep1 | SINGLE | input | PRJNA141085 | SYSY |
| SRR494614 | HEK293T-rep1 | SINGLE | ip | PRJNA141085 | SYSY |
| SRR494615 | HEK293T-rep2 | SINGLE | input | PRJNA141085 | SYSY |
| SRR494616 | HEK293T-rep2 | SINGLE | ip | PRJNA141085 | SYSY |
| SRR494617 | HEK293T-rep3 | SINGLE | input | PRJNA141085 | NEB |
| SRR494618 | HEK293T-rep3 | SINGLE | ip | PRJNA14108 | NEB |

| | | | | | |
|---|---|---|---|---|---|
| | | | | 5 | |
| SRR847358 | HeLa-rep1 | SINGLE | input | PRJNA202057 | SYSY |
| SRR847359 | HeLa-rep2 | SINGLE | input | PRJNA202057 | SYSY |
| SRR847360 | HeLa-rep1 | SINGLE | ip | PRJNA202057 | SYSY |
| SRR847361 | HeLa-rep2 | SINGLE | ip | PRJNA202057 | SYSY |
| SRR847370 | HeLa-rep1 | SINGLE | input | PRJNA202057 | SYSY |
| SRR847371 | HeLa-rep2 | SINGLE | input | PRJNA202057 | SYSY |
| SRR847372 | HeLa-rep1 | SINGLE | ip | PRJNA202057 | SYSY |
| SRR847373 | HeLa-rep2 | SINGLE | ip | PRJNA202057 | SYSY |
| SRR1035213 | hESC-rep1 | SINGLE | input | PRJNA229454 | SYSY |
| SRR1035214 | hESC-rep1 | SINGLE | ip | PRJNA229454 | SYSY |
| SRR1035215 | hESC-rep2 | SINGLE | input | PRJNA229454 | SYSY |
| SRR1035216 | hESC-rep2 | SINGLE | ip | PRJNA229454 | SYSY |
| SRR1035221 | hESC-rep1 | SINGLE | input | PRJNA229454 | SYSY |
| SRR1035222 | hESC-rep1 | SINGLE | ip | PRJNA229454 | SYSY |
| SRR1035223 | hESC-rep2 | SINGLE | input | PRJNA229454 | SYSY |
| SRR1035224 | hESC-rep2 | SINGLE | ip | PRJNA229454 | SYSY |
| SRR1035217 | Endoderm-cells-rep1 | SINGLE | input | PRJNA229454 | SYSY |
| SRR1035218 | Endoderm-cells-rep1 | SINGLE | ip | PRJNA229454 | SYSY |
| SRR1035219 | Endoderm-cells-rep2 | SINGLE | input | PRJNA229454 | SYSY |
| SRR1035220 | Endoderm-cells-rep2 | SINGLE | ip | PRJNA229454 | SYSY |
| SRR5239086 | MOLM13-rep1 | SINGLE | input | PRJNA371833 | Millipore |
| SRR5239087 | MOLM13-rep1 | SINGLE | input | PRJNA37183 | Millipore |

| | | | | | |
|---|---|---|---|---|---|
| | | | | 3 | |
| SRR5239088 | MOLM13-rep2 | SINGLE | input | PRJNA371833 | Millipore |
| SRR5239089 | MOLM13-rep2 | SINGLE | input | PRJNA371833 | Millipore |
| SRR5239098 | MOLM13-rep1 | SINGLE | ip | PRJNA371833 | Millipore |
| SRR5239099 | MOLM13-rep1 | SINGLE | ip | PRJNA371833 | Millipore |
| SRR5239100 | MOLM13-rep2 | SINGLE | ip | PRJNA371833 | Millipore |
| SRR5239101 | MOLM13-rep2 | SINGLE | ip | PRJNA371833 | Millipore |
| SRR2648293 | MT4-rep1 | SINGLE | input | PRJNA298743 | SYSY |
| SRR2648294 | MT4-rep1 | SINGLE | ip | PRJNA298743 | SYSY |
| SRR2648296 | MT4-rep2 | SINGLE | input | PRJNA298743 | SYSY |
| SRR2648297 | MT4-rep2 | SINGLE | ip | PRJNA298743 | SYSY |
| SRR5931733 | HepG2-rep1 | SINGLE | input | PRJNA398233 | SYSY |
| SRR5931734 | HepG2-rep1 | SINGLE | ip | PRJNA398233 | SYSY |
| SRR5931735 | HepG2-rep2 | SINGLE | input | PRJNA398233 | SYSY |
| SRR5931736 | HepG2-rep2 | SINGLE | ip | PRJNA398233 | SYSY |
| SRR7992454 | NHDF-rep2 | SINGLE | ip | PRJNA451188 | NEB |
| SRR7992455 | NHDF-rep1 | SINGLE | ip | PRJNA451188 | NEB |
| SRR7992458 | NHDF-rep1 | SINGLE | input | PRJNA451188 | NEB |
| SRR7992459 | NHDF-rep3 | SINGLE | ip | PRJNA451188 | NEB |
| SRR7992460 | NHDF-rep3 | SINGLE | input | PRJNA451188 | NEB |
| SRR7992461 | NHDF-rep2 | SINGLE | input | PRJNA451188 | NEB |
| SRR4310472 | HEK293T-rep1 | SINGLE | input | PRJNA344999 | SYSY |
| SRR4310473 | HEK293T-rep2 | SINGLE | input | PRJNA34499 | SYSY |

| | | | | | |
|---|---|---|---|---|---|
| | | | | 9 | |
| SRR4310474 | HEK293T-rep3 | SINGLE | input | PRJNA344999 | SYSY |
| SRR4310478 | HEK293T-rep1 | SINGLE | ip | PRJNA344999 | SYSY |
| SRR4310479 | HEK293T-rep2 | SINGLE | ip | PRJNA344999 | SYSY |
| SRR4310480 | HEK293T-rep3 | SINGLE | ip | PRJNA344999 | SYSY |
| SRR4042257 | Jurkat-rep1 | SINGLE | input | PRJNA339176 | SYSY |
| SRR4042258 | Jurkat-rep2 | SINGLE | input | PRJNA339176 | SYSY |
| SRR4042263 | Jurkat-rep1 | SINGLE | ip | PRJNA339176 | SYSY |
| SRR4042264 | Jurkat-rep2 | SINGLE | ip | PRJNA339176 | SYSY |
| SRR903368 | U2OS-rep1 | SINGLE | input | PRJNA208737 | SYSY |
| SRR903369 | U2OS-rep2 | SINGLE | input | PRJNA208737 | SYSY |
| SRR903370 | U2OS-rep3 | SINGLE | input | PRJNA208737 | SYSY |
| SRR903374 | U2OS-rep1 | SINGLE | ip | PRJNA208737 | SYSY |
| SRR903375 | U2OS-rep2 | SINGLE | ip | PRJNA208737 | SYSY |
| SRR903376 | U2OS-rep3 | SINGLE | ip | PRJNA208737 | SYSY |
| SRR456551 | HepG2-rep1 | SINGLE | ip | PRJNA158111 | SYSY |
| SRR456552 | HepG2-rep2 | SINGLE | ip | PRJNA158111 | SYSY |
| SRR456553 | HepG2-rep3 | SINGLE | ip | PRJNA158111 | SYSY |
| SRR456555 | HepG2-rep1 | SINGLE | input | PRJNA158111 | SYSY |
| SRR456556 | HepG2-rep2 | SINGLE | input | PRJNA158111 | SYSY |
| SRR456557 | HepG2-rep3 | SINGLE | input | PRJNA158111 | SYSY |
| SRR5925271 | HeLa-rep1 | PAIRED | input | PRJNA397879 | SYSY |
| SRR5925272 | HeLa-rep2 | PAIRED | input | PRJNA397879 | SYSY |
| SRR5925273 | HeLa-rep1 | PAIRED | ip | PRJNA397879 | SYSY |
| SRR5925274 | HeLa-rep2 | PAIRED | ip | PRJNA397879 | SYSY |
| SRR6686554 | HepG2-rep1 | SINGLE | input | PRJNA43347 | SYSY |

| | | | | | |
|---|---|---|---|---|---|
| | | | | 2 | |
| SRR6686555 | HepG2-rep2 | SINGLE | input | PRJNA433472 | SYSY |
| SRR6686556 | HepG2-rep3 | SINGLE | input | PRJNA433472 | SYSY |
| SRR6686557 | HepG2-rep1 | SINGLE | ip | PRJNA433472 | SYSY |
| SRR6686558 | HepG2-rep2 | SINGLE | ip | PRJNA433472 | SYSY |
| SRR6686559 | HepG2-rep3 | SINGLE | ip | PRJNA433472 | SYSY |
| SRR6954135 | HeLa-rep1 | SINGLE | input | PRJNA448996 | SYSY |
| SRR6954136 | HeLa-rep2 | SINGLE | input | PRJNA448996 | SYSY |
| SRR6954137 | HeLa-rep1 | SINGLE | ip | PRJNA448996 | SYSY |
| SRR6954138 | HeLa-rep2 | SINGLE | ip | PRJNA448996 | SYSY |
| SRR14765588 | HEK293T-rep1 | PAIRED | input | PRJNA634388 | SYSY |
| SRR14765589 | HEK293T-rep1 | PAIRED | ip | PRJNA634388 | SYSY |
| SRR14765590 | HEK293T-rep2 | PAIRED | input | PRJNA634388 | SYSY |
| SRR14765591 | HEK293T-rep2 | PAIRED | ip | PRJNA634388 | SYSY |
| 2020082301 | hESC-rep1 | PAIRED | input | PRJNA665206 | NEB |
| 2020082303 | hESC-rep1 | PAIRED | ip | PRJNA665206 | NEB |
| 2020082305 | hESC-rep2 | PAIRED | input | PRJNA665206 | NEB |
| 2020082307 | hESC-rep2 | PAIRED | ip | PRJNA665206 | NEB |
| CRR072994 | HeLa-rep1 | PAIRED | input | PRJCA001180 | Millipore |
| CRR072995 | HeLa-rep1 | PAIRED | ip | PRJCA001180 | Millipore |
| CRR072996 | HeLa-rep2 | PAIRED | input | PRJCA001180 | Millipore |
| CRR072997 | HeLa-rep2 | PAIRED | ip | PRJCA001180 | Millipore |
| CRR073001 | K562-rep1 | PAIRED | ip | PRJCA00118 | Millipore |

| | | | | | |
|---|---|---|---|---|---|
| | | | | 0 | |
| CRR073003 | K562-rep2 | PAIRED | ip | PRJCA001180 | Millipore |
| CRR073000 | K562-rep1 | PAIRED | input | PRJCA001180 | Millipore |
| CRR073002 | K562-rep2 | PAIRED | input | PRJCA001180 | Millipore |
| CRR073007 | WPMY-rep1 | PAIRED | ip | PRJCA001180 | Millipore |
| CRR073009 | WPMY-rep2 | PAIRED | ip | PRJCA001180 | Millipore |
| CRR073006 | WPMY-rep1 | PAIRED | input | PRJCA001180 | Millipore |
| CRR073008 | WPMY-rep2 | PAIRED | input | PRJCA001180 | Millipore |
| CRR073013 | Jurkat-rep1 | PAIRED | ip | PRJCA001180 | Millipore |
| CRR073015 | Jurkat-rep2 | PAIRED | ip | PRJCA001180 | Millipore |
| CRR073012 | Jurkat-rep1 | PAIRED | input | PRJCA001180 | Millipore |
| CRR073014 | Jurkat-rep2 | PAIRED | input | PRJCA001180 | Millipore |
| CRR042274 | HEK293T-rep1 | PAIRED | ip | PRJCA001180 | Millipore |
| CRR042276 | HEK293T-rep2 | PAIRED | ip | PRJCA001180 | Millipore |
| CRR055565 | HEK293T-rep3 | PAIRED | ip | PRJCA001180 | Millipore |
| CRR042275 | HEK293T-rep1 | PAIRED | input | PRJCA001180 | Millipore |
| CRR042277 | HEK293T-rep2 | PAIRED | input | PRJCA001180 | Millipore |
| CRR055566 | HEK293T-rep3 | PAIRED | input | PRJCA001180 | Millipore |

Supplementary Table 2 m6A-seq data information for different human tissues

| Run accession | Sample title | Library layout | Sample type | Study accession | Antibody info |
|---|---|---|---|---|---|
| CRR042290 | Heart-4-2 | PAIRED | ip | PRJCA001180 | Millipore |
| CRR042291 | Heart-4-2 | PAIRED | input | PRJCA001180 | Millipore |
| CRR055527 | Heart-1-1 | PAIRED | ip | PRJCA001180 | Millipore |
| CRR055528 | Heart-1-1 | PAIRED | input | PRJCA001180 | Millipore |
| CRR042302 | Skin-4-2 | PAIRED | ip | PRJCA001180 | Millipore |

| | | | | | |
|---|---|---|---|---|---|
| CRR042303 | Skin-4-2 | PAIRED | input | PRJCA001180 | Millipore |
| CRR042304 | Skin-1-1 | PAIRED | ip | PRJCA001180 | Millipore |
| CRR042305 | Skin-1-1 | PAIRED | input | PRJCA001180 | Millipore |
| CRR042284 | Cerebellum-5-3 | PAIRED | ip | PRJCA001180 | Millipore |
| CRR073016 | Cerebellum-7-4 | PAIRED | ip | PRJCA001180 | Millipore |
| CRR042285 | Cerebellum-5-3 | PAIRED | input | PRJCA001180 | Millipore |
| CRR073017 | Cerebellum-7-4 | PAIRED | input | PRJCA001180 | Millipore |
| CRR042286 | Cerebrum-5-3 | PAIRED | ip | PRJCA001180 | Millipore |
| CRR055553 | Cerebrum-6-3 | PAIRED | ip | PRJCA001180 | Millipore |
| CRR042287 | Cerebrum-5-3 | PAIRED | input | PRJCA001180 | Millipore |
| CRR055554 | Cerebrum-6-3 | PAIRED | input | PRJCA001180 | Millipore |
| CRR042312 | Thyroid-gland-4-2 | PAIRED | ip | PRJCA001180 | Millipore |
| CRR042314 | Thyroid-gland-5-3 | PAIRED | ip | PRJCA001180 | Millipore |
| CRR042313 | Thyroid-gland-4-2 | PAIRED | input | PRJCA001180 | Millipore |
| CRR042315 | Thyroid-gland-5-3 | PAIRED | input | PRJCA001180 | Millipore |
| CRR042306 | Stomach-4-2 | PAIRED | ip | PRJCA001180 | Millipore |
| CRR042308 | Stomach-5-3 | PAIRED | ip | PRJCA001180 | Millipore |
| CRR042307 | Stomach-4-2 | PAIRED | input | PRJCA001180 | Millipore |
| CRR042309 | Stomach-5-3 | PAIRED | input | PRJCA001180 | Millipore |
| CRR055551 | Jejunum-4-2 | PAIRED | ip | PRJCA001180 | Millipore |
| CRR055555 | Jejunum-5-3 | PAIRED | ip | PRJCA001180 | Millipore |
| CRR055552 | Jejunum-4-2 | PAIRED | input | PRJCA001180 | Millipore |
| CRR055556 | Jejunum-5-3 | PAIRED | input | PRJCA001180 | Millipore |
| CRR055543 | Appendix-3-2 | PAIRED | ip | PRJCA001180 | Millipore |
| CRR055557 | Appendix-5-3 | PAIRED | ip | PRJCA001180 | Millipore |
| CRR055544 | Appendix-3-2 | PAIRED | input | PRJCA001180 | Millipore |
| CRR055558 | Appendix-5-3 | PAIRED | input | PRJCA001180 | Millipore |
| CRR055559 | Rectum-5-3 | PAIRED | ip | PRJCA001180 | Millipore |
| CRR055563 | Rectum-4-2 | PAIRED | ip | PRJCA001180 | Millipore |
| CRR055560 | Rectum-5-3 | PAIRED | input | PRJCA001180 | Millipore |
| CRR055564 | Rectum-4-2 | PAIRED | input | PRJCA001180 | Millipore |
| CRR042280 | Aorta-4-2 | PAIRED | ip | PRJCA001180 | Millipore |
| CRR055531 | Aorta-1-1 | PAIRED | ip | PRJCA001180 | Millipore |
| CRR042281 | Aorta-4-2 | PAIRED | input | PRJCA001180 | Millipore |
| CRR055532 | Aorta-1-1 | PAIRED | input | PRJCA001180 | Millipore |
| CRR055547 | Esophagus-3-2 | PAIRED | ip | PRJCA001180 | Millipore |
| CRR055561 | Esophagus-4-2 | PAIRED | ip | PRJCA001180 | Millipore |
| CRR055548 | Esophagus-3-2 | PAIRED | input | PRJCA001180 | Millipore |
| CRR055562 | Esophagus-4-2 | PAIRED | input | PRJCA001180 | Millipore |
| CRR055529 | Spleen-1-1 | PAIRED | ip | PRJCA001180 | Millipore |
| CRR055536 | Spleen-2-1 | PAIRED | ip | PRJCA001180 | Millipore |
| CRR055541 | Spleen-3-2 | PAIRED | ip | PRJCA001180 | Millipore |
| CRR055530 | Spleen-1-1 | PAIRED | input | PRJCA001180 | Millipore |
| CRR055535 | Spleen-2-1 | PAIRED | input | PRJCA001180 | Millipore |

| | | | | | |
|---|---|---|---|---|---|
| CRR055542 | Spleen-3-2 | PAIRED | input | PRJCA001180 | Millipore |
| CRR042318 | Urinary-bladder-4-2 | PAIRED | ip | PRJCA001180 | Millipore |
| CRR042320 | Urinary-bladder-5-3 | PAIRED | ip | PRJCA001180 | Millipore |
| CRR055539 | Urinary-bladder-2-1 | PAIRED | ip | PRJCA001180 | Millipore |
| CRR042319 | Urinary-bladder-4-2 | PAIRED | input | PRJCA001180 | Millipore |
| CRR042321 | Urinary-bladder-5-3 | PAIRED | input | PRJCA001180 | Millipore |
| CRR055540 | Urinary-bladder-2-1 | PAIRED | input | PRJCA001180 | Millipore |
| CRR042296 | Lung-4-2 | PAIRED | ip | PRJCA001180 | Millipore |
| CRR055533 | Lung-2-1 | PAIRED | ip | PRJCA001180 | Millipore |
| CRR073018 | Lung-2-4 | PAIRED | ip | PRJCA001180 | Millipore |
| CRR073020 | Lung-4-4 | PAIRED | ip | PRJCA001180 | Millipore |
| CRR042297 | Lung-4-2 | PAIRED | input | PRJCA001180 | Millipore |
| CRR055534 | Lung-2-1 | PAIRED | input | PRJCA001180 | Millipore |
| CRR073019 | Lung-2-4 | PAIRED | input | PRJCA001180 | Millipore |
| CRR073021 | Lung-4-4 | PAIRED | input | PRJCA001180 | Millipore |
| SRR8209837 | Cerebellum-1 | PAIRED | ip | PRJNA506210 | SYSY |
| SRR8209839 | Cerebellum-2 | PAIRED | ip | PRJNA506210 | SYSY |
| SRR8209841 | Cerebellum-3 | PAIRED | ip | PRJNA506210 | SYSY |
| SRR8209836 | Cerebellum-1 | PAIRED | input | PRJNA506210 | SYSY |
| SRR8209838 | Cerebellum-2 | PAIRED | input | PRJNA506210 | SYSY |
| SRR8209840 | Cerebellum-3 | PAIRED | input | PRJNA506210 | SYSY |
| SRR8209843 | FrontalCortex-1 | PAIRED | ip | PRJNA506210 | SYSY |
| SRR8209845 | FrontalCortex-2 | PAIRED | ip | PRJNA506210 | SYSY |
| SRR8209847 | FrontalCortex-3 | PAIRED | ip | PRJNA506210 | SYSY |
| SRR8209842 | FrontalCortex-1 | PAIRED | input | PRJNA506210 | SYSY |
| SRR8209844 | FrontalCortex-2 | PAIRED | input | PRJNA506210 | SYSY |
| SRR8209846 | FrontalCortex-3 | PAIRED | input | PRJNA506210 | SYSY |
| SRR8209849 | Heart-1 | PAIRED | ip | PRJNA506210 | SYSY |
| SRR8209853 | Heart-2 | PAIRED | ip | PRJNA506210 | SYSY |
| SRR820985 | Heart-3 | PAIRED | ip | PRJNA50621 | SYSY |

| | | | | | |
|---|---|---|---|---|---|
| 5 | | | | 0 | |
| SRR8209848 | Heart-1 | PAIRED | input | PRJNA506210 | SYSY |
| SRR8209850 | Heart-2 | PAIRED | input | PRJNA506210 | SYSY |
| SRR8209854 | Heart-3 | PAIRED | input | PRJNA506210 | SYSY |
| SRR8209857 | Kidney-1 | PAIRED | ip | PRJNA506210 | SYSY |
| SRR8209859 | Kidney-2 | PAIRED | ip | PRJNA506210 | SYSY |
| SRR8209861 | Kidney-3 | PAIRED | ip | PRJNA506210 | SYSY |
| SRR8209856 | Kidney-1 | PAIRED | input | PRJNA506210 | SYSY |
| SRR8209858 | Kidney-2 | PAIRED | input | PRJNA506210 | SYSY |
| SRR8209860 | Kidney-3 | PAIRED | input | PRJNA506210 | SYSY |
| SRR8209863 | Liver-1 | PAIRED | ip | PRJNA506210 | SYSY |
| SRR8209865 | Liver-2 | PAIRED | ip | PRJNA506210 | SYSY |
| SRR8209867 | Liver-3 | PAIRED | ip | PRJNA506210 | SYSY |
| SRR8209862 | Liver-1 | PAIRED | input | PRJNA506210 | SYSY |
| SRR8209864 | Liver-2 | PAIRED | input | PRJNA506210 | SYSY |
| SRR8209866 | Liver-3 | PAIRED | input | PRJNA506210 | SYSY |
| SRR8209869 | Lung-1 | PAIRED | ip | PRJNA506210 | SYSY |
| SRR8209871 | Lung-2 | PAIRED | ip | PRJNA506210 | SYSY |
| SRR8209873 | Lung-3 | PAIRED | ip | PRJNA506210 | SYSY |
| SRR8209868 | Lung-1 | PAIRED | input | PRJNA506210 | SYSY |
| SRR8209870 | Lung-2 | PAIRED | input | PRJNA506210 | SYSY |
| SRR8209872 | Lung-3 | PAIRED | input | PRJNA506210 | SYSY |
| SRR820987 | Muscle-1 | PAIRED | ip | PRJNA50621 | SYSY |

| | | | | | |
|---|---|---|---|---|---|
| 5 | | | | 0 | |
| SRR8209877 | Muscle-2 | PAIRED | ip | PRJNA506210 | SYSY |
| SRR8209879 | Muscle-3 | PAIRED | ip | PRJNA506210 | SYSY |
| SRR8209874 | Muscle-1 | PAIRED | input | PRJNA506210 | SYSY |
| SRR8209876 | Muscle-2 | PAIRED | input | PRJNA506210 | SYSY |
| SRR8209878 | Muscle-3 | PAIRED | input | PRJNA506210 | SYSY |
| SRR8209881 | Spleen-1 | PAIRED | ip | PRJNA506210 | SYSY |
| SRR8209883 | Spleen-2 | PAIRED | ip | PRJNA506210 | SYSY |
| SRR8209885 | Spleen-3 | PAIRED | ip | PRJNA506210 | SYSY |
| SRR8209880 | Spleen-1 | PAIRED | input | PRJNA506210 | SYSY |
| SRR8209882 | Spleen-2 | PAIRED | input | PRJNA506210 | SYSY |
| SRR8209884 | Spleen-3 | PAIRED | input | PRJNA506210 | SYSY |
| SRR7130857 | heart-1 | SINGLE | ip | PRJNA464886 | Abcam |
| SRR7130858 | heart-2 | SINGLE | ip | PRJNA464886 | Abcam |
| SRR7130859 | heart-3 | SINGLE | ip | PRJNA464886 | Abcam |
| SRR7130879 | heart-1 | PAIRED | input | PRJNA464886 | Abcam |
| SRR7130880 | heart-2 | PAIRED | input | PRJNA464886 | Abcam |
| SRR7130881 | heart-3 | PAIRED | input | PRJNA464886 | Abcam |
| SRR7130860 | kidney-2 | SINGLE | ip | PRJNA464886 | Abcam |
| SRR7130861 | kidney-3 | SINGLE | ip | PRJNA464886 | Abcam |
| SRR7130862 | kidney-4 | SINGLE | ip | PRJNA464886 | Abcam |
| SRR7130882 | kidney-2 | PAIRED | input | PRJNA464886 | Abcam |
| SRR713088 | kidney-3 | PAIRED | input | PRJNA46488 | Abcam |

| | | | | | |
|---|---|---|---|---|---|
| 3 | | | | 6 | |
| SRR7130884 | kidney-4 | PAIRED | input | PRJNA464886 | Abcam |
| SRR7130863 | liver-1 | SINGLE | ip | PRJNA464886 | Abcam |
| SRR7130864 | liver-2 | SINGLE | ip | PRJNA464886 | Abcam |
| SRR7130865 | liver-3 | SINGLE | ip | PRJNA464886 | Abcam |
| SRR7130885 | liver-1 | PAIRED | input | PRJNA464886 | Abcam |
| SRR7130886 | liver-2 | PAIRED | input | PRJNA464886 | Abcam |
| SRR7130887 | liver-3 | PAIRED | input | PRJNA464886 | Abcam |
| SRR7130872 | placenta-2 | SINGLE | ip | PRJNA464886 | Abcam |
| SRR7130873 | placenta-4 | SINGLE | ip | PRJNA464886 | Abcam |
| SRR7130874 | placenta-6 | SINGLE | ip | PRJNA464886 | Abcam |
| SRR7130894 | placenta-2 | PAIRED | input | PRJNA464886 | Abcam |
| SRR7130895 | placenta-4 | PAIRED | input | PRJNA464886 | Abcam |
| SRR7130896 | placenta-6 | PAIRED | input | PRJNA464886 | Abcam |
| SRR7130866 | lung-4 | SINGLE | ip | PRJNA464886 | Abcam |
| SRR7130867 | lung-5 | SINGLE | ip | PRJNA464886 | Abcam |
| SRR7130888 | lung-4 | PAIRED | input | PRJNA464886 | Abcam |
| SRR7130889 | lung-5 | PAIRED | input | PRJNA464886 | Abcam |
| SRR7130868 | muscle-4 | SINGLE | ip | PRJNA464886 | Abcam |
| SRR7130869 | muscle-5 | SINGLE | ip | PRJNA464886 | Abcam |
| SRR7130890 | muscle-4 | PAIRED | input | PRJNA464886 | Abcam |
| SRR7130891 | muscle-5 | PAIRED | input | PRJNA464886 | Abcam |
| SRR713087 | stomach-4 | SINGLE | ip | PRJNA46488 | Abcam |

| | | | | | |
|---|---|---|---|---|---|
| 0 | | | | 6 | |
| SRR7130871 | stomach-5 | SINGLE | ip | PRJNA464886 | Abcam |
| SRR7130892 | stomach-4 | PAIRED | input | PRJNA464886 | Abcam |
| SRR7130893 | stomach-5 | PAIRED | input | PRJNA464886 | Abcam |
| SRR7130854 | brain-1 | SINGLE | ip | PRJNA464886 | Abcam |
| SRR7130855 | brain-2 | SINGLE | ip | PRJNA464886 | Abcam |
| SRR7130856 | brain-3 | SINGLE | ip | PRJNA464886 | Abcam |
| SRR7130875 | brain-1 | PAIRED | input | PRJNA464886 | Abcam |
| SRR7130876 | brain-2 | PAIRED | input | PRJNA464886 | Abcam |
| SRR7130877 | brain-3 | PAIRED | input | PRJNA464886 | Abcam |
| SRR7130878 | brain-4 | PAIRED | input | PRJNA464886 | Abcam |

Supplementary Table 3 m6A-seq data information for different mouse cell lines

| run accession | sample title | library layout | sample type | study accession | antibody info |
|---|---|---|---|---|---|
| SRR6144434 | NPC-rep1 | SINGLE | input | PRJNA413499 | SYSY |
| SRR6144435 | NPC-rep2 | SINGLE | input | PRJNA413499 | SYSY |
| SRR6144436 | NPC-rep1 | SINGLE | ip | PRJNA413499 | SYSY |
| SRR6144437 | NPC-rep2 | SINGLE | ip | PRJNA413499 | SYSY |
| SRR1596085 | mESC-rep1 | SINGLE | ip | PRJNA262906 | SYSY |
| SRR1596086 | mESC-rep1 | SINGLE | input | PRJNA262906 | SYSY |
| SRR1596087 | mESC-rep2 | SINGLE | ip | PRJNA262906 | SYSY |
| SRR1596088 | mESC-rep2 | SINGLE | input | PRJNA262906 | SYSY |
| SRR1596089 | mESC-rep3 | SINGLE | ip | PRJNA262906 | SYSY |
| SRR1596090 | mESC-rep3 | SINGLE | input | PRJNA262906 | SYSY |
| SRR1596097 | MEF-rep1 | SINGLE | ip | PRJNA262906 | SYSY |
| SRR1596098 | MEF-rep1 | SINGLE | input | PRJNA262906 | SYSY |
| SRR1596099 | MEF-rep2 | SINGLE | ip | PRJNA262906 | SYSY |
| SRR1596100 | MEF-rep2 | SINGLE | input | PRJNA262906 | SYSY |
| SRR1048180 | 3T3-L1-rep1 | SINGLE | input | PRJNA231244 | SYSY |
| SRR1048181 | 3T3-L1-rep1 | SINGLE | ip | PRJNA231244 | SYSY |
| SRR1575981 | 3T3-L1-rep2 | SINGLE | input | PRJNA231244 | SYSY |
| SRR1575982 | 3T3-L1-rep2 | SINGLE | ip | PRJNA231244 | SYSY |
| SRR6163656 | NSPC-rep1 | SINGLE | input | PRJNA414058 | NEB |
| SRR6163657 | NSPC-rep2 | SINGLE | input | PRJNA414058 | NEB |

| | | | | | |
|---|---|---|---|---|---|
| SRR6163658 | NSPC-rep3 | SINGLE | input | PRJNA414058 | NEB |
| SRR6163662 | NSPC-rep1 | SINGLE | ip | PRJNA414058 | NEB |
| SRR6163663 | NSPC-rep2 | SINGLE | ip | PRJNA414058 | NEB |
| SRR6163664 | NSPC-rep3 | SINGLE | ip | PRJNA414058 | NEB |
| SRR7235655 | Dendritic-cells-rep1 | SINGLE | input | PRJNA473808 | NEB |
| SRR7235656 | Dendritic-cells-rep2 | SINGLE | input | PRJNA473808 | NEB |
| SRR7235659 | Dendritic-cells-rep1 | SINGLE | ip | PRJNA473808 | NEB |
| SRR7235660 | Dendritic-cells-rep2 | SINGLE | ip | PRJNA473808 | NEB |
| SRR8130792 | Dendritic-cells-rep1 | PAIRED | ip | PRJNA499089 | NEB |
| SRR8130793 | Dendritic-cells-rep2 | PAIRED | ip | PRJNA499089 | NEB |
| SRR8130796 | Dendritic-cells-rep1 | PAIRED | input | PRJNA499089 | NEB |
| SRR8130797 | Dendritic-cells-rep2 | PAIRED | input | PRJNA499089 | NEB |
| SRR14765644 | mESC-rep1 | PAIRED | input | PRJNA634388 | NEB |
| SRR14765645 | mESC-rep1 | PAIRED | ip | PRJNA634388 | NEB |
| SRR14765646 | mESC-rep2 | PAIRED | input | PRJNA634388 | NEB |
| SRR14765647 | mESC-rep2 | PAIRED | ip | PRJNA634388 | NEB |

Supplementary Table 4 m6A-seq data information for different mouse tissues

| run accession | sample title | library layout | sample type | study accession | antibody info |
|---|---|---|---|---|---|
| SRR496283 | Brain-1 | SINGLE | ip | PRJNA141085 | SYSY |
| SRR496284 | Brain-1 | SINGLE | input | PRJNA141085 | SYSY |
| SRR496285 | Brain-2 | SINGLE | input | PRJNA141085 | SYSY |
| SRR496287 | Brain-2 | SINGLE | ip | PRJNA141085 | SYSY |
| SRR1596091 | Embryoid-bodies-1 | SINGLE | ip | PRJNA262906 | SYSY |
| SRR1596092 | Embryoid-bodies-1 | SINGLE | input | PRJNA262906 | SYSY |
| SRR1596093 | Embryoid-bodies-2 | SINGLE | ip | PRJNA262906 | SYSY |
| SRR1596094 | Embryoid-bodies-2 | SINGLE | input | PRJNA262906 | SYSY |
| SRR1596096 | Embryoid-bodies-3 | SINGLE | input | PRJNA262906 | SYSY |
| SRR1745915 | Embryoid-bodies-3 | SINGLE | ip | PRJNA262906 | SYSY |
| SRR5468919 | Testis-1 | SINGLE | ip | PRJNA383941 | SYSY |
| SRR5468920 | Testis-2 | SINGLE | ip | PRJNA383941 | SYSY |
| SRR5468921 | Testis-1 | SINGLE | input | PRJNA383941 | SYSY |
| SRR5468922 | Testis-2 | SINGLE | input | PRJNA383941 | SYSY |
| SRR866997 | Midbrain-1 | SINGLE | ip | PRJNA205151 | SYSY |
| SRR866998 | Midbrain-1 | SINGLE | input | PRJNA205151 | SYSY |
| SRR866999 | Midbrain-2 | SINGLE | ip | PRJNA205151 | SYSY |
| SRR867000 | Midbrain-2 | SINGLE | input | PRJNA205151 | SYSY |
| SRR867001 | Midbrain-3 | SINGLE | ip | PRJNA205151 | SYSY |
| SRR867002 | Midbrain-3 | SINGLE | input | PRJNA205151 | SYSY |
| SRR520481 | Hippocampus | SINGL | input | PRJNA36884 | SYSY |

| | | | | | |
|---|---|---|---|---|---|
| 1 | -2-weeks-1 | E | | 9 | |
| SRR5204812 | Hippocampus-2-weeks-2 | SINGLE | input | PRJNA368849 | SYSY |
| SRR5204813 | Hippocampus-2-weeks-1 | SINGLE | ip | PRJNA368849 | SYSY |
| SRR5204814 | Hippocampus-2-weeks-2 | SINGLE | ip | PRJNA368849 | SYSY |
| SRR5204815 | Hippocampus-6-weeks-1 | SINGLE | input | PRJNA368849 | SYSY |
| SRR5204816 | Hippocampus-6-weeks-2 | SINGLE | input | PRJNA368849 | SYSY |
| SRR5204817 | Hippocampus-6-weeks-1 | SINGLE | ip | PRJNA368849 | SYSY |
| SRR5204818 | Hippocampus-6-weeks-2 | SINGLE | ip | PRJNA368849 | SYSY |
| CRR055567 | Heart-1-5 | PAIRED | input | PRJCA001180 | Millipore |
| CRR055568 | Heart-1-5 | PAIRED | ip | PRJCA001180 | Millipore |
| CRR072984 | Heart-1-4 | PAIRED | input | PRJCA001180 | Millipore |
| CRR072985 | Heart-1-4 | PAIRED | ip | PRJCA001180 | Millipore |
| CRR055583 | Heart-2-5 | PAIRED | input | PRJCA001180 | Millipore |
| CRR055584 | Heart-2-5 | PAIRED | ip | PRJCA001180 | Millipore |
| CRR055569 | Spleen-1-5 | PAIRED | input | PRJCA001180 | Millipore |
| CRR055570 | Spleen-1-5 | PAIRED | ip | PRJCA001180 | Millipore |
| CRR055585 | Spleen-2-5 | PAIRED | input | PRJCA001180 | Millipore |
| CRR055586 | Spleen-2-5 | PAIRED | ip | PRJCA001180 | Millipore |
| CRR055573 | Liver-1-5 | PAIRED | input | PRJCA001180 | Millipore |
| CRR055574 | Liver-1-5 | PAIRED | ip | PRJCA001180 | Millipore |
| CRR072982 | Liver-1-4 | PAIRED | input | PRJCA001180 | Millipore |
| CRR072983 | Liver-1-4 | PAIRED | ip | PRJCA001180 | Millipore |
| CRR055589 | Liver-2-5 | PAIRE | input | PRJCA00118 | Millipor |

| | | | | | |
|---|---|---|---|---|---|
| | | D | | 0 | e |
| CRR055590 | Liver-2-5 | PAIRED | ip | PRJCA001180 | Millipore |
| CRR055593 | Cerebellum-2-5 | PAIRED | input | PRJCA001180 | Millipore |
| CRR055594 | Cerebellum-2-5 | PAIRED | ip | PRJCA001180 | Millipore |
| CRR055577 | Cerebellum-1-5 | PAIRED | input | PRJCA001180 | Millipore |
| CRR055578 | Cerebellum-1-5 | PAIRED | ip | PRJCA001180 | Millipore |
| CRR055581 | Hypothalamus-1-5 | PAIRED | input | PRJCA001180 | Millipore |
| CRR055582 | Hypothalamus-1-5 | PAIRED | ip | PRJCA001180 | Millipore |
| CRR055597 | Hypothalamus-2-5 | PAIRED | input | PRJCA001180 | Millipore |
| CRR055598 | Hypothalamus-2-5 | PAIRED | ip | PRJCA001180 | Millipore |
| CRR055575 | Cerebrum-1-5 | PAIRED | input | PRJCA001180 | Millipore |
| CRR055576 | Cerebrum-1-5 | PAIRED | ip | PRJCA001180 | Millipore |
| CRR055591 | Cerebrum-2-5 | PAIRED | input | PRJCA001180 | Millipore |
| CRR055592 | Cerebrum-2-5 | PAIRED | ip | PRJCA001180 | Millipore |
| CRR055579 | Brainstem-1-5 | PAIRED | input | PRJCA001180 | Millipore |
| CRR055580 | Brainstem-1-5 | PAIRED | ip | PRJCA001180 | Millipore |
| CRR055595 | Brainstem-2-5 | PAIRED | input | PRJCA001180 | Millipore |
| CRR055596 | Brainstem-2-5 | PAIRED | ip | PRJCA001180 | Millipore |
| CRR055587 | Lung-2-5 | PAIRED | input | PRJCA001180 | Millipore |
| CRR055588 | Lung-2-5 | PAIRED | ip | PRJCA001180 | Millipore |
| CRR055571 | Lung-1-5 | PAIRED | input | PRJCA001180 | Millipore |
| CRR055572 | Lung-1-5 | PAIRED | ip | PRJCA001180 | Millipore |
| CRR072986 | Lung-1-4 | PAIRE | input | PRJCA00118 | Millipor |

| | | | | | |
|---|---|---|---|---|---|
| | | D | | 0 | e |
| CRR072987 | Lung-1-4 | PAIRED | ip | PRJCA001180 | Millipore |
| CRR072988 | Lung-2-4 | PAIRED | input | PRJCA001180 | Millipore |
| CRR072989 | Lung-2-4 | PAIRED | ip | PRJCA001180 | Millipore |

Supplementary Table 5 ChIP-seq data information for HeLa and mESC

| run accession | sample title | library layout | sample type | study accession |
|---|---|---|---|---|
| SRR7130897 | HeLa-METTL3 | SINGLE | input | PRJNA464886 |
| SRR7130898 | HeLa-METTL3 | SINGLE | ip | PRJNA464886 |
| SRR8545657 | mESC-METTL3 | PAIRED | input | PRJNA521368 |
| SRR12187085 | mESC-METTL3-rep1 | PAIRED | ip | PRJNA521368 |
| SRR12187193 | mESC-METTL3-rep2 | PAIRED | ip | PRJNA521368 |
